# Patterns of pan-genome occupancy and gene co-expression under water-deficit in *Brachypodium distachyon*

**DOI:** 10.1101/2021.07.21.453242

**Authors:** Rubén Sancho, Pilar Catalán, Bruno Contreras-Moreira, Thomas E. Juenger, David L. Des Marais

## Abstract

Natural populations are characterized by abundant genetic diversity driven by a range of different types of mutation. The tractability of sequencing complete genomes has allowed new insights into the variable composition of genomes, summarized as a species pan-genome. These analyses demonstrate that many genes are absent from the first reference genomes, whose analysis dominated the initial years of the genomic era. Our field now turns towards understanding the functional consequence of these highly variable genomes. Here, we analyzed weighted gene co-expression networks from leaf transcriptome data for drought response in the purple false brome *Brachypodium distachyon* and the differential expression of genes putatively involved in adaptation to this stressor. We specifically asked whether genes with variable “occupancy” in the pan-genome – genes which are either present in all studied genotypes or missing in some genotypes – show different distributions among co-expression modules. Co-expression analysis united genes expressed in drought-stressed plants into nine modules covering 72 hub genes (87 hub isoforms), and genes expressed under controlled water conditions into 13 modules, covering 190 hub genes (251 hub isoforms). We find that low occupancy pan-genes are under-represented among several modules, while other modules are over-enriched for low-occupancy pan-genes. We also provide new insight into the regulation of drought response in *B. distachyon*, specifically identifying one module with an apparent role in primary metabolism that is strongly responsive to drought. Our work shows the power of integrating pan-genomic analysis with transcriptomic data using factorial experiments to understand the functional genomics of environmental response.

## INTRODUCTION

Soil water availability is a critical factor determining plant growth, development, and reproduction (Bohnert, Nelson, & Jensen, 1995). Plants are able to cope with and acclimate to a range of soil water contents through the reprogramming of their physiology, growth, and development over time scales ranging from hours to seasons (Chaves, Maroco, & Pereira, 2003). Many of these acclimation strategies arise from altered transcriptional profiles (Fisher et al., 2016; Miao, Han et al., 2017). Drought-responsive gene regulatory pathways have been investigated extensively in model plant systems such as *Arabidopsis thaliana*, maize, and rice (Borah et al., 2017; Hayano-Kanashiro et al., 2009; Janiak, Kwa, & Szarejko, 2015; Nakashima, Ito, & Yamaguchi-Shinozaki, 2009; Nakashima, Yamaguchi-Shinozaki, & Shinozaki, 2014). A clear emerging theme, however, is that diverse species and varieties of plants exhibit diverse stress response mechanisms (Des Marais et al., 2012; Juenger, 2013; Pinheiro & Chaves, 2011), often controlled by complex regulatory networks. Understanding the genetic control of this phenotypic diversity is a priority for understanding the response of natural populations to climate change, and for designing resilient crop species (Benfey & Mitchell-Olds, 2008).

Recent studies have brought attention to the remarkable variation in gene content among plant populations (Alonge et al., 2020; Gao et al., 2019; Gordon et al., 2017; Haberer et al., 2020), reflected in a species’ pan-genome. A pan-genome refers to the genomic content of a species as a whole, rather than the composition of a single, reference, individual genotype (Koonin & Wolf, 2008). In practice, pan-genomes are estimated by deeply resequencing the genomes of a diversity panel of genotypes, often using a reference genome to aid in final assembly and annotation (Lei et al., 2021). In the diploid model grass *Brachypodium distachyon*, genomic analysis of 56 inbred natural “accessions” revealed that the total pan-genome of the species comprised nearly twice the number of genes in any single accession (Gordon et al., 2017). Remarkably, only 73% of genes in a given accession are found in at least 95% of the other accessions (Gordon et al., 2017) – so-called “core genes” (universal or nearly universal genes; Koonin & Wolf, 2008) and “soft-core genes” (found in at least 95% of accessions; Kaas et al. 2012) – suggesting that a large number of genes are unique to subsets of accessions or even to individual accessions. The list of core genes in *B. distachyon* is enriched for annotations associated with essential cellular processes such as primary metabolism. Lower-occupancy genes, or “shell genes,” are found in 5-94% of accessions and their annotations are enriched for many processes related to environmental response, including disease resistance. Similar patterns have been observed in the pan-genomes of *Arabidopsis thaliana*, barley, sunflower, and an ever-growing number of additional plant species (Bayer et al., 2020; Contreras-Moreira et al., 2017; Hübner et al., 2019). The DNA sequences of core genes bear the hallmark of strong purifying selection and are typically expressed at a higher level and in more tissues as compared to shell genes. Shell genes may be the result of gene duplications or deletions in the ancestor of a subset of studied genotypes and, indeed, the vast majority of shell genes in *B. distachyon* appear to be functional, as homologs are found in other species’ genomes (Gordon et al., 2017).

The preceding observations raise the intriguing possibility that shell genes may represent segregating variation that could be shaped by natural selection and thereby facilitate local adaptation or adaptive responses to a variable environment. Multiple studies in *Arabidopsis thaliana* demonstrate the role of segregating functional gene copies – effectively large-effect mutations – in shaping whole-plant response to the abiotic environment (Monroe et al., 2016, 2018). The phenotypic effect size of a mutation can determine the likelihood that the mutant will become fixed in a population, with large-effect mutations more likely than not to confer deleterious phenotypes that may be removed from populations by natural selection (Fisher, 1930). The observation that any two accessions of *B. distachyon* likely differ in the presence or absence of hundreds of functional gene copies begs the question as to how potentially function-changing gene deletions escape the purging effects of purifying selection. Pan-genomics requires that we reconceptualize how we interpret “gene loss” as we move beyond a reference-genome view of genome function. Genes identified as “missing” from a subset of accessions might represent deletions of genes whose function was no longer constrained by purifying selection in some novel environment, duplicated genes that originated in a subset of accessions and thus are absent in other accessions, or paralogs that were both present in a common ancestor and then reciprocally lost in subsets of accessions.

The pleiotropic effect of a mutation can be affected by the number of genes with which it interacts (Jeong et al., 2001); if a gene has relatively few interacting partners then its presence or absence in a particular accession may have a small fitness effect and thus be maintained in populations. Similarly, the efficacy of selection to purge deleterious alleles may be reduced if a gene is only expressed in a subset of environments experienced by a species (Paaby & Rockman, 2014). In the context of pan-genomes, we consider gene presence/absence polymorphisms as mutations, and we explore functional gene turnover by testing the hypothesis that shell genes and core genes differ in their topological positions in environmentally responsive gene co-expression networks.

Gene co-expression networks are widely used to interpret functional genomic data by assessing patterns of correlation among genes via a threshold that assigns a connection weight to each gene pair (Langfelder & Horvath, 2008; Zhang & Horvath, 2005). Sets of genes with similar expression profiles are assigned to modules by applying graph clustering algorithms (Mao et al., 2009). Genes, or “nodes,” in such networks show considerable variation in the extent to which their expression co-varies with other genes. As such, co-expression networks are generally considered “scale-free” in the sense that few nodes have many neighboring nodes while many nodes have few neighboring nodes (Guelzim et al., 2002).

Modules are often comprised of genes with similar functions (Stuart et al., 2003; Wolfe, Kohane, & Butte, 2005). High connectivity “hub” genes that show a large number of interactions with other genes within a weighted co-expression network are candidates for key players in regulating cellular processes (Albert, Jeong, & Barabási, 2000; Carlson et al., 2006; Dong & Horvath, 2007). As such, hub genes might be expected to encode essential cellular functions and thus show pleiotropic effects when mutated or deleted. By contrast, genes with fewer close co-expression relationships are often situated on the periphery of networks and might, therefore, exhibit fewer pleiotropic effects when missing or mutated (Des Marais et al., 2017a; Masalia, Bewick, & Burke, 2017; Porth et al., 2014). In this context, we hypothesize that pan-genome core genes may be over-represented among co-expression network “hub-genes,” as both appear to be involved in core cellular processes and may therefore show deleterious effects when deleted. Conversely, we hypothesize that pan-genome “shell genes” – whose patterns of expression and thereby phenotypic effects are more restricted and condition-specific – will be enriched among lowly connected (non-hub) genes in gene co-expression networks.

Here, we study the relationship between a plant’s pan-genome and its gene co-expression networks using *Brachypodium distachyon. Brachypodium* is a small genus of the subfamily Pooideae (Poaceae) that contains ∼20 species distributed worldwide (Catalán et al., 2016; Scholthof et al., 2018; Hasterok et al. 2022). The annual diploid species *B. distachyon* is a model for temperate cereals and biofuel grasses (Vogel et al. 2010; Mur et al. 2011; Catalán et al. 2014; Scholthof et al. 2018); a reference genome for one *B. distachyon* accession, Bd21 (IBI, 2010) is now complemented by 54 deeply resequenced natural accessions (Gordon et al., 2017). Recent studies demonstrate the utility of *B. distachyon* and its close congeners for elucidating the evolution and ecology of plant-abiotic interactions, focusing especially on responses to soil drying, aridity, and water use strategy (Des Marais & Juenger, 2016; Des Marais et al., 2017b; Monroe et al. 2021; Fisher et al., 2016; Handakumbura et al., 2019; Manzaneda et al., 2015, 2012; Martínez et al., 2018; Skalska et al., 2020; Verelst et al., 2013). In the present study, we first identify and characterize gene co-expression modules associated with response to soil drying. We then test the hypothesis that the occupancy of pan-genes – whether they are part of the shell or core gene sets of the pan-genome – is associated with their connectivity in the *B. distachyon* gene co-expression network. Our work demonstrates the dynamic nature of plant genomes and sets up future work on the functional consequences of diversity on the evolution of gene regulatory networks.

## MATERIALS AND METHODS

### Plant material, experimental design, total-RNA extraction and 3’ cDNA tag libraries preparation

Sampling herein follows our earlier work documenting physiological and developmental response of 33 diploid natural accessions of *Brachypodium distachyon* (L.) P. Beauv. to soil drying (Figure 1; Table S1) (Des Marais et al., 2017b). The sampled accessions were inbred for more than five generations (Filiz et al., 2009; Vogel et al., 2006, 2009) and represent the geographic and ecological diversity of *B. distachyon* across the Mediterranean region. Whole genome resequencing data is available for all studied accessions (Gordon et al., 2017).

**Figure 1.**
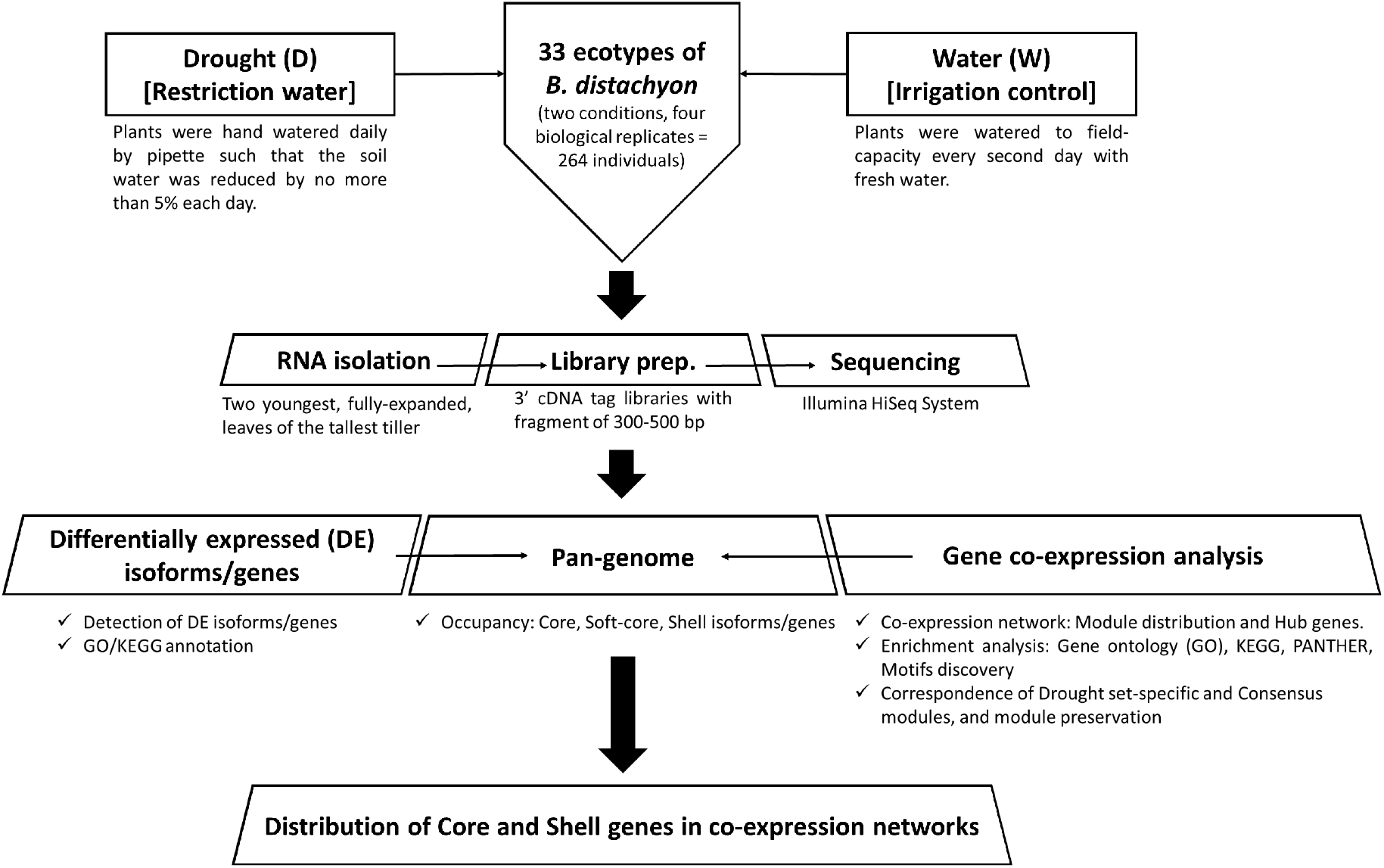
Summary of the experimental design and analyses performed in the 33 accessions of the model grass *Brachypodium distachyon* under drought (D) and water (control) (W) conditions.

A total of 264 individual plants from the 33 accessions were grown under two greenhouse conditions, restriction of water (drought, D) and well-watered (water, W). We sampled four biological replicates per treatment and accession combinations [33 accessions x 4 replicates x 2 treatments (D and W)]. For a full description of the growth and treatment conditions, please see Des Marais, Lasky, et al. (2017b). In short, seeds were stratified at 6°C for 14d and then greenhouse-grown in 600 mL of Profile porous ceramic rooting media (Profile Products) in Deepot D40H pots (650 mL; Stuewe & Sons). For the first 21d of growth, all plants were watered to field capacity every other day. Daytime high temperatures ranged from 23°C to 28°C and night-time lows from 14°C to 18°C. On day 21 four treatment regimes were implemented: Cool Wet, Cool Drought, Hot Wet, and Hot Drought, with Drought and Wet plants spatially randomized within single blocks of Hot or Cool conditions. The hot treatment raised air temperatures in the plant canopies by ∼10°C. Because temperature was confounded with experimental block in the design used for the current study, we did not include a temperature effect in any of the statistical models used herein. Well-watered plants (hereafter “Water”) were watered to field-capacity every second day with fresh water, whereas drought plants (hereafter “Drought”) were hand-watered daily by pipette such that the soil water was reduced by 5% each day. The final soil water content of Drought plants on day 11 of treatment – 32d post-germination – was 45% field capacity, which corresponds to a decrease in water potential of 1.2 MPa as compared to field capacity in this growth media (Des Marais et al., 2012).

For each plant, the two youngest, fully expanded leaves of the tallest tiller were excised with a razor blade at the base of the lamina and flash-frozen on liquid nitrogen. Tissue was ground to a fine powder under liquid nitrogen using a Mixer Mill MM 300 (Retsch GmbH). RNA was extracted using the Sigma Spectrum Total Plant RNA kit, including on-column DNase treatment, following the manufacturer’s protocol, and quantified using a NanoDrop spectrophotometer (Thermo Scientific). We used a RNA-Seq library protocol (3’ cDNA tag libraries with fragment of 300-500 bp) for sequencing on the Illumina HiSeq platform adapted from Meyer et al. (2011). This Tag-Seq method yields only one sequence per expressed transcript in the RNA pool, allowing for higher sequencing coverage per gene as a function of total sequencing effort (Tandonnet & Torres, 2017).

### Pre-processing of sequences, quantifying transcript abundances, normalizing procedures, and controlling batch effects

Sequencing was carried out using an Illumina HiSeq2500 platform (100 bp Single-End, SE, sequencing). Quality control of SE reads was performed with FastQC software. Adapters and low quality reads were removed and filtered with Trimmomatic-0.32 (Bolger, Lohse, & Usadel, 2014). Total numbers of raw and filtered SE reads for each accession and treatment are shown in Table S2. Quantifying the abundances of transcripts from RNA-Seq data was done with Kallisto v0.43.1 (Bray et al., 2016). To accommodate the library preparation and sequencing protocols (3’ tag from fragments of 300-500 bp), pseudoalignments of RNA-Seq data were carried out using as references 500 bp from the 3’ tails of the *B. distachyon*_314_v3.1 transcriptome (IBI 2010; https://phytozome.jgi.doe.gov/pz/portal.html#!info?alias=Org_Bdistachyon). We applied estimated average fragment lengths of 100 bp and standard deviations of fragment length of 20. Resulting numbers of transcripts per million (TPM) were recorded.

Exploratory analysis of the data set and the subsequent filtering and normalization of transcripts abundance between samples, and the *in silico* technical replicate steps (bootstrap values computed with Kallisto), were conducted with the Sleuth package (Pimentel et al., 2017). A total of 16,386 targets (transcripts/isoforms) were recovered after the normalizing and filtering step using Sleuth. This program was also used for batch-correction of data and of differentially expressed (DE) genes. To account for library preparation batch effects, date of library preparation was included as a covariate with condition variable in the full model (Table S2).

### Co-expression network analysis of normalized transcripts abundance

Co-expression networks for the Drought and Water (control) data sets were carried out using the transcripts per million (TPM) estimates and the R package WGCNA (Langfelder and Horvath 2008). We analyzed 16,386 transcripts that were filtered and normalized for 127 Drought and 124 Water individual samples (individual plants). After the removal of putative outliers, we retained 121 Drought and 108 Water samples that were used for network construction.

Identical parameters were used for the Drought and the Water data sets to construct their respective co-expression networks. The *BlockwiseModules* function was used to perform automatic network construction and module detection on the large expression data set of transcripts. Parameters for co-expression network construction were fitted checking different values. We chose the Pearson correlation and unsigned network type, soft thresholding power 6 (high scale free, R^2^>0.85), a minimum module size of 30, and a medium sensitivity (deepSplit = 2) for the cluster splitting. The topological overlap matrix (TOM) was generated using the TOMtype unsigned approach. Module clustering was performed with function *cutreeDynamic* and the Partitioning Around Medoids (PAM) option activated. Module merging was conducted with mergeCutHeight set to 0.30. K_diff_ was calculated using WGCNA to estimate the relationship between connectivity among genes within *vs* between co-expression modules.

Both isoform and gene counts were calculated. Isoform counts included all transcripts identified (e.g., Bradi1g1234.1; Bradi1g1234.2; Bradi1g1234.3) and gene counts only included different genes expressed, thus different isoforms from the same gene were reduced to a single gene count in each case (e.g., Bradi1g12345.1 and Bradi1g12345.2 are two isoforms of one gene, Bradi1g12345).

### Detection of highly connected nodes (hub genes/ isoforms) within co-expression networks

Three representative descriptors of modules, module eigengene (ME), intramodular connectivity (k_IM_), and eigengene-based connectivity (k_ME_; or its equivalent module membership, MM) were calculated using the WGCNA package. Briefly, ME is defined as the first principal component of a given module and is often considered to represent the gene expression profiles within the module. k_IM_ measures how connected, or co-expressed, a given gene is with respect to the genes of a particular module. Thus, intra-modular connectivity is also the connectivity in the subnetwork defined by the module. MM is the correlation of gene expression profile with the module eigengene (ME) of a given module. MM values close to 1 or −1 indicate genes highly connected to the module. The sign of MM indicates a positive or a negative relationship between a gene and the eigengene of the module (Langfelder & Horvath, 2010). Genes with absolute MM value over 0.9 were considered “hub genes.” Correlations between MM values transformed by a power of β = 6 and k_IM_ values were also calculated.

### Pan-genome occupancy of clustered, hub, and differentially expressed genes across accessions

Because the *B. distachyon* accessions studied herein comprise a subset of those included in the original pan-genome (Gordon et al. 2017), we re-ran the clustering procedures used in our earlier analysis with only the 33 accessions used here. We clustered CDS sequences from the annotated genomes of each of the studied accessions to define core, soft-core, and shell genes with the software GET_HOMOLOGUES-EST v03012018 (Contreras-Moreira et al., 2017) using the OMCL algorithm (-M) and a stringent percent-sequence identity threshold (–S 98). The resulting pan-genome matrix was interrogated to identify “core” genes observed in all 33 accessions, “soft-core” genes observed in 32 and 31, and “shell” genes observed in 30 or fewer accessions. Occupancy was defined as the number of accessions that contain a particular gene model. We tested whether each module showed an excess or deficit of shell genes as compared to genome averages of pan-gene occupancy using a Fisher’s Exact Test, as implemented in the R programming language (R Core Team, 2022).

### Enrichment and GO/KEGG annotation of clustered genes

Gene ontology (GO) and the Kyoto Encyclopedia of Genes and Genomes (KEGG) annotations for the *B. distachyon* 314 (Bd21 acccession) v.3.1 reference genome were retrieved (http://phytozome.jgi.doe.gov/; IBI 2010). Gene lists were tested for functional enrichments with the PANTHER (Protein ANalysis THrough Evolutionary Relationships) overrepresentation test (http://www.pantherdb.org). The original *B. distachyon* 314 (Bd21 accession) v.3.1 gene ids were converted to v.3.0 with help from Ensembl Plants (Howe et al., 2020) to match those in PANTHER16.0 (Mi et al., 2021). Tests were conducted on all genes and on both conditions -- Drought and Water -- applying the Fisher’s Exact test with False Discovery Rate (FDR) multiple test correction. This analysis was applied on different data sets: all genes, and pan-genome core, soft-core, and shell genes for each co-expressed module.

### Drought versus Watered modular structure preservation and comparison between consensus and set-specific modules

Permutation tests were performed to check for preservation of the module topology in the Drought (discovery data) and the Water (test data) networks using both the approach of Langfelder et al. (Langfelder, Luo, Oldham, & Horvath, 2011) as well as the modulePreservation function of the NetRep (Ritchie et al. 2016) R package with null=”all” (include all nodes) option for RNA-Seq data. All NetRep test statistics (Module coherence, Average node contribution, Concordance of node contributions, Density of correlation structure, Concordance of correlation structure, Average edge weight, and Concordance of weighted degree) were evaluated through permutation analysis; therefore, a module was considered preserved if all the statistics had a permutation test P-value < 0.01. Searching for modules that could play a role in drought response, we focused on Drought modules that were unpreserved in the Water network (P-value > 0.01 in at least 1 of the seven statistics presented in Ritchie et al. (2016). The Consensus modules (Cons) and the respective relationships between Consensus and Drought (D) and Water (W) modules were performed as described in Langfelder and Horvath (2008).

### Annotations of upstream DNA motifs in the co-expression modules

Genes assigned to modules in the Drought and Water networks were further analyzed with the objective of discovering DNA motifs putatively involved in their expression in each module. Motif analysis was carried out using a protocol based on RSAT::Plants (Ksouri et al., 2021). This approach allowed us to discover DNA motifs enriched in the promoter regions of co-expressed genes and to match them to curated signatures of experimentally described transcription factors. First, −500 bp (upstream) to +200 bp (downstream) sequences around the transcription start sites of the genes detected in each module and 50 random negative controls from other gene promoter regions of equal size were extracted from the *B. distachyon* Bd21 v3.0 (Ensembl Plants version 46) reference genome. Then, the peak-motifs (Thomas-Chollier et al., 2012) option was used to discover enriched motifs applying a 2^nd^ order Markov genomic model, and GO enrichment was computed for them. The analyses generated a report with links to similar curated motifs in the database footprintDB as scored with normalized correlation (Ncor) (Sebastian and Contreras-Moreira 2014). For each module a highly supported DNA motif was selected according to Ncor, e-value, and the number of sites (i.e., putative cis-regulatory elements, CREs) used to compile the motif. The matrix-scan tool (Turatsinze et al., 2008), with a weight threshold set to 70% of the motif length (60% for the small module D8), was used to scan the discovered motifs and to identify individual genes within each module harboring putative CREs.

All the protein sequences predicted for each co-expression module were analyzed using iTAK (Plant Transcription factor & Protein Kinase Identifier and Classifier) online (v1.6) (Zheng et al., 2016) to annotate their respective transcription factors, transcriptional regulators and related protein kinases (classification system by Lehti-Shiu and Shiu (2012)).

### Differentially expressed (DE) isoforms/genes

In order to determine how many isoforms and genes were differentially expressed between the two treatments (D vs W), the two data sets were analyzed through the sleuth_result function (Pimentel et al., 2017). This function computes likelihood ratio tests (LRT) for null (no treatment effect) and alternative (treatment effect) hypotheses, attending to the full and reduced fitted models. A significant q-value ≤ 1E-6 threshold was fixed to detect DE isoforms. The 50 most differentially expressed genes (25 up-regulated and 25 down-regulated) were classified based on fold-change of the average TPM values between the drought and water treatments.

### Data availability

The *Brachypodium distachyon* filtered RNA-Seq data were deposited in the ENA (European Nucleotide Archive; https://www.ebi.ac.uk/ena) with the consecutive accession numbers from ERR6133302 to ERR6133575. All the R scripts, supplementary Excel files, TPM counts and adjacency matrices are available in the Github repository identified by the following doi: xxxxxxxxxx created using Zenodo.

## RESULTS

Our RNA-Seq dataset comprised 121 Drought and 108 Water samples, obtained from 33 inbred accessions of *B. distachyon* (Figure 1). After the filtering and quality control steps, we identified 16,386 transcripts in our analyses comprising 4,941 isoforms of 3,789 differentially expressed (DE) genes between the water and drought treatments (Supplementary files S1; S2). Of these genes, 2,808 were upregulated (3,591 isoforms) and 980 downregulated (1,350 isoforms) in D vs W conditions. One gene (Bradi1g09950) showed both up-regulated (Bradi1g09950.2) and down-regulated (Bradi1g09950.1) isoforms under drought conditions (Supplementary file S2).

### Modular distribution of genes in the gene co-expression networks

The 33 genetically diverse accessions, along with random experimental variance, provided a wealth of expression variance that was leveraged to estimate major axes of variation – co-expression modules – from the RNA-Seq data. The modular distribution of genes in the Drought and Water gene co-expression networks showed differences both in the number and the size of the modules (Figures S1a,b and S2a,b; Tables 1a,b and S3a,b). The Drought co-expression network comprised 9 modules (D1-D9) containing a total of 5,020 isoforms (min = 38, max = 2,477 isoforms per module), corresponding to 4,006 genes (min = 27, max = 1,986 genes per module). The largest D1 module contained 15.1% of the isoforms (16.1% of the genes) whereas two modules (D2, D3) clustered 4-6% of the isoforms and genes each and the remaining modules clustered ≤ 2% of isoforms and genes each (Fig. S1a; Table 1a). 11,366 isoforms (69.4%; 8,313 genes, 67.4%) were not clustered in any Drought module using our criteria (gray or “zero” D0 module) (Figs. S1a and S2a; Table 1a). The Water co-expression network showed 13 modules (W1-W13) containing a total of 6,711 isoforms (min = 48, max = 1,866 isoforms per module), corresponding to 5,407 genes (min = 40, max = 1,439 genes per module). The largest W1 contained 11.4% of the isoforms (11.6% of the genes) whereas six modules (W2-W7) clustered over >2-6% of the isoforms and genes and the remaining six modules ≤ 2% each (Fig. S1b; Table 1b). 9,675 isoforms (59.0%; 6,934 genes, 56.0%) were not clustered in any Water module (gray or “zero” W0 module; Figs. S1b and S2b; Table 1b). Hereafter, co-expression modules identified in the Drought RNA-Seq dataset will be labeled with the prefix “D” while those modules identified in the Water (control) dataset will have the “W” prefix.

**Table 1.** Statistics of the number and percentage of isoforms (all and hub isoforms) and genes (all and hub genes), and mean k_diff_ (the difference between intra- and inter-modular connectivity) per module for each Drought **(a)** and Water **(b)** co-expression networks. The quantity of total counts of genes in drought and water networks is different because we counted the genes for each module independently. When several isoforms of the same gene were clustered in different modules, the gene was counted multiple times; however, it was counted only once if it clustered in the same module. For example, if D1 has three isoforms (Bradi1g10000.1, Bradi1g10000.2 and Bradi1g10000.3) of the same gene, it was counted only once (Bradi1g10000); when isoforms were found in different modules (e.g., Bradi1g10000.1 in D1, Bradi1g10000.2 in D2, Bradi1g10000.3 in D3), the gene was counted three times.

| Drought modules |  | isoforms (nodes) per module |  | genes per module |  | mean Kdiff |
| --- | --- | --- | --- | --- | --- | --- |
|  |  | all | hub<br>(KME>0.9) | all | hub<br>(KME>0.9) |  |
| D0 | Gray | 11366 (69.4%) | nd | 8313 (67.4%) | nd | nd |
| D1 | Turquoise | 2477 (15.1%) | 4 (0.16%) | 1986 (16.1%) | 3 (0.15%) | 7.9 |
| D2 | Blue | 979 (6.0%) | 20 (2.04%) | 750 (6.1%) | 13 (1.73%) | 3.7 |
| D3 | Brown | 750 (4.6%) | 41 (5.47%) | 627 (5.1%) | 38 (6.06%) | 10.1 |
| D4 | Yellow | 318 (1.9%) | 4 (1.26%) | 258 (2.1%) | 4 (1.55%) | 3.7 |
| D5 | Green | 214 (1.3%) | 4 (1.87%) | 169 (1.4%) | 3 (1.78%) | 0.8 |
| D6 | Red | 111 (0.7%) | 10 (9.01%) | 95 (0.8%) | 7 (7.37%) | -1.5 |
| D7 | Black | 68 (0.4%) | 4 (5.88%) | 48 (0.4%) | 4 (8.3%) | -0.2 |
| D8 | Pink | 65 (0.4%) | 0 (0%) | 57 (0.5%) | 0 (0%) | -5.9 |
| D9 | Magenta | 38 (0.2%) | 0 (0%) | 27 (0.2%) | 0 (0%) | -8.8 |
| Total counts |  | 16386 | 87 (0.53%) | 12330 | 72 (0.58%) |  |
| Total unique counts |  | 16386 | 87 (0.53%) | 12137 | 72 (0.59%) |  |
| Total unique counts<br>(excluding gray module) |  | 5020 | 87 (1.73%) | 4006 | 72 (1.80%) |  |

(b)
| Water modules |  | isoforms (nodes) per module |  | genes per module |  | mean<br>Kdiff |
| --- | --- | --- | --- | --- | --- | --- |
|  |  | all | hub<br>(KME>0.9) | all | hub<br>(KME>0.9) |  |
| W0 | Gray | 9675 (59.0%) | nd | 6934 (56.0%) | nd | nd |
| W1 | Turquoise | 1866 (11.4%) | 149 (7.98%) | 1439 (11.6%) | 115 (7.99%) | 26.8 |
| W2 | Blue | 870 (5.3%) | 4 (0.46%) | 727 (5.9%) | 3 (0.41%) | -3.7 |
| W3 | Brown | 827 (5.0%) | 2 (0.24%) | 696 (5.6%) | 1 (0.14%) | 0.3 |
| W4 | Yellow | 701 (4.3%) | 2 (0.29%) | 590 (4.8%) | 2 (0.34%) | -0.7 |
| W5 | Green | 607 (3.7%) | 12 (1.98%) | 509 (4.1%) | 10 (1.96%) | 3.1 |
| W6 | Red | 435 (2.7%) | 9 (2.07%) | 358 (2.9%) | 8 (2.08%) | 2.1 |
| W7 | Black | 390 (2.4%) | 26 (6.67%) | 313 (2.5%) | 23 (7.35%) | 4.6 |
| W8 | Pink | 288 (1.7%) | 2 (0.69%) | 245 (2.0%) | 2 (0.82%) | -2.2 |
| W9 | Magenta | 282 (1.7%) | 22 (7.80) | 239 (1.9%) | 21 (8.79%) | 5.8 |
| W10 | Purple | 246 (1.5%) | 5 (2.03%) | 185 (1.5%) | 2 (1.08%) | -1.4 |
| W11 | Greenyellow | 89 (0.5%) | 8 (8.99%) | 55 (0.4%) | 1 (1.81%) | -3.1 |
| W12 | Tan | 62 (0.4%) | 1 (1.61%) | 51 (0.4%) | 1 (1.96%) | -0.9 |
| W13 | Salmon | 48 (0.3%) | 9 (18.75%) | 40 (0.3%) | 1 (2.5%) | -1.6 |
| Total counts |  | 16386 | 251 (1.53%) | 12381 | 190 (1.53%) |  |
| Total unique counts |  | 16386 | 251 (1.53%) | 12137 | 190 (1.57%) |  |
| Total unique counts<br>(excluding gray<br>module) |  | 6711 | 251 (3.74%) | 5407 | 190 (3.51%) |  |

While the genes comprising co-expression modules likely represent functionally related genes, modules are not discrete entities and many genes within a module are likely co-expressed with genes in other modules. The largest Drought modules showed a positive (D1, D2, D3, D4, and D5) or slightly negative (D6 and D7) mean k_diff_, while the smallest modules had more negative values (D8 and D9) (Table 1a). Negative k_diff_ values for a node indicate that connectivity out of the module is higher than intra-modular connectivity. Similarly, the Water modules showed positive mean k_diff_ values in the largest modules (W1, W3, W5, W6, W7, and W9) or slightly negative (W4), with the exception of one large and one intermediate modules (W2 and W8) both having negative values similar to those of the smallest modules (Table 1b). High positive linear correlations between MM values and k_IM_ values were recovered in both Drought (Fig. S3a) and Water (Fig. S3b) networks, thus validating the criterion of high MM (>0.9) for selecting hub genes. Collectively, these results suggest that modules of both Drought and Water networks were statistically well-supported but also that many of their genes likely share transcriptional activity with genes included in other modules.

### Preservation and correspondence of Drought and Water networks

We tested the hypothesis that some co-expression modules are only observed under one of our two treatment conditions using two approaches. First, a permutation test was performed using NetRep (Ritchie et al., 2016) to test for the preservation of module topology in the Drought versus the Water networks, defined as discovery and test data sets, respectively. This test comprises seven topological statistics on each module and condition (drought vs water), quantifying the replication of the structural relationship between nodes composing each module under the null hypothesis that the module of interest is not preserved. All Drought and Water modules were topologically preserved according to the seven NetRep statistics (permutation test P-values < 0.01). We further tested for correspondence between Drought (D) and Water (W) set-specific and Drought-Water Consensus (Cons) co-expression modules using WGCNA (Fig. 2a,b). Consensus modules are those shared by two or more networks (Langfelder & Horvath, 2007). We found that several Drought specific modules were comprised of no or very few isoforms included in the Consensus modules (Fig. 2a). Thus, the Drought modules D5 (green), D7 (black) and D9 (magenta) did not show a clear correspondence with consensus modules. The seemingly conflicting results between NetRep – which detected module preservation between Drought and Water modules – and WGCNA – which identified three modules that were not preserved – reflected the different sensitivities of these two approaches to changing co-expression relationships within modules. This suggested that all modules were preserved in a broad sense between treatments, but that nodes within modules D5, D7, and D9 could have slightly different co-expression relationships than those observed in the corresponding modules in the Water treatment. We also found Water specific modules (Fig. 2b), W1 (turquoise), W6 (red), W8 (pink), and W12 (tan), that did not overlap with the Consensus modules or overlapped only with the non-co-expressed gray module. W1 and W6 overlapped with the gray consensus module, whereas the smaller modules W8 and W12 did not overlap with any consensus module.

**Figure 2.**
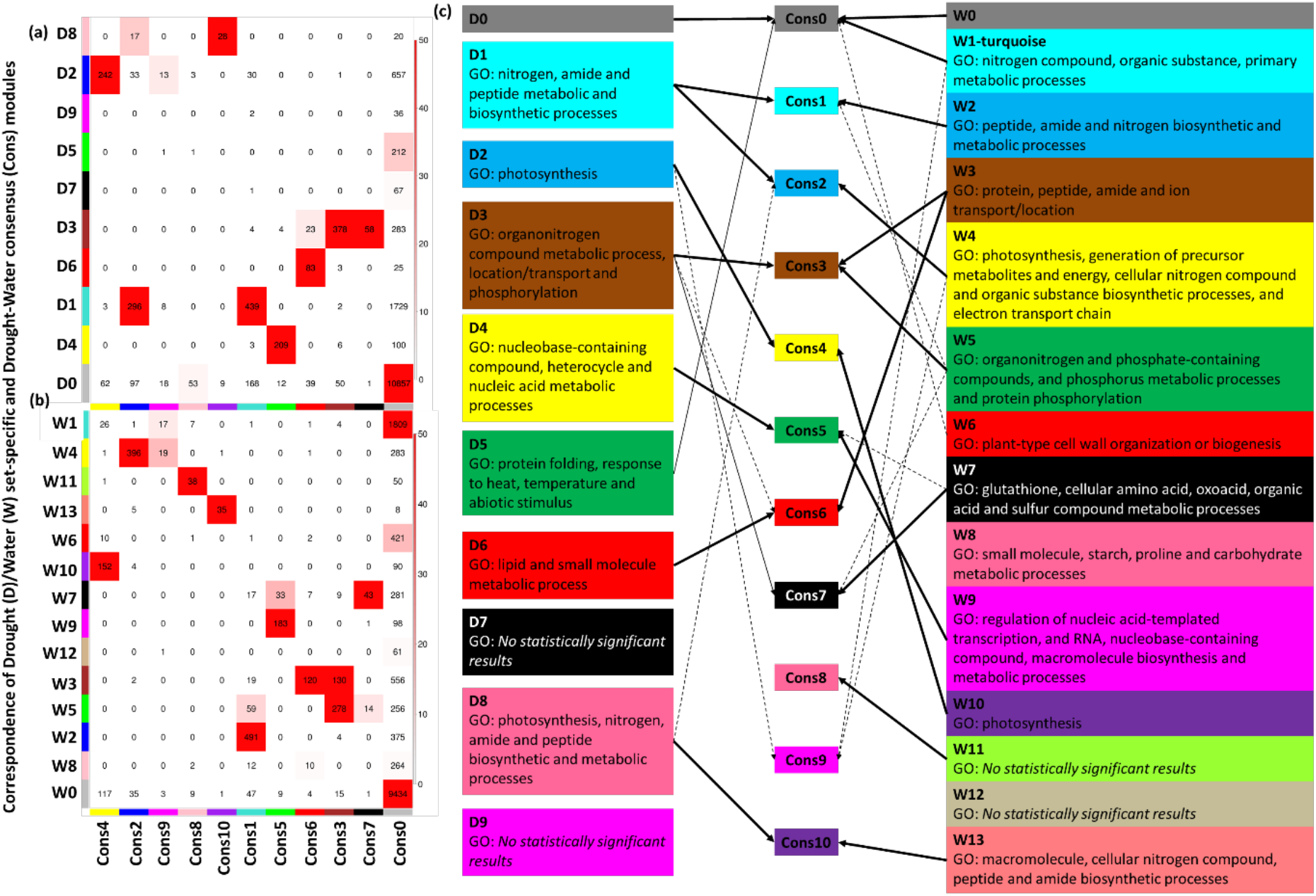
Correspondence (number of nodes) of Drought (D) (**a**) and Water (W) (**b**) set-specific and Drought-Water consensus (Cons) modules. Each row of the table corresponds to one Drought/Water set-specific module, and each column corresponds to one consensus module. Numbers in the table indicate node counts in the intersection of the corresponding modules. Coloring of the table encodes − log(p), with p being the Fisher’s exact test p-value for the overlap of the two modules. The stronger the red color, the more significant the overlap is. (**c**) Comparison of the GO enrichments for each D and W module, according to the correlation with the common consensus module.

Seven modules in the Drought network and eleven in the Water network showed a significant GO term enrichment (Table 2a,b; Supplementary file S3). Both networks shared modules enriched for different biological processes. Potentially equivalent modules between the D and W networks were inferred attending to the common consensus module with which they overlapped (Figure 2a,b) and their GO enrichments (Fig. 2c). Thus, the D1 and W2 modules matched Cons1 and were GO-enriched in nitrogen, amide, and peptide metabolic and biosynthetic processes, D2 and W10 matched Cons4 and were enriched in the photosynthesis, D3 and W3 matched Cons3 and were enriched in processes of transport and locations of compounds, D4 and W9 were enriched in nucleic acid metabolic processes, and D8 and W13 in nitrogen, amide, and peptide biosynthetic and metabolic processes. However, some modules showed enrichment in biological process unique to one or the other co-expression network. For example, D5 is enriched in genes involved in protein folding, response to heat, temperature, and abiotic stimulus, and W6 in genes predicted to be involved in cell wall organization or biogenesis. Collectively, these co-expression analyses suggest that the detected modules represent groups of functionally related genes, and that many functional relationships among genes were conserved in the Water and Drought networks.

**Table 2.**
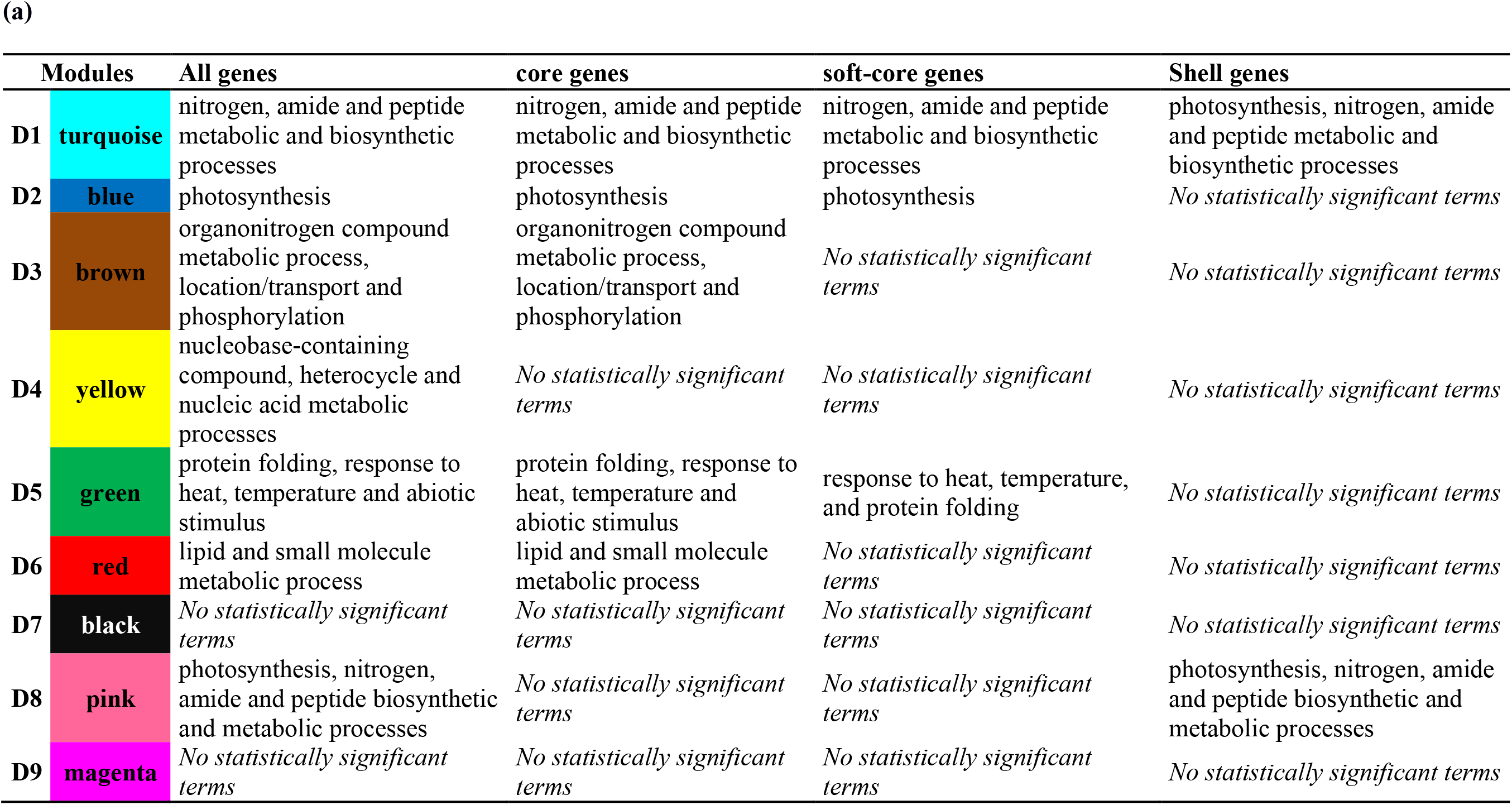
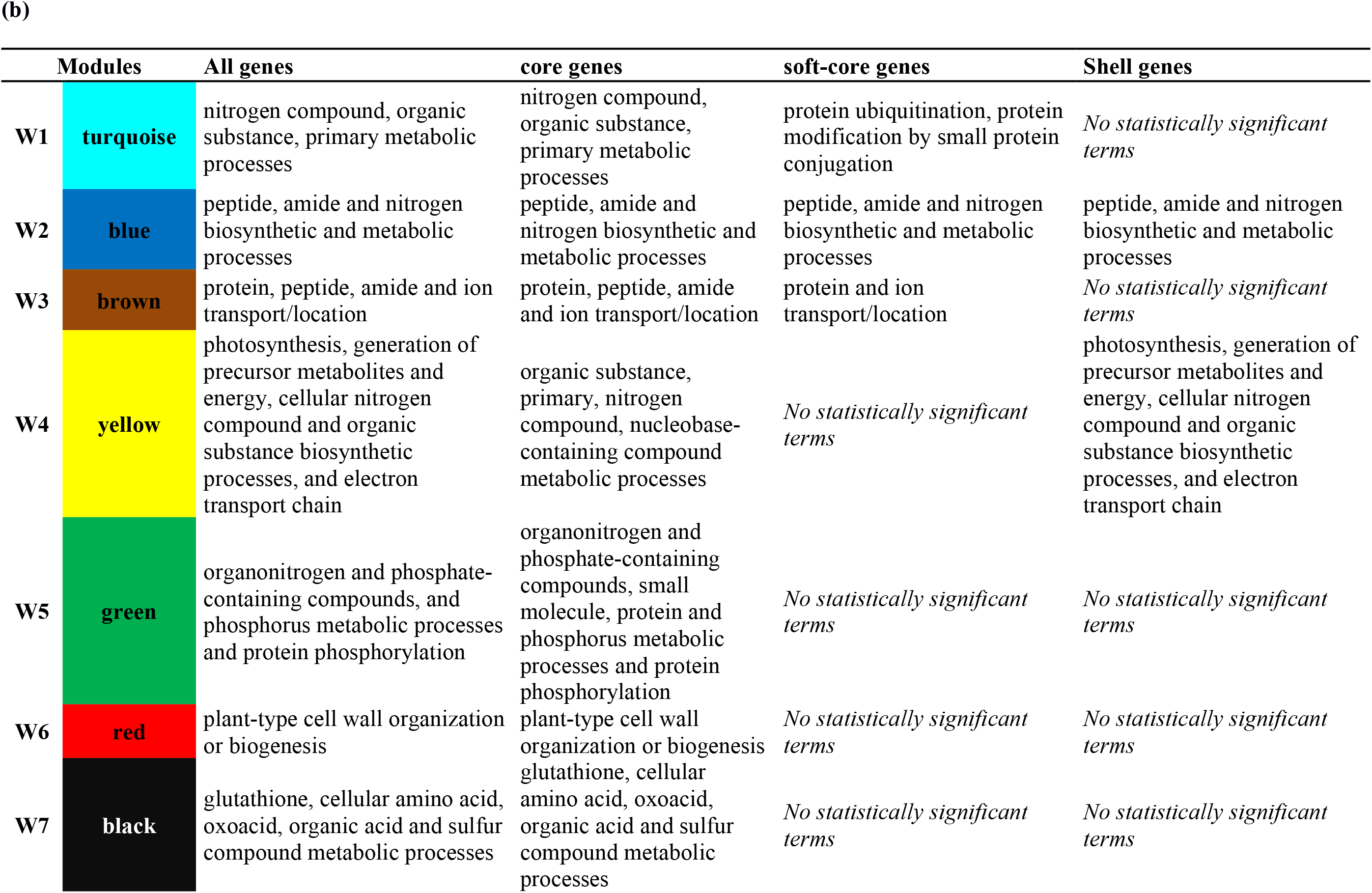
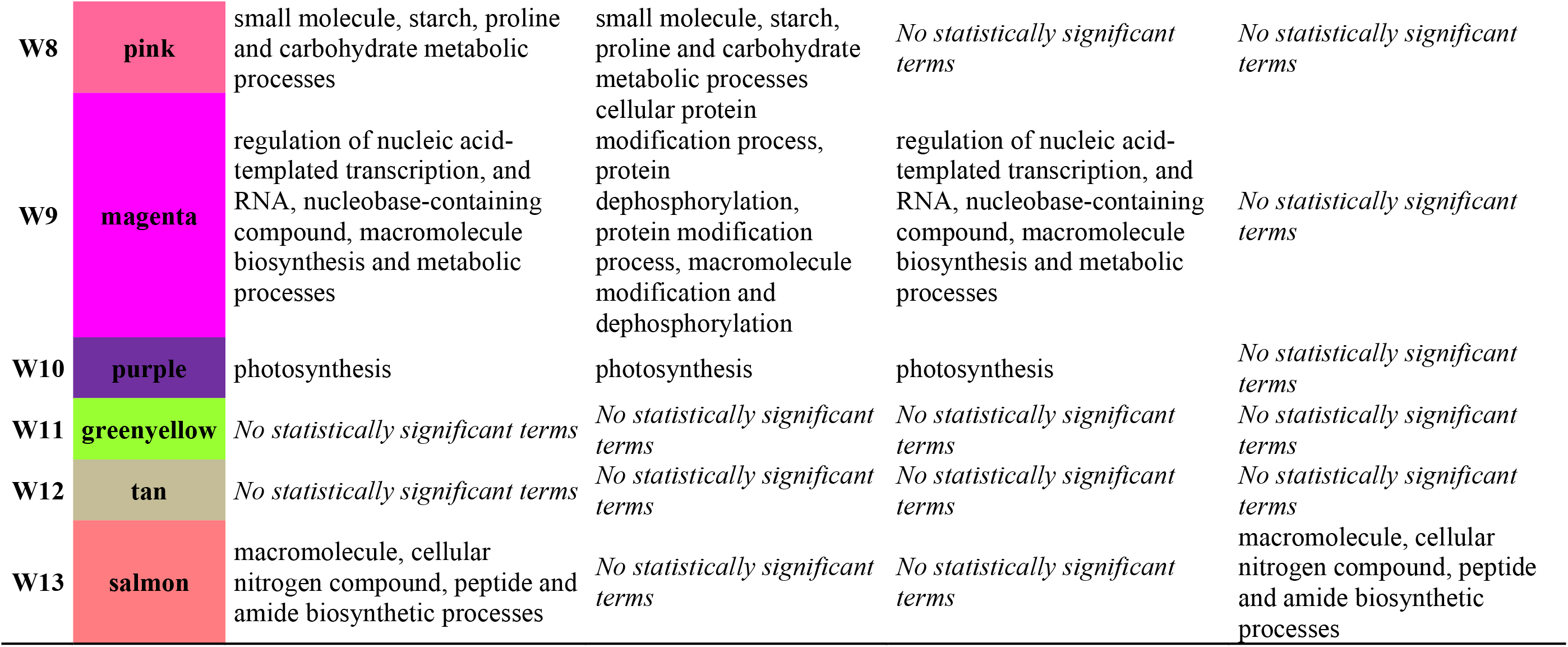
Summary of the enrichment analysis according to the statistically significant GO biological process for the genes (all, core, soft-core and shell genes) clustered in the Drought (D) (**a**) and Water (W) (**b**) modules applying the statistical overrepresentation test of Panther (http://pantherdb.org/) tool. The biological processes are summarized according to the lowest False Discovery Rate (FDR) values (see Supplementary file S3).

### Regulatory motifs of genes in the Drought and Water modules

We detected statistically over-represented sequence motifs upstream of the gene sequences in several modules. These motifs represent putative *cis*-regulatory elements (CREs) located in proximal promoter regions of co-expressed genes in the Drought (Table 3a; Fig. S4a) and Water (Table 3b; Fig. S4b) networks. A generally low but variable proportion of genes harboring putative CREs in their proximal promoters was detected in each module. The drought (Table 3a) and water (Table 3b) modules showed between 3.5-54.2% and 0.1-23% of genes with the predicted CREs, respectively. Genes in the same module shared a conserved regulatory architecture. For example, calmodulin-binding CREs were enriched in the D4 (associated with nucleobase-containing compound, heterocycle and nucleic acid metabolic processes GO terms) and W9 modules (also enriched for nucleobase-containing compound GO processes, among others). We also observed enriched motifs in treatment-specific modules, like the proximal promoters of 27.8% of the genes in module D5 contained CREs similar to those bound by transcription factor B-3 (Table 3a) (Scharf et al., 2012), known to regulate heat shock responses in *Arabidopsis thaliana* (Bechtold et al., 2013; Guo et al., 2016; Nover et al., 2001).

**Table 3.**
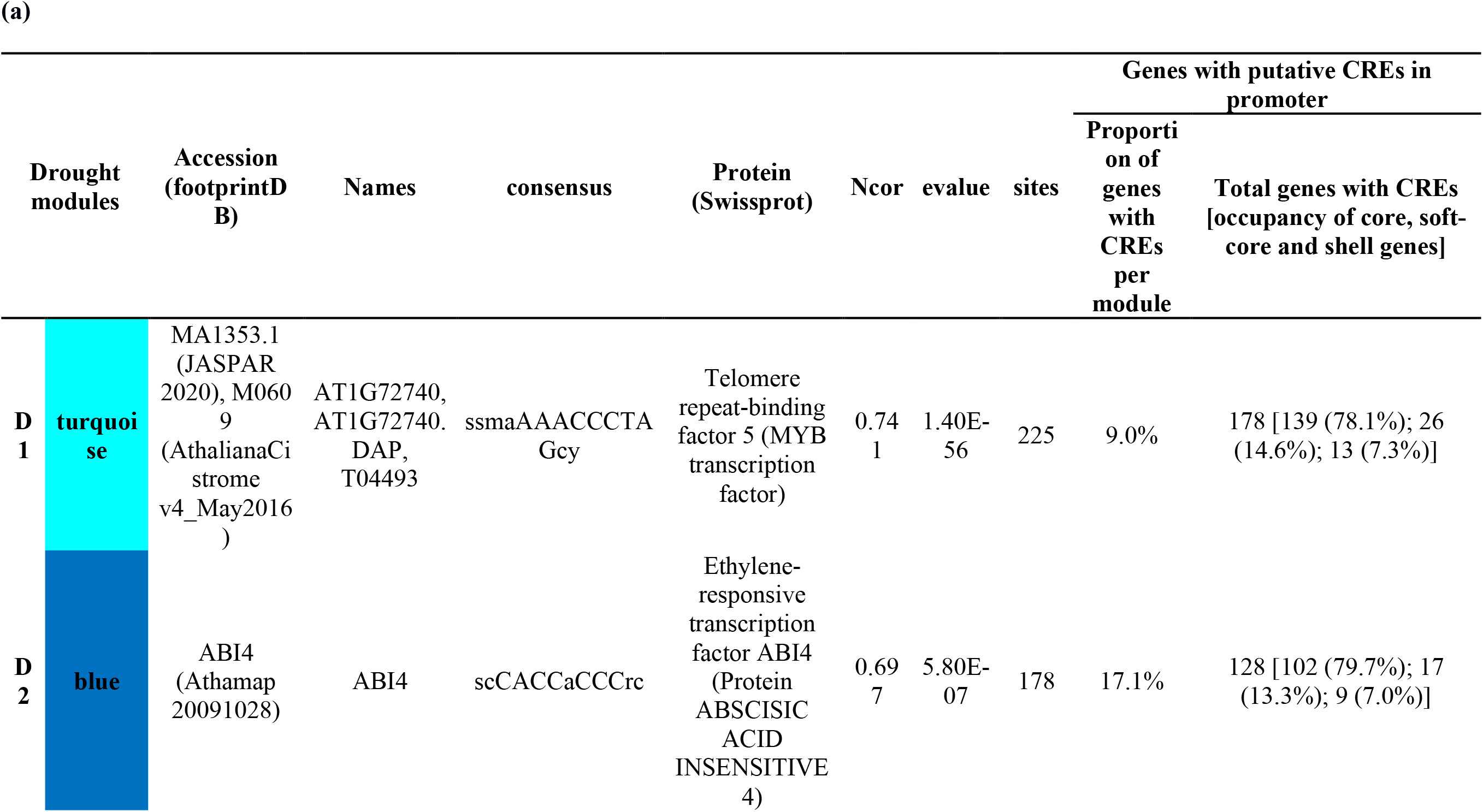
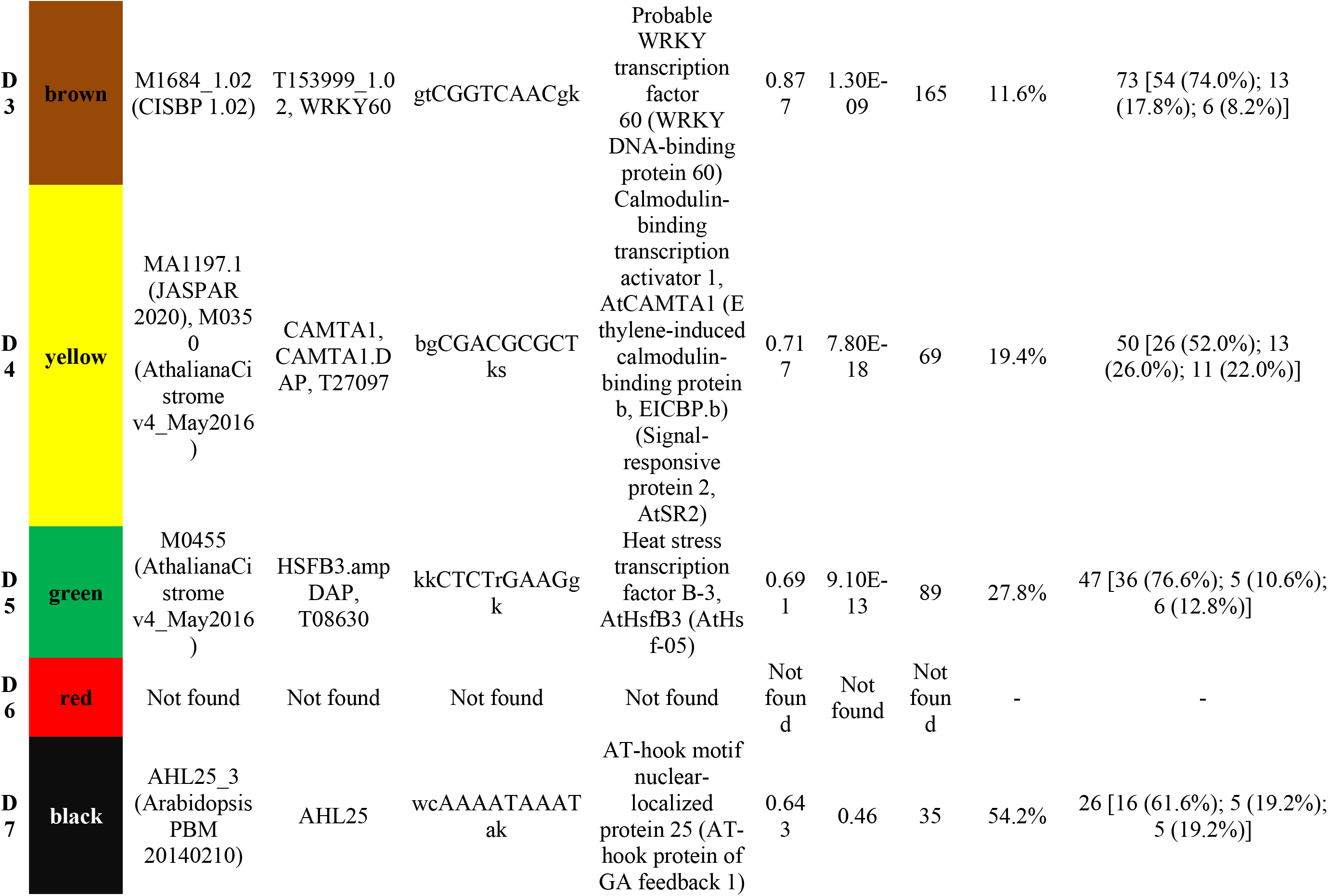
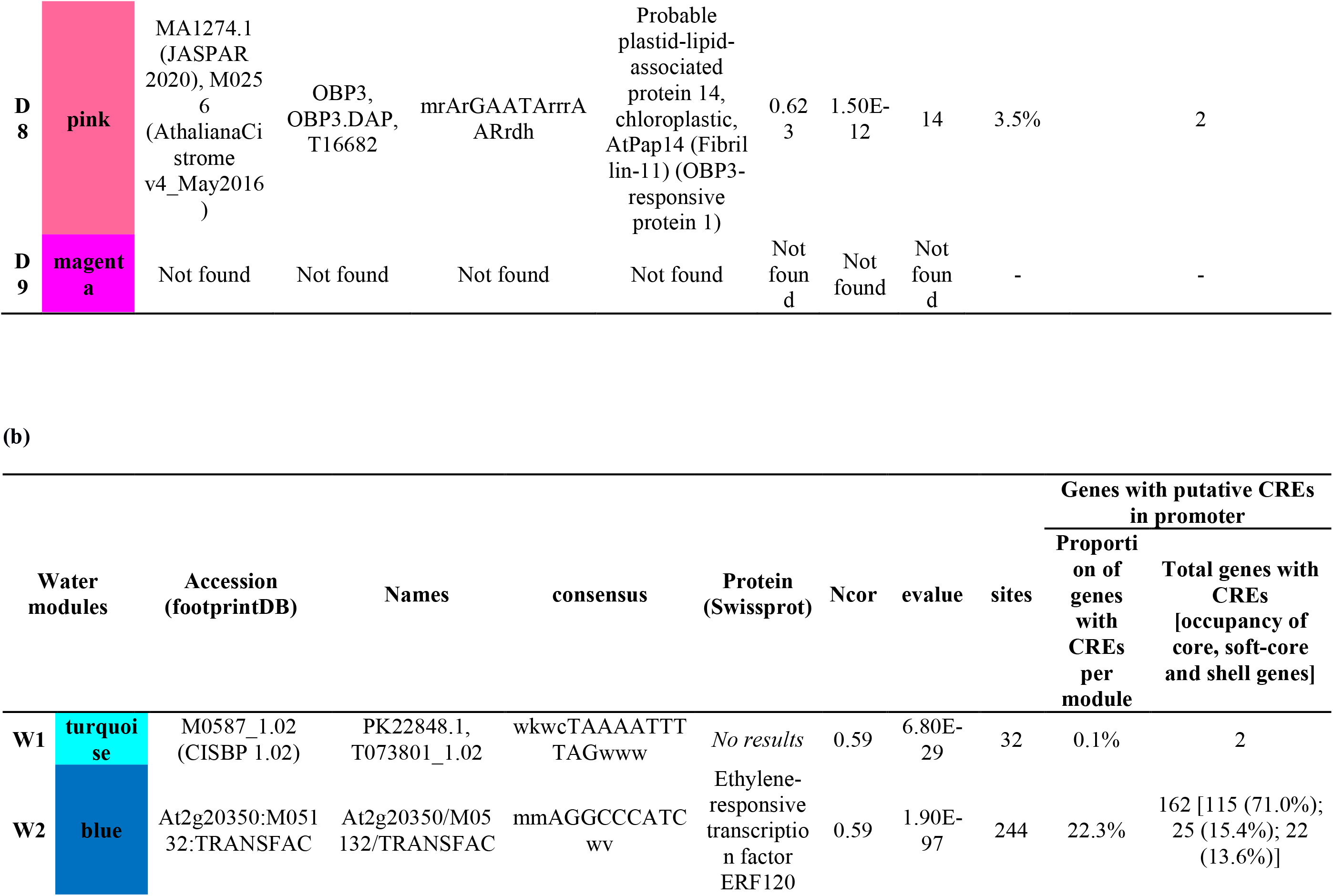
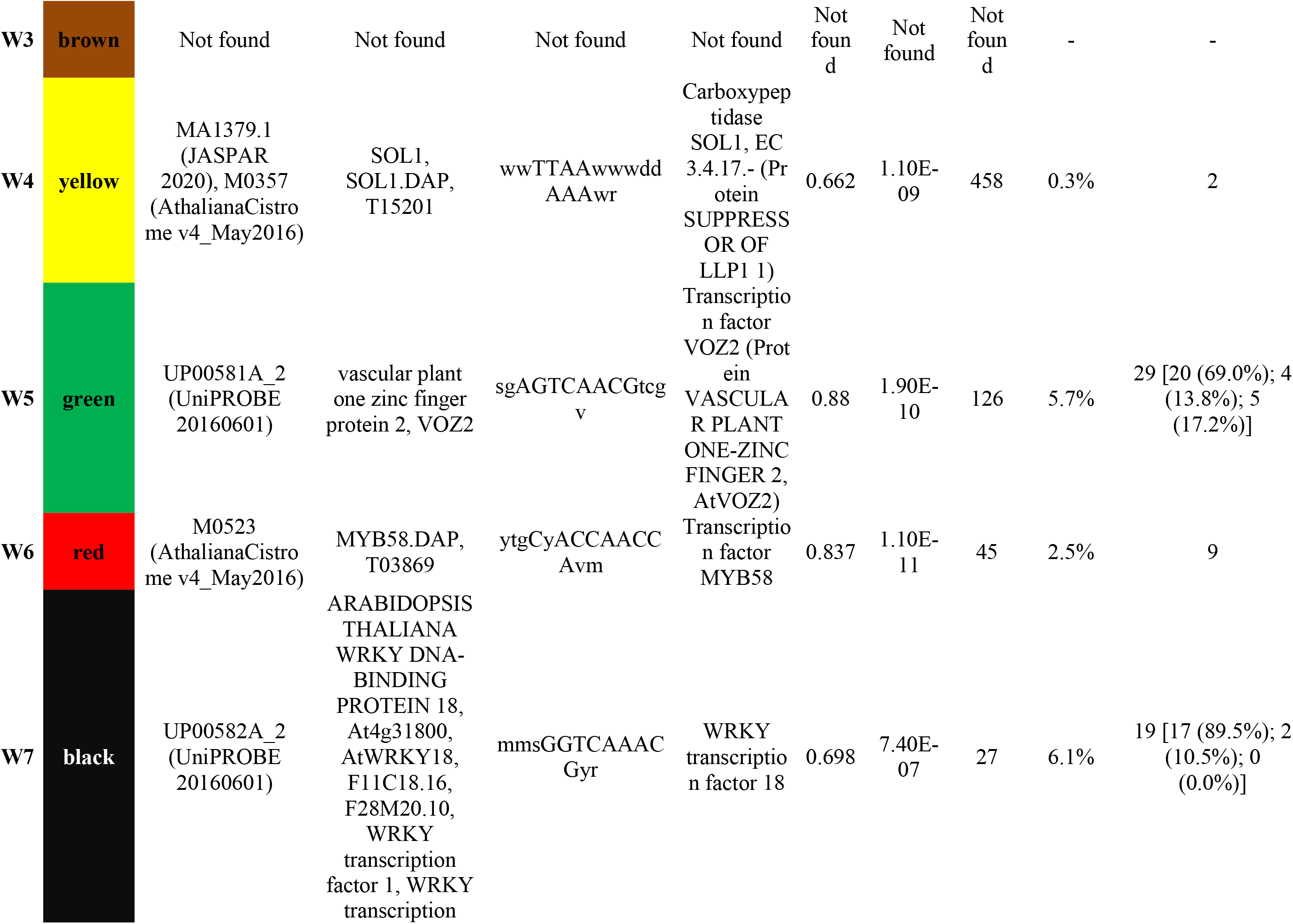
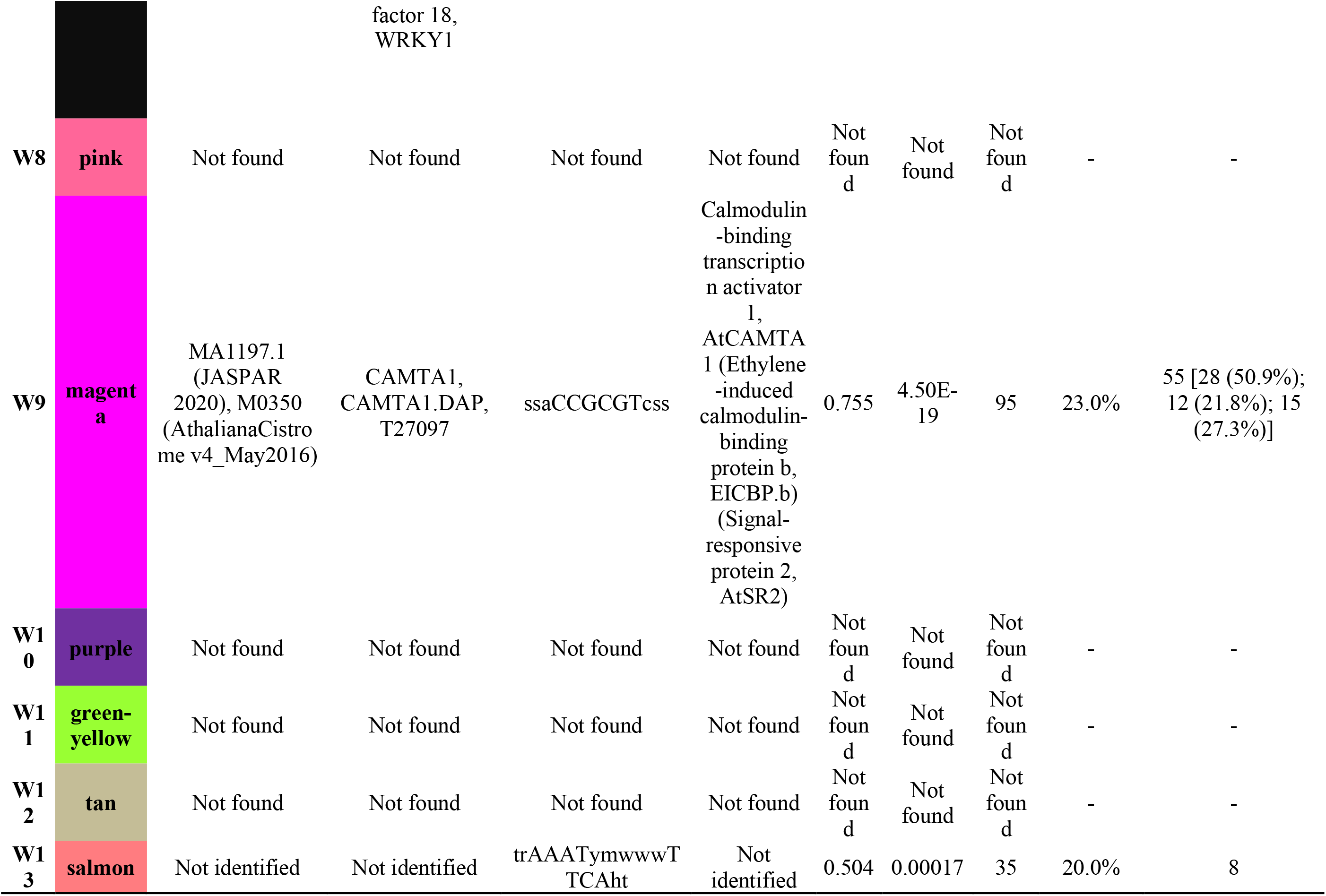
DNA motifs and cis-regulatory elements (CREs) of co-expressed genes assigned to Drought (D) (**a**) and Water (W) (**b**) modules. Modules (numeric and color codes of the modules); Accession (footprintDB), Names (gene name); Consensus sequence of motif; Protein name (Swissprot); Ncor (normalized correlation score); e-value; sites (number of sites used to compile the DNA motif); Proportion of genes with CREs per module as detected by matrix-scan in −500 to +200 bp windows; Total genes with CREs [occupancy of core, soft-core and shell genes]. The dashes indicate that no results were retrieved.

Additionally, all the predicted encoded proteins in each module were analysed to annotate the transcription factors (TF), transcriptional regulators (TR), and kinases (Table S4a,b). For example, TRs annotated in the D5 module – identified above as putatively drought-specific -- are involved in responses to abscisic acid, heat stress, water deprivation and defense, and in zinc, chromatin, and metal ion binding (Table S4a).

### Hub nodes of the Drought and Water networks

Hub nodes were detected in both the Drought (Table 1a) and the Water (Table 1b) networks. A total of 87 (0.53%; 1.73% excluding grey module) hub node isoforms from 72 (0.58%; 1.8% excluding grey module) hub genes were identified in the Drought network, and 251 (1.53%; 3.74% excluding grey module) hub node isoforms from 190 (1.53%; 3.51% excluding grey module) hub genes in the Water network. Roughly, more than twice per-module fraction of hubs was detected in the Water network (1.53% hub nodes/ 1.53% hub genes; Table 1b) compared to the Drought network (0.53% hub nodes/ 0.58% hub genes; Table 1a).

### Pan-genome analyses: occupancy of all clustered and hub genes

The studied pan-genome subset contained 34,310 pan-genome clusters (hereafter “pan-genes”) which were classified by the number of accessions with a gene model represented in a given cluster to determine their occupancy. We found 16,057 (46.8%) core gene clusters with at least one member in every accession. We analyzed these core genes along with the 5,642 (16.4%) clusters from the soft-core pan-genome (occupancy in 31 or 32 accessions) to account for gene annotation errors and uncertainty with orthology assignments. In contrast, there were 12,611 (36.8%) shell genes (i.e., observed in fewer than 31 accessions). Of the 34,310 pan-genes, 15,848 were represented by sequenced RNA tags, after our filtering and normalizing steps. The occupancy of the expressed pan-genes was 12,137 (76.6%) core, 1,869 (11.8%) soft-core and 1,842 (11.6%) shell genes. The discrepancy between the total number of pan-genes and that of genes represented by RNA-Seq tags may have been caused by the filtering of lowly expressed genes. Additionally, many genes in the genome or pan-genome were likely not expressed in leaf tissue at the developmental stage assessed herein. However, our results were consistent with prior studies which found that pan-genome core genes are frequently more highly expressed (Gao et al., 2019; Gordon et al., 2017; Tao et al., 2021) and therefore more likely to be sampled in RNA-Sequencing libraries (Figure S5).

The distribution of pan-gene occupancy varied considerably among modules in both Drought and Water networks (Table 4a,b; Fig. 3a,b). Most modules in both co-expression networks showed ratios of shell genes consistent with the genome averages (Fisher’s Test; Table 4a,b), with notable exceptions. Among Drought modules, D8 showed a considerable excess of shell genes (63.2%). Conversely, D2 (10.0%) and D3 (11.2%) each show a deficit of shell genes. Water co-expression modules showed considerably more variation in pan-gene occupancy. Most large modules in the water network showed significant deficits of shell genes, though W4 (27.8%), W9 (21.8%), W11 (25.5%), W12 (27.5%), and W13 (60.0%) had an excess of shell genes relative to the genome averages (note that W11, W12, and W13 were very small modules with fewer than 60 genes each).

**Figure 3.**
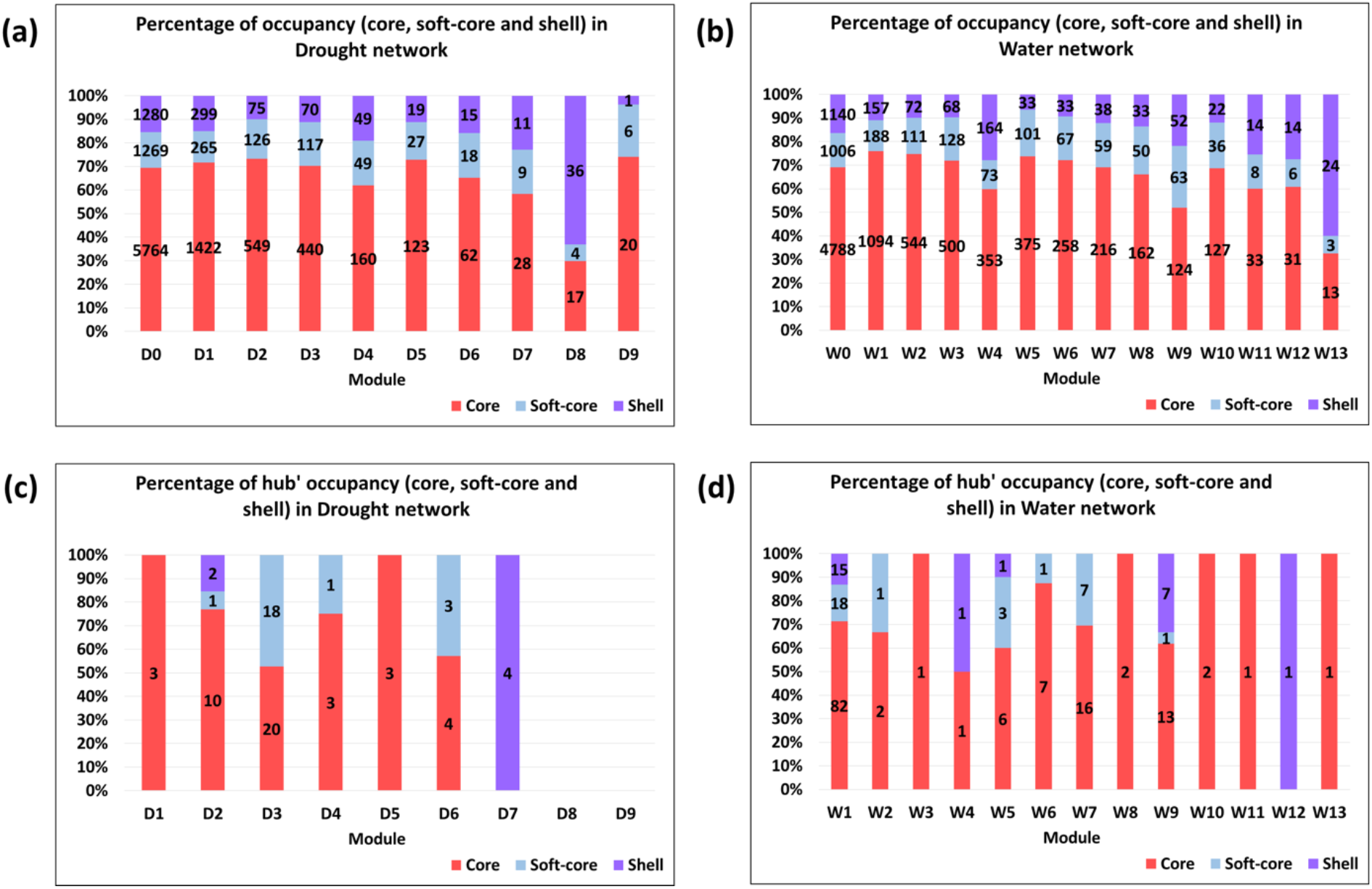
Proportion (%) of occupancies [core (33); soft-core (31-32) and shell (≤30)] of the co-expressed genes (**a; b**) and hub genes (**c; d**) for each Drought (D) and Water (W) network.

**Table 4.**
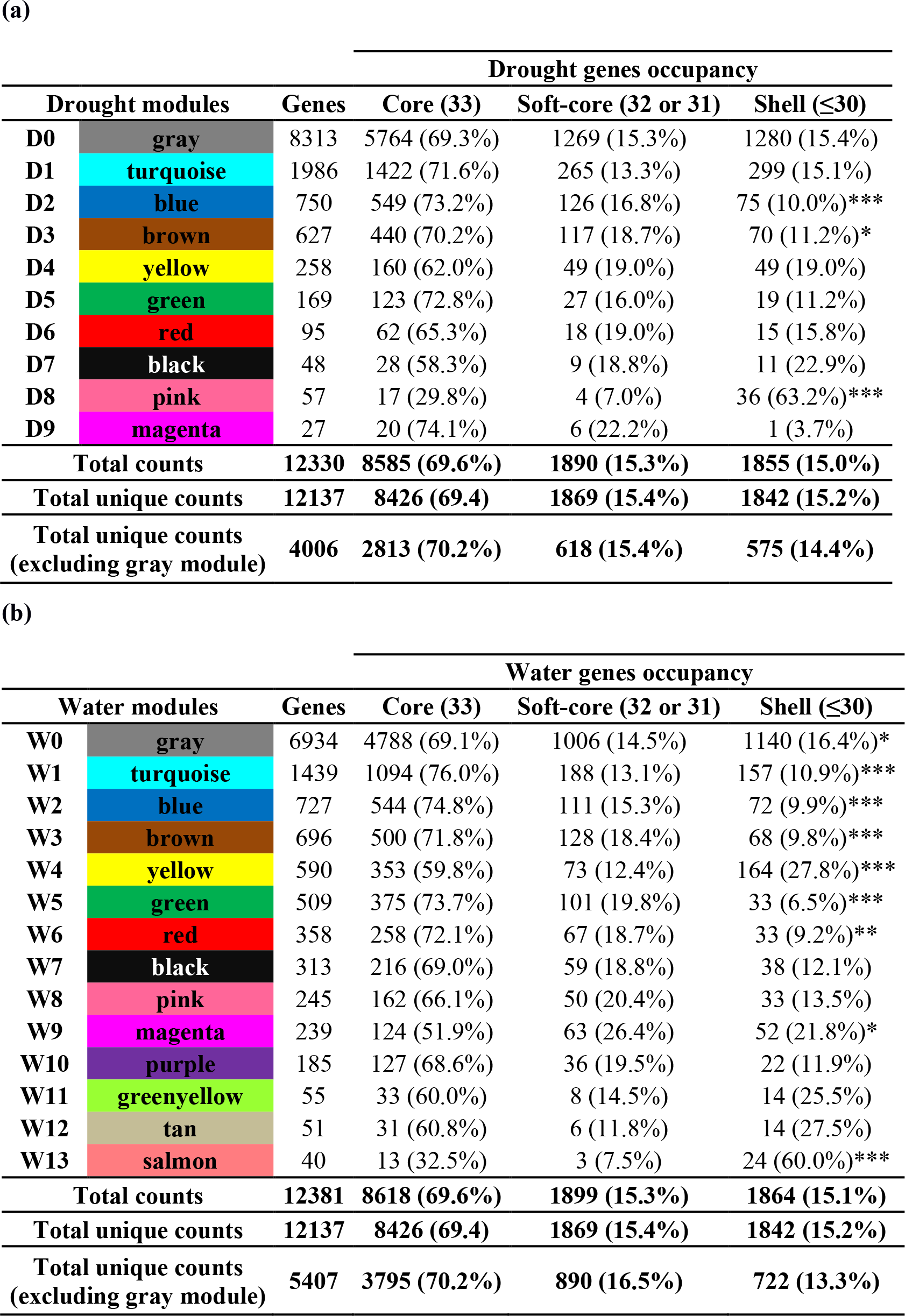
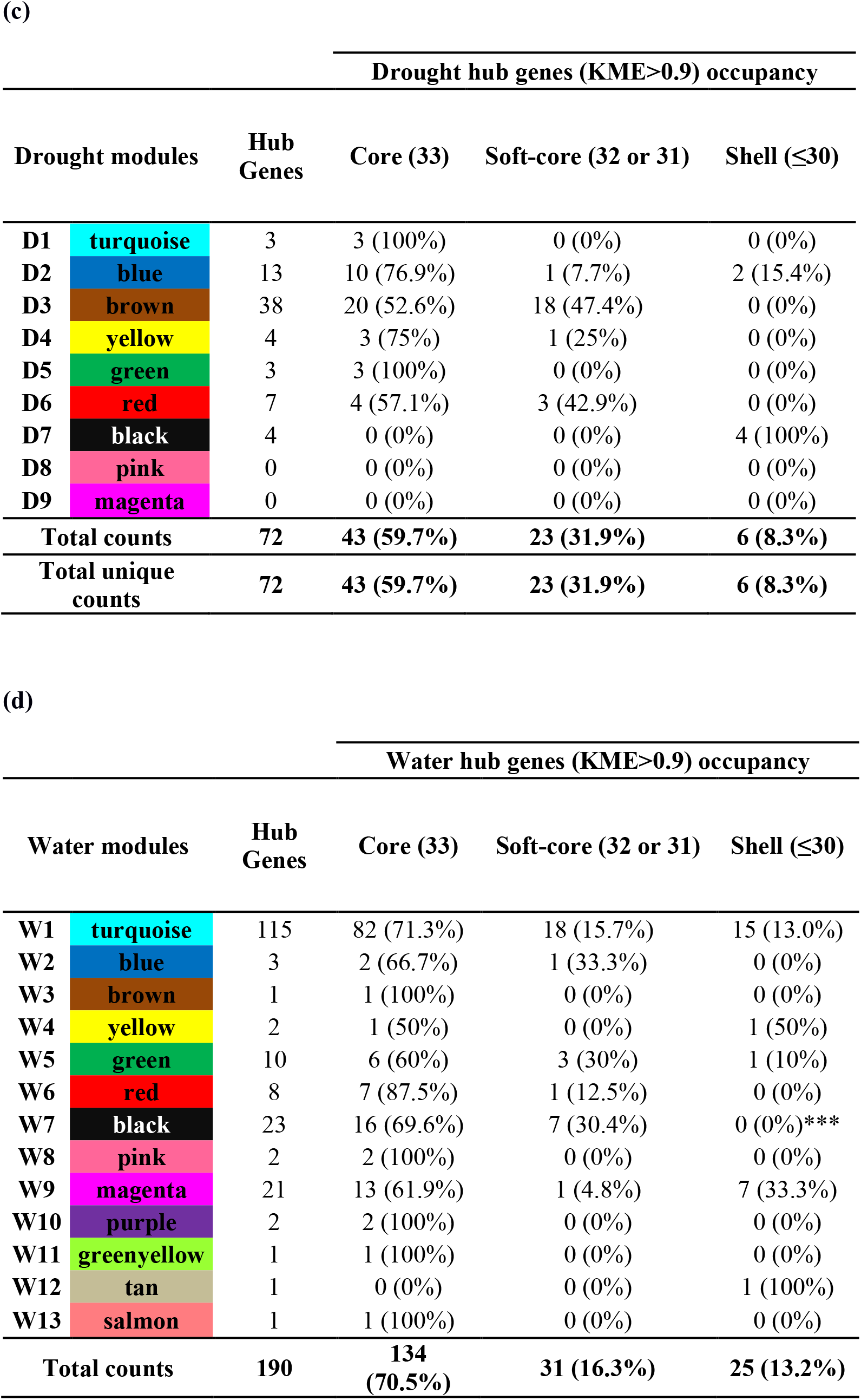
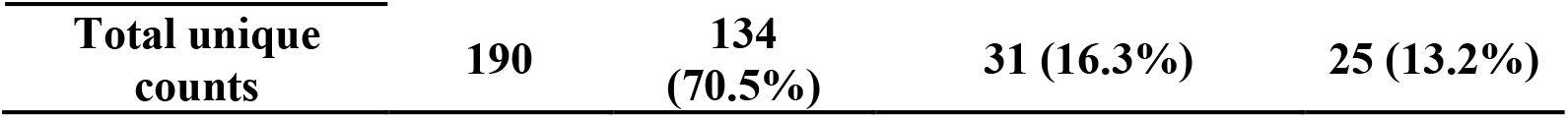
Occupancies [core (33 accessions); soft-core (31-32 accessions) and shell (≤30 accessions)] of the co-expressed genes **(a; b)** and hub genes **(c; d)** for each module of the Drought (D) and Water (W) co-expression networks. Asterisks indicate significant differences comparing shell and total genes between modules and network by Fisher test (*p ≤ 0.05; **p ≤ 0.01; ***p ≤ 0.001).

To investigate the proportion of putative hub genes in each module that were members of the shell gene sets, greater departures from genome averages were required to reach statistical significance given the comparatively small number of these genes (Table 4c,d; Fig. 3c,d). Among the Drought modules, D8 and D9 did not show hub genes whereas other modules showed a predominance (>80%) of core or soft-core hub genes with the exception of D7, whose four hub genes were shell genes (Table 4c). Among the Water modules, W9 had a high proportion of shell hub genes (33.3%) as well as W4 and W12 though these were poorly enriched with hub genes, whereas W7 showed a low percentage of hub shell genes. Only the shell genes of module W7 are significantly under-represented among the hub genes of the Water modules, according to the Fisher test (Table 4d).

As expected, core and soft-core genes were mostly enriched in the same GO terms as those in their respective complete modules (Table 2a,b). Similarly, shell genes corresponded to genes involved in the same biological process as those included in their complete modules. However, the most significant GO term of the shell genes of D1 was photosynthesis, which was not found to be significant among core genes. Similarly, the shell genes of D8 were enriched in photosynthesis, nitrogen, amide, and peptide biosynthetic and metabolic processes terms, while core gene sets did not show any significant enrichments (Table 2a,b and supplementary file S3).

We analyzed the pan-genome occupancy of genes with putative CREs in the modules with at least 10 detected genes. These corresponded predominantly to core genes (>50%) in both networks, whereas 7-22% of the genes with predicted motifs in the Drought network and 0-27.3% in the Water network corresponded to shell genes (Table 3a,b).

### Enrichments of differentially expressed genes with a pan-genomic perspective

72.2%, 17.8% and 10% of the differentially expressed genes with upregulated isoforms in the drought condition were core, soft-core, and shell genes, respectively. On the other hand, 67.5%, 20.9% and 11.6% of the genes with downregulated isoforms in the drought condition were core, soft-core, and shell genes, respectively (Table 5; supplementary file S2). 59.3% of DE isoforms (58.7% of DE genes) and 47.4% DE isoforms (44.6% DE genes) of Drought (Table S5a) and Water (Table S5b) networks, respectively, were not assigned to any modules (i.e., they are members of the gray or zero module). Among the hub nodes (isoforms), 64.4% (56/87) and 34.3% (86/251) of them correspond to DE isoforms of the Drought (Table S6a) and Water (Table S6b) networks, respectively.

**Table 5.** Pan-genome analysis of the differentially expressed (DE) isoforms according to their gene occupancy: core (33 accessions), soft-core (31-32 accessions) and shell (≤30 accessions).* One gene was represented by one up-regulated and one down-regulated isoform.

|  | **Gene occupancy** |  |  |  |
| --- | --- | --- | --- | --- |
|  | **core** | **soft-core** | **shell** | **TOTAL** |
| **upregulated** | 2027 (72.2%) | 500 (17.8%) | 281 (10%) | **2808** |
| **downregulated** | 661 (67.5%) | 205 (20.9%) | 114 (11.6%) | **980** |
| **up/down-regulated** | 1* | 0 | 0 | **1** |
| **TOTAL** | **2689** | **705** | **395** | **3789** |

Five of the nine Drought co-expression modules had a predominance (>50%) of upregulated DE isoforms except for the large modules D2 and D3 and the small modules D8 and D9 (Table S5a). Similarly, among Water modules, only one large module (W4) and three small modules (W10, W11 and W13) had a predominance of downregulated DE isoforms in the drought condition compared to the water condition (Table S5b).

21 out of the 25 most strongly upregulated genes did not cluster with other genes in the drought co-expression network (i.e., they were members of the grey D0 module), while the majority of these strongly upregulated genes did cluster in a module in the water network (Table S7). The 25 most strongly upregulated genes by drought showed a range of predicted functions which included two predicted dehydrins (Decena et al. 2021), two ABA-associated proteins, and two lipid transfer proteins (LTPs) (Table S7). LTPs were among the most highly induced transcripts in an *Arabidopsis thaliana* experiment that imposed very similar soil drying conditions to those imposed in the present study (Des Marais et al., 2012). The 25 most strongly downregulated genes showed markedly different patterns than the most strongly upregulated genes. Most of the genes cluster in D2 and W10, and four of these genes (three clustered in D2 and one in W10) were hub genes (Table S7). Both of these modules were enriched for genes involved in photosynthesis (Table 2a,b) and the annotations for many of these genes suggested associations with the light reactions of photosynthesis.

## DISCUSSION

Large scale transcriptome data sets have been used to construct co-expression networks for gene and gene regulation discovery in model plant systems and crops (Aoki et al, 2007, 2016; Masalia et al., 2017; Miao et al., 2017). The co-expression network approach allows testing hypotheses on gene functions from their putative regulatory interactions with other functionally known genes classified in the same modules (Mochida et al., 2011), and on links between signalling pathways and phenotypic response to environmental stress (Des Marais et al., 2012). However, the mechanisms of and consequences for genetic diversity in environmentally responsive gene regulatory networks are less often considered (Sun & Dinneny, 2018). Our system-level approach allowed us to construct a drought-responsive gene co-expression network from leaf tissue transcriptome profiles of *B. distachyon* accessions and to identify modules of putatively co-regulated genes within it. We integrated these network hypotheses with information about gene presence/absence variation as represented in the *B. distachyon* pan-genome.

### Regulatory control of *Brachypodium* response to soil drying

*B. distachyon* is an annual species native to seasonally dry environments in the Mediterranean, where it has likely evolved mechanisms to tolerate short-term soil drying during the growing season as well as unpredictably timed end-of-season drought (López-Álvarez et al., 2015). Several past studies have identified mechanisms of response to soil drying comprising transcriptomic, metabolic, physiological, and developmental plasticity (Bertolini et al., 2013; Chen et al., 2016; Decena et al., 2021; Des Marais, et al., 2016, 2017b; Fisher et al., 2016; Gordon et al., 2014; Handakumbura et al., 2019; Luo et al., 2011; Manzaneda et al., 2015; Priest et al., 2014; Ruíz et al., 2016; Verelst et al., 2013), as well as considerable genetic diversity of response (GxE; Des Marais et al., 2017b; Handakumbura et al., 2019). Priest et al. (2014) provided the first transcriptomic assessment of response to drying, exposing the Bd21 accession to a simulated severe drying stress by removing plants from soil to desiccate on a lab benchtop. These authors observed a strong transcriptional signature of down-regulated photosynthesis, cell division, and cell growth. Subsequent work imposing a more gradual soil drying stress in Bd21 found the opposite pattern, directly observing sustained cell division and transcriptomic patterns of altered primary metabolism, rather than outright downregulation (Verelst et al., 2013). Indeed, studies imposing moderate drying on diverse *B. distachyon* accessions revealed increased leaf mass per area and greater root biomass in several accessions in response to drying (Des Marais et al., 2017b; Handakumbura et al., 2019), both of which require considerable investment of carbohydrates. In the present study we do, however, observe several strongly downregulated genes with annotated functions related to the light reactions of photosynthesis as well as RuBisCO assembly and function (Table S7).

How can we reconcile these transcriptional signatures of reduced photosynthesis with the observation that carbohydrate-intensive processes like root growth continue under drying? The effects of soil drying on photosynthesis are complex, and the reduction of internal leaf CO_2_ (c_i_) caused by stomatal closure can affect the redox status of cells (Pinheiro & Chaves, 2011). As the Calvin Cycle reduces available CO_2_, it is also a strong sink for energy captured by the photosystems. As this sink is lowered by decreased c_i_, continued high irradiance can lead to increased expression of photoprotective mechanisms and decreased expression of photochemistry as cells try to protect themselves from excess energy (Demmig-Adams & Adams, 1996). As a result, studies of soil drying responses often observe decreased activity of the photosystems and increased expression of, for example, photorespiration or other energy sinks (Wingler et al., 1999). While we have no direct measurements of photorespiration or the quantum yield of photosystem II in the current study, our observation of decreased expression of transcripts associated with photosystem proteins is consistent with these mechanisms.

In light of this past evidence for an important role of photosynthesis and primary metabolism in drying response, we focus here on Drought module 5 (D5). D5 showed a low correlation with the Consensus modules (Fig. 2a), consistent with the hypothesis that the genes in this module are involved in regulating plant response to drought stress. Its co-expressed genes, both core and soft-core, are involved in protein folding, response to heat, temperature, and abiotic stimulus (Table 2a). The four DE hub nodes (three genes) of D5 were upregulated in the drought condition compared to the water condition (Table S8) and each of these four hubs is a core gene. Molecular chaperones, especially heat shock proteins (HSPs), were predominantly annotated in both co-expressed and upregulated DE genes (Table S8). Related to the presence of chaperones, the annotated DNA motifs concurred with the GO enrichment of the protein folding and the Heat stress transcription factor B-3 (Table 3a).

### Topological position of pan-genes

One key conclusion with respect to the evolution of pan-genomes and gene regulation arise from our analysis. First, pan-genes are non-randomly assorted among co-expression modules in *B. distachyon.* This observation suggests that the functions conferred by some co-expression modules are likely under stronger purifying selection than those conferred by other modules. These latter modules represent possible regulatory variation on which natural selection may act. Such a prospect was anticipated by Wagner and Altenberg when they argued that modularity allows for evolutionary tinkering (Wagner & Altenberg, 1996). Our results point towards a role for segregating gene copies in generating this modular variation. This conclusion extends earlier work indicated that low-occupancy genes tend to be enriched for functional classes of genes putatively involved in local adaptation such as disease resistance and gene regulation (Gordon et al., 2017).

Pleiotropy can have a strong effect on the rate of molecular evolution and on the roles that functional gene variants might play in evolutionary change. Pleiotropy is often correlated with the position of a gene or protein in biochemical and gene regulatory networks (Erwin & Davidson, 2009; Jeong et al., 2001), which are now readily inferred from high dimensional datasets such as the genome-wide gene co-expression networks studies herein. Here, we considered the case of potentially large-effect mutations – segregating gene copies identified from a grass pan-genome – and ask whether such “pan-genes” are unevenly distributed in gene co-expression networks. Focusing on the well-Watered (control) environment, we find that shell genes – pan-genes found in fewer than 31 of our studied accessions -- are statistically under-represented among the genes in five of the six largest (in terms of total number of genes) co-expression modules (Table 4b). These large modules are enriched for GO terms comprising essential processes such as protein synthesis, primary metabolism, various processes related to phosphorus metabolism and signalling, and cell wall organization (Table 2b). Moreover, shell genes are generally under-represented among module hub genes (diagnosed as those whose expression most highly correlated with the module as a whole, and thus possibly the most topologically connected among genes in a module) in these five Water modules (Table 4d). Collectively, these results support the hypothesis that core pan-genome genes are centrally located in gene co-expression networks and involved in biological processes likely to be under strong purifying selection.

Water module 4 (W4), comprising 590 genes, is the only large water module that is statistically enriched for shell genes (Table 4b). Shell genes in W4 are associated with a range of GO terms including processes related to photosynthesis (Table 2b). In general, the lists of shell genes in modules do not tend to have strong GO enrichments, perhaps owing to the relatively small numbers of genes in these lists. Interestingly, among the Drought modules, the only module for which shell genes do have an enrichment (D8; Table 4a) includes GO terms associated with photosynthesis (Table 2a). Shell genes represent genes found in some sampled accessions but missing in others, suggesting that *B. distachyon* may harbor genetic diversity in molecular pathways related to photosynthesis. Whether these segregating variants represent adaptive genetic diversity reflecting the broad geographical coverage of our sampling, or simply mildly deleterious copy number variants on their way to being lost will require a larger population sample. Previously, we demonstrated significant genetic variation among the same *Brachypodium* accessions used herein for leaf carbon content, leaf C:N ratios, and water use efficiency (WUE; Des Marais, et al.; 2017b). Among these, WUE was significantly associated with principal components summarizing climate diversity; it is possible that some of the segregating variation in photosynthesis gene presence/absence is involved in local adaptation to climate.

The co-expression network approach we employed here, in concert with a modern pan-genomic perspective on segregating genetic variation, allows us to identify subsets of genes involved in core metabolic processes such as photosynthesis, suggesting possible candidate genes regulating natural genetic variation in resource assimilation and growth. These genes make attractive targets for further hypothesis testing via genome editing. Collectively, our work demonstrates the importance of accounting for both gene copy number variation and regulatory interactions in studying genome function and evolution.

## ACKNOWLEDGEMENTS

We thank R. Hopkins, E. Sukamtoh, J. Bonnette, and B. Whitaker for assistance with data collection. This work was supported by the USDA (NIFA-2011-67012-30663) to D.L.D., NSF (IOS-0922457) to T.E.J., the Spanish Ministry of Science and Innovation CGL2016-79790-P and PID2019-108195GB-I00, University of Zaragoza UZ2016_TEC02 grant projects to P.C., and the Joint Genome Institute FP00006675 contract to P.C. and FP00006746 to D.L.D.. R.S. was funded by a Spanish Ministry of Economy and Competitiveness (Mineco) FPI PhD fellowship, Mineco and Ibercaja-CAI mobility grants and Instituto de Estudios Altoaragoneses grant. B.C.M. was funded by Fundación ARAID. P.C. and R.S. were partially funded by a European Social Fund/Spanish Aragón Government Bioflora grant.

## DATA ACCESSIBILITY AND BENEFIT-SHARING

### Data Accessibility

RNA Sequence data are available at the ENA (European Nucleotide Archive; https://www.ebi.ac.uk/ena) with the consecutive accession numbers from ERR6133302 to ERR6133575.

All the R scripts, supplementary Excel files, TPM counts and adjacency matrices are available in the Github repository identified by the following doi: xxxxxxxxxx created using Zenodo.

Experimental details and metadata for samples studied herein were reported previously (Des Marais et al. 2017b).

### Benefit Sharing

All accessions used in the current study originated from collections generated by the *Brachypodium* research community over several decades. Benefits from this research accrue from the sharing of our data and results on public databases as described above.

## AUTHOR CONTRIBUTION

D.L.D. and T.E.J. designed the research. D.L.D. performed the experimental manipulations and generated the data. D.L.D., R.S., and B.C.-M. performed the analyses. D.L.D. and R.S. wrote the primary draft of the manuscript, and all authors contributed to the final draft of the manuscript.

