## Supplementary file for "Patterns of pan-genome occupancy and gene co-expression under water-deficit in *Brachypodium distachyon*"

### Supplementary Tables

**Table S1.** Natural accessions of *Brachypodium distachyon* used in the study. Information on elevation (meters above sea level, masl), latitude and longitude of collection sites.

| Accession | Collection location | Elevation (masl) | Latitude | Longitude |
|---|---|---|---|---|
| ABR2 | Hérault, France | 371 | 43° 36' 15.343" N | 3° 15' 46.580" E |
| ABR3 | Aísa, Huesca, Spain | 1928 | 42° 10' 49.8" N | 0° 4' 23.2" W |
| ABR4 | Arén, Huesca, Spain | 480 | 42° 15' 45.54" N | 0° 43' 0.48" E |
| ABR5 | Jaca, Huesca, Spain | 828 | 42° 34' 23.45" N | 0° 33' 49.39" W |
| ABR6 | Los Arcos, Navarra, Spain | 484 | 42° 34' 27.48" N | 2° 11' 5.39" W |
| ABR8 | Siena, Italy | 272 | 43° 18' 52.423" N | 11° 19' 10.902" E |
| Adi10 | Adiyaman, Turkey | 510 | 37° 46' 14.5" N | 38° 21' 8.2" E |
| Adi12 | Adiyaman, Turkey | 510 | 37° 46' 14.5" N | 38° 21' 8.2" E |
| Adi2 | Adiyaman, Turkey | 510 | 37° 46' 14.5" N | 38° 21' 8.2" E |
| Bd1-1 | Soma, Manisa, Turkey | 141 | 39° 11' 27.44" N | 27° 36' 28.59" E |
| Bd18-1 | Kaman, Kırşehir Province, Turkey | 1101 | 39° 22' 4.25" N | 33° 43' 48.91" E |
| Bd21 | near Salakudin, Iraq | 42 | 33° 45' 39.18" N | 44° 24' 11.07" E |
| Bd21-3 | near Salakudin, Iraq | 42 | 33° 45' 39.18" N | 44° 24' 11.07" E |
| Bd2-3 | Iraq | 42 | 33° 45' 39.18" N | 44° 24' 11.07" E |
| Bd30-1 | Dilar, Granada, Spain | 1220 | 36° 59' 25.76" N | 3° 33' 31.44" W |
| Bd3-1 | Iraq | 42 | 33° 45' 39.18" N | 44° 24' 11.07" E |
| BdTR10c | Turkey | 1288 | 37° 46' 41.64" N | 31° 53' 5.68" E |
| BdTR11g | Kirklareli, Turkey | 124 | 41° 25' 17.86" N | 27° 28' 36.81" E |
| BdTR11i | Turkey | 363 | 39° 44' 17.39" N | 28° 2' 24.71" E |
| BdTR12c | Turkey | 1035 | 39° 44' 53.45" N | 34° 39' 1.15" E |
| BdTR13a | Ankara, Turkey | 787 | 39° 45' 23.35" N | 32° 25' 56.46" E |
| BdTR1i | Aydin, Turkey | 841 | 38° 5' 35.03" N | 28° 34' 59.02" E |
| BdTR2b | Turkey | 667 | 40° 4' 55.55" N | 31° 19' 52.01" E |
| BdTR2g | Ankara, Turkey | 1596 | 40° 23' 37.13" N | 32° 59' 7.32" E |
| BdTR3c | Turkey | 1957 | 36° 46' 58.92" N | 32° 57' 46.71" E |
| BdTR5i | Turkey | 1596 | 40° 23' 37.13" N | 32° 59' 7.32" E |
| BdTR9k | Eskişehir, Turkey | 932 | 39° 45' 10.62" N | 30° 47' 19.07" E |
| Bis1 | Bismil, Turkey | 529 | 37° 52' 35.6" N | 41° 0' 54.3" E |
| Kah1 | Kahta, Turkey | 665 | 37° 44' 2.3" N | 38° 32' 0.2" E |
| Kah5 | Kahta, Turkey | 665 | 37° 44' 2.3" N | 38° 32' 0.2" E |
| Koz1 | Kozluk, Turkey | 853 | 38° 9' 8.2.6" N | 41° 36' 34.8" E |
| Koz3 | Kozluk, Turkey | 853 | 38° 9' 8.2.6" N | 41° 36' 34.8" E |
| Ron2 | Roncal, Navarra, Spain | 594 | 42° 46' 50" N | 0° 57' 48'' W |

**Table S2.** RNA sequencing (RNA-Seq) data and drought/water experimental design information. Accessions; Raw SE (raw single-end reads); Filtered SE (filtered single-end reads); RNA extraction session (arbitrary “date”: “a”, “b”, “c”, “d”, “e”, “f”, “g”, “h”, “i”); condition (drought: D; water: W).

| Accessions | Raw SE | Filtered SE | date | condition |
|---|---|---|---|---|
| ABR2 | 6,601,127 | 1,667,289 | e | D |
| ABR2 | 5,746,463 | 2,222,313 | f | D |
| ABR2 | 4,804,333 | 3,147,028 | a | D |
| ABR2 | 4,224,149 | 2,595,889 | d | D |
| ABR2 | 6,489,984 | 2,092,468 | f | W |
| ABR2 | 1,784,829 | 1,063,952 | g | W |
| ABR2 | 9,874,691 | 3,725,483 | b | W |
| ABR2 | 3,846,175 | 3,164,341 | c | W |
| ABR3 | 4,625,149 | 2,900,061 | f | D |
| ABR3 | 3,151,382 | 2,788,129 | g | D |
| ABR3 | 5,725,171 | 4,069,591 | a | D |
| ABR3 | 5,398,049 | 4,056,956 | d | D |
| ABR3 | 3,653,050 | 937,687 | f | W |
| ABR3 | 11,283,377 | 5,216,959 | f | W |
| ABR3 | 3,858,632 | 2,705,509 | d | W |
| ABR3 | 2,157,282 | 1,444,249 | e | W |
| ABR4 | 7,774,753 | 2,116,971 | e | D |
| ABR4 | 1,978,027 | 1,439,774 | h | D |
| ABR4 | 8,113,578 | 5,410,834 | b | D |
| ABR4 | 8,290,472 | 6,252,911 | e | D |
| ABR4 | 4,772,086 | 4,265,809 | g | W |
| ABR4 | 3,402,337 | 2,799,591 | h | W |
| ABR4 | 7,152,112 | 3,173,759 | b | W |
| ABR4 | 2,723,962 | 1,649,000 | b | W |
| ABR5 | 1,518,507 | 1,059,416 | g | D |
| ABR5 | 6,700,459 | 3,350,008 | h | D |
| ABR5 | 8,415,687 | 3,613,083 | a | D |
| ABR5 | 6,372,579 | 5,203,682 | d | D |
| ABR5 | 4,243,346 | 3,411,799 | g | W |
| ABR5 | 3,074,709 | 2,519,731 | h | W |
| ABR5 | 2,960,971 | 1,520,797 | e | W |
| ABR5 | 6,207,230 | 5,076,436 | e | W |
| ABR6 | 38,184,219 | 5,211,877 | f | D |
| ABR6 | 7,337,310 | 4,028,305 | h | D |
| ABR6 | 8,407,721 | 4,506,524 | b | D |
| ABR6 | 6,849,023 | 4,180,661 | d | D |
| ABR6 | 5,861,023 | 2,762,096 | f | W |
| ABR6 | 4,014,346 | 2,894,224 | g | W |
| ABR6 | 1,572,086 | 726,106 | d | W |
| ABR6 | 3,770,757 | 3,115,437 | e | W |
| ABR8 | 2,180,204 | 1,433,019 | f | D |
| ABR8 | 5,980,806 | 4,808,335 | h | D |
| ABR8 | 4,735,655 | 3,098,240 | d | D |
| ABR8 | 1,859,648 | 1,440,451 | e | D |

| | | | | |
|---|---|---|---|---|
| ABR8 | 3,581,107 | 2,791,425 | e | W |
| ABR8 | 5,048,561 | 4,447,005 | h | W |
| ABR8 | 2,869,311 | 2,334,201 | a | W |
| ABR8 | 3,629,754 | 2,176,382 | d | W |
| Adi10 | 9,078,234 | 7,870,859 | g | D |
| Adi10 | 11,252,107 | 8,816,557 | h | D |
| Adi10 | 5,609,457 | 2,720,611 | b | D |
| Adi10 | 6,469,852 | 3,074,409 | b | D |
| Adi10 | 5,166,031 | 2,863,430 | f | W |
| Adi10 | 4,485,655 | 3,153,124 | h | W |
| Adi10 | 7,357,845 | 5,076,624 | c | W |
| Adi10 | 4,229,599 | 2,998,656 | d | W |
| Adi12 | 3,315,119 | 2,365,916 | g | D |
| Adi12 | 5,397,127 | 3,730,142 | h | D |
| Adi12 | 9,718,366 | 5,938,112 | b | D |
| Adi12 | 6,297,978 | 4,527,496 | d | D |
| Adi12 | 5,337,436 | 1,167,870 | f | W |
| Adi12 | 7,483,059 | 4,806,165 | f | W |
| Adi12 | 5,214,017 | 2,600,489 | b | W |
| Adi12 | 3,053,578 | 1,436,316 | b | W |
| Adi2 | 7,134,806 | 5,658,048 | h | D |
| Adi2 | 1,954,259 | 1,614,746 | h | D |
| Adi2 | 6,758,134 | 3,379,245 | b | D |
| Adi2 | 3,395,135 | 2,943,975 | d | D |
| Adi2 | 2,906,937 | 2,464,347 | e | W |
| Adi2 | 4,695,911 | 3,948,557 | g | W |
| Adi2 | 7,454,548 | 6,916,945 | e | W |
| Bd1-1 | 4,922,864 | 2,007,767 | f | D |
| Bd1-1 | 6,601,540 | 5,082,930 | g | D |
| Bd1-1 | 6,601,084 | 2,998,518 | a | D |
| Bd1-1 | 1,983,400 | 1,149,142 | d | D |
| Bd1-1 | 4,801,056 | 4,145,854 | e | W |
| Bd1-1 | 4,995,276 | 3,971,153 | g | W |
| Bd1-1 | 4,890,577 | 2,222,735 | b | W |
| Bd18-1 | 1,209,291 | 852,834 | g | D |
| Bd18-1 | 4,300,181 | 3,294,256 | d | D |
| Bd18-1 | 7,005,425 | 1,677,013 | e | W |
| Bd18-1 | 2,234,251 | 1,486,291 | g | W |
| Bd18-1 | 6,450,898 | 3,558,449 | c | W |
| Bd18-1 | 11,453,626 | 6,881,259 | c | W |
| Bd21 | 9,012,664 | 6,908,706 | g | D |
| Bd21 | 8,393,000 | 6,685,084 | g | D |
| Bd21 | 4,388,610 | 2,454,344 | c | D |
| Bd21 | 3,209,402 | 2,440,369 | e | D |
| Bd21 | 3,890,732 | 3,443,385 | e | W |
| Bd21 | 2,731,571 | 1,598,308 | g | W |
| Bd21 | 5,577,765 | 3,037,099 | c | W |
| Bd21 | 7,866,176 | 6,578,328 | d | W |
| Bd21-3 | 4,373,022 | 1,664,189 | f | D |

| | | | | |
|---|---|---|---|---|
| Bd21-3 | 11,184,245 | 8,830,207 | h | D |
| Bd21-3 | 3,725,637 | 2,509,645 | d | D |
| Bd21-3 | 7,060,460 | 4,794,417 | d | D |
| Bd21-3 | 2,704,777 | 1,424,564 | g | W |
| Bd21-3 | 9,189,178 | 3,010,421 | b | W |
| Bd21-3 | 5,597,966 | 3,521,383 | e | W |
| Bd2-3 | 6,426,336 | 3,985,982 | e | D |
| Bd2-3 | 7,233,290 | 5,270,217 | g | D |
| Bd2-3 | 5,158,114 | 2,190,269 | a | D |
| Bd2-3 | 4,562,719 | 3,905,164 | d | D |
| Bd2-3 | 4,698,918 | 4,091,484 | g | W |
| Bd2-3 | 6,104,643 | 5,397,405 | g | W |
| Bd2-3 | 4,932,312 | 4,315,897 | d | W |
| Bd2-3 | 672,413 | 377,700 | d | W |
| Bd30-1 | 15,450,940 | 8,648,687 | f | D |
| Bd30-1 | 7,739,481 | 6,118,678 | g | D |
| Bd30-1 | 1,926,448 | 1,521,180 | c | D |
| Bd30-1 | 5,678,800 | 4,691,441 | e | D |
| Bd30-1 | 17,729,697 | 13,269,740 | f | W |
| Bd30-1 | 5,724,855 | 5,259,550 | g | W |
| Bd30-1 | 4,774,069 | 3,231,139 | a | W |
| Bd30-1 | 6,729,545 | 4,714,110 | a | W |
| Bd3-1 | 7,153,771 | 3,363,646 | f | D |
| Bd3-1 | 7,847,984 | 5,946,898 | f | D |
| Bd3-1 | 6,279,816 | 5,731,603 | e | D |
| Bd3-1 | 1,516,791 | 1,088,093 | f | W |
| Bd3-1 | 4,861,752 | 1,968,962 | f | W |
| Bd3-1 | 2,790,729 | 2,020,517 | a | W |
| Bd3-1 | 7,500,512 | 5,520,814 | a | W |
| BdTR10c | 4,833,695 | 3,959,199 | f | D |
| BdTR10c | 8,262,740 | 3,977,196 | f | D |
| BdTR10c | 1,993,453 | 1,323,181 | d | D |
| BdTR10c | 6,191,972 | 4,901,492 | e | D |
| BdTR10c | 2,634,263 | 1,940,567 | e | W |
| BdTR10c | 5,139,972 | 1,885,709 | f | W |
| BdTR10c | 2,498,383 | 1,355,283 | d | W |
| BdTR10c | 1,790,136 | 1,192,950 | d | W |
| BdTR11g | 6,265,037 | 4,947,291 | f | D |
| BdTR11g | 8,131,691 | 6,832,907 | h | D |
| BdTR11g | 5,063,729 | 2,509,484 | c | D |
| BdTR11g | 7,408,854 | 6,200,830 | d | D |
| BdTR11g | 4,149,389 | 756,407 | f | W |
| BdTR11g | 8,603,073 | 6,133,854 | g | W |
| BdTR11g | 3,415,642 | 2,633,429 | a | W |
| BdTR11g | 6,786,389 | 4,707,163 | b | W |
| BdTR11i | 5,819,425 | 842,218 | f | D |
| BdTR11i | 2,378,944 | 1,592,929 | g | D |
| BdTR11i | 4,136,864 | 2,389,466 | d | D |
| BdTR11i | 2,559,294 | 1,126,434 | d | D |

| | | | | |
|---|---|---|---|---|
| BdTR11i | 11,864,916 | 6,251,158 | e | W |
| BdTR11i | 4,231,383 | 3,079,181 | i | W |
| BdTR11i | 5,684,403 | 2,986,219 | b | W |
| BdTR11i | 2,671,800 | 1,639,304 | e | W |
| BdtR12c | 6,176,408 | 1,353,488 | e | D |
| BdTR13a | 3,360,398 | 2,395,414 | e | D |
| BdTR13a | 1,894,752 | 1,223,359 | g | D |
| BdTR13a | 8,824,689 | 5,023,187 | c | D |
| BdTR13a | 1,860,137 | 1,349,117 | e | D |
| BdTR13a | 9,513,938 | 3,604,562 | e | W |
| BdTR13a | 5,531,061 | 1,859,726 | f | W |
| BdTR13a | 4,519,318 | 3,247,705 | a | W |
| BdTR13a | 3,950,096 | 3,098,152 | e | W |
| BdTR1i | 4,696,550 | 2,983,773 | g | D |
| BdTR1i | 5,377,545 | 4,540,394 | g | D |
| BdTR1i | 6,541,054 | 4,913,208 | d | D |
| BdTR1i | 2,626,574 | 1,432,016 | d | D |
| BdTR1i | 5,647,332 | 5,068,792 | g | W |
| BdTR1i | 5,969,862 | 5,321,731 | g | W |
| BdTR1i | 4,937,157 | 3,015,216 | c | W |
| BdTR1i | 4,294,152 | 3,640,765 | e | W |
| BdTR2b | 5,387,175 | 4,634,162 | e | D |
| BdTR2b | 4,369,920 | 4,017,001 | g | D |
| BdTR2b | 5,347,090 | 3,816,668 | a | D |
| BdTR2b | 7,664,367 | 6,172,380 | d | D |
| BdTR2b | 5,527,842 | 4,695,446 | e | W |
| BdTR2b | 4,076,062 | 3,451,682 | h | W |
| BdTR2b | 851,547 | 518,275 | d | W |
| BdTR2b | 3,216,653 | 2,214,486 | d | W |
| BdTR2g | 8,019,598 | 1,877,329 | e | D |
| BdTR2g | 7,613,459 | 2,207,658 | e | D |
| BdTR2g | 1,660,817 | 847,456 | d | D |
| BdTR2g | 5,672,244 | 5,066,239 | e | D |
| BdTR2g | 8,509,693 | 4,559,083 | g | W |
| BdTR2g | 6,190,328 | 4,640,646 | i | W |
| BdTR2g | 3,343,180 | 1,915,583 | d | W |
| BdTR2g | 5,106,301 | 3,857,120 | e | W |
| BdTR3c | 8,983,763 | 4,631,760 | e | D |
| BdTR3c | 5,783,183 | 3,418,183 | f | D |
| BdTR3c | 11,749,346 | 5,279,440 | c | D |
| BdTR3c | 3,845,280 | 2,589,362 | e | D |
| BdTR3c | 6,719,760 | 1,732,909 | e | W |
| BdTR3c | 10,425,380 | 5,456,807 | f | W |
| BdTR3c | 4,636,494 | 2,930,343 | c | W |
| BdTR3c | 1,864,046 | 955,753 | d | W |
| BdTR5i | 6,147,955 | 5,646,130 | g | D |
| BdTR5i | 6,615,513 | 6,141,256 | g | D |
| BdTR5i | 9,911,861 | 7,003,424 | c | D |
| BdTR5i | 5,101,192 | 4,085,464 | c | D |

| | | | | |
|---|---|---|---|---|
| BdTR5i | 10,175,612 | 3,244,779 | e | W |
| BdTR5i | 8,170,908 | 7,550,963 | g | W |
| BdTR5i | 21,415,184 | 16,945,841 | e | W |
| BdTR9k | 7,259,406 | 1,829,780 | e | D |
| BdTR9k | 4,411,548 | 930,976 | f | D |
| BdTR9k | 6,494,900 | 4,214,763 | a | D |
| BdTR9k | 4,568,566 | 3,358,484 | b | D |
| BdTR9k | 5,057,129 | 3,277,704 | e | W |
| BdTR9k | 438,159 | 290,059 | h | W |
| BdTR9k | 1,547,011 | 848,560 | d | W |
| BdTR9k | 1,551,245 | 818,788 | d | W |
| Bis1 | 10,011,946 | 3,363,014 | e | D |
| Bis1 | 6,751,170 | 5,619,119 | h | D |
| Bis1 | 12,089,934 | 8,812,063 | d | D |
| Bis1 | 1,718,896 | 1,275,179 | e | D |
| Bis1 | 2,480,717 | 1,920,222 | e | W |
| Bis1 | 3,737,209 | 2,567,122 | e | W |
| Bis1 | 1,997,874 | 1,089,328 | d | W |
| Bis1 | 3,721,297 | 2,853,085 | e | W |
| Kah1 | 3,626,318 | 2,577,194 | e | D |
| Kah1 | 7,985,749 | 6,827,694 | g | D |
| Kah1 | 7,086,549 | 3,085,245 | b | D |
| Kah1 | 3,852,565 | 2,846,075 | e | D |
| Kah1 | 4,006,900 | 3,496,340 | e | W |
| Kah1 | 5,137,740 | 3,398,223 | f | W |
| Kah1 | 2,338,856 | 1,504,107 | e | W |
| Kah1 | 8,404,640 | 6,208,590 | e | W |
| Kah5 | 10,194,420 | 4,100,570 | f | D |
| Kah5 | 2,280,340 | 1,927,109 | g | D |
| Kah5 | 6,221,252 | 3,284,009 | b | D |
| Kah5 | 3,528,651 | 3,016,609 | d | D |
| Kah5 | 5,593,350 | 1,861,012 | f | W |
| Kah5 | 5,239,891 | 4,620,182 | g | W |
| Kah5 | 2,577,573 | 1,856,909 | b | W |
| Kah5 | 4,445,867 | 3,605,558 | e | W |
| Koz1 | 3,505,791 | 2,654,375 | f | D |
| Koz1 | 3,753,891 | 2,934,115 | h | D |
| Koz1 | 2,186,858 | 989,478 | d | D |
| Koz1 | 3,372,454 | 2,456,387 | d | D |
| Koz1 | 6,847,592 | 5,801,961 | h | W |
| Koz1 | 1,347,983 | 440,861 | h | W |
| Koz1 | 4,448,408 | 3,044,017 | d | W |
| Koz1 | 2,776,127 | 2,116,334 | e | W |
| Koz3 | 4,185,182 | 3,591,463 | e | D |
| Koz3 | 3,676,421 | 2,971,293 | g | D |
| Koz3 | 2,038,664 | 1,698,405 | c | D |
| Koz3 | 8,083,293 | 6,657,015 | d | D |
| Koz3 | 9,774,753 | 4,793,443 | f | W |
| Koz3 | 3,739,971 | 3,045,871 | g | W |

| | | | | |
|---|---|---|---|---|
| Koz3 | 6,580,350 | 2,869,464 | c | W |
| Koz3 | 9,788,997 | 6,103,038 | c | W |
| Ron2 | 4,058,417 | 3,049,059 | e | D |
| Ron2 | 4,734,388 | 1,587,136 | f | D |
| Ron2 | 8,603,600 | 5,384,036 | b | D |
| Ron2 | 4,578,734 | 573,134 | f | W |
| Ron2 | 4,806,195 | 2,029,483 | f | W |
| Ron2 | 5,141,488 | 2,478,911 | a | W |
| Ron2 | 3,283,641 | 2,784,591 | e | W |

**Table S3.** Pearson's correlation among the module eigengenes (ME) for each Drought (**a**) and Water (**b**) network.

**(a)**

| Drought network | | gray | turquoise | blue | brown | yellow | green | red | black | pink | magenta |
|---|---|---|---|---|---|---|---|---|---|---|---|
| | | **ME0** | **ME1** | **ME2** | **ME3** | **ME4** | **ME5** | **ME6** | **ME7** | **ME8** | **ME9** |
| gray | **ME0** | 1 | 0.62 | 0.13 | 0.051 | 0.2 | -0.21 | -0.13 | -0.019 | 0.22 | 0.12 |
| turquoise | **ME1** | 0.62 | 1 | -0.56 | 0.18 | 0.33 | 0.018 | 0.069 | 0.11 | -0.18 | -0.41 |
| blue | **ME2** | 0.13 | -0.56 | 1 | -0.2 | -0.21 | -0.27 | -0.13 | -0.094 | 0.29 | 0.63 |
| brown | **ME3** | 0.051 | 0.18 | -0.2 | 1 | 0.25 | 0.1 | 0.52 | -0.028 | -0.19 | -0.2 |
| yellow | **ME4** | 0.2 | 0.33 | -0.21 | 0.25 | 1 | 0.095 | 0.1 | 0.1 | 0.052 | -0.01 |
| green | **ME5** | -0.21 | 0.018 | -0.27 | 0.1 | 0.095 | 1 | -0.065 | 0.016 | -0.24 | -0.44 |
| red | **ME6** | -0.13 | 0.069 | -0.13 | 0.52 | 0.1 | -0.065 | 1 | -0.013 | -0.34 | -0.27 |
| black | **ME7** | -0.019 | 0.11 | -0.094 | -0.028 | 0.1 | 0.016 | -0.013 | 1 | 0.022 | -0.19 |
| pink | **ME8** | 0.22 | -0.18 | 0.29 | -0.19 | 0.052 | -0.24 | -0.34 | 0.022 | 1 | 0.36 |
| magenta | **ME9** | 0.12 | -0.41 | 0.63 | -0.2 | -0.01 | -0.44 | -0.27 | -0.19 | 0.36 | 1 |

**(b)**

| Water network | | gray | turquoise | blue | brown | yellow | green | red | black | pink | magenta | purple | greenyellow | tan | salmon |
|---|---|---|---|---|---|---|---|---|---|---|---|---|---|---|---|
| | | ME0 | ME1 | ME2 | ME3 | ME4 | ME5 | ME6 | ME7 | ME8 | ME9 | ME10 | ME11 | ME12 | ME13 |
| gray | ME0 | 1 | -0.5 | 0.57 | -0.045 | -0.65 | 0.33 | 0.012 | 0.33 | 0.027 | 0.088 | 0.061 | 0.14 | 0.023 | -0.026 |
| turquoise | ME1 | -0.5 | 1 | -0.37 | -0.2 | -0.21 | -0.21 | -0.18 | -0.19 | -0.12 | -0.14 | -0.35 | -0.46 | -0.18 | -0.22 |
| blue | ME2 | 0.57 | -0.37 | 1 | 0.57 | -0.29 | 0.55 | 0.32 | 0.25 | 0.57 | -0.081 | -0.2 | -0.23 | 0.19 | -0.015 |
| brown | ME3 | -0.045 | -0.2 | 0.57 | 1 | 0.23 | 0.65 | 0.28 | 0.23 | 0.51 | 0.029 | -0.15 | -0.25 | 0.1 | 0.04 |
| yellow | ME4 | -0.65 | -0.21 | -0.29 | 0.23 | 1 | -0.11 | 0.068 | -0.15 | 0.0011 | -0.043 | 0.16 | 0.32 | 0.12 | 0.36 |
| green | ME5 | 0.33 | -0.21 | 0.55 | 0.65 | -0.11 | 1 | -0.085 | 0.47 | 0.16 | 0.18 | -0.27 | -0.0083 | -0.0087 | 0.061 |
| red | ME6 | 0.012 | -0.18 | 0.32 | 0.28 | 0.068 | -0.085 | 1 | 0.022 | 0.46 | -0.21 | 0.6 | -0.45 | 0.11 | -0.38 |
| black | ME7 | 0.33 | -0.19 | 0.25 | 0.23 | -0.15 | 0.47 | 0.022 | 1 | 0.098 | 0.29 | 0.011 | 0.068 | 0.039 | -0.076 |
| pink | ME8 | 0.027 | -0.12 | 0.57 | 0.51 | 0.0011 | 0.16 | 0.46 | 0.098 | 1 | -0.034 | -0.058 | -0.45 | 0.31 | -0.13 |
| magenta | ME9 | 0.088 | -0.14 | -0.081 | 0.029 | -0.043 | 0.18 | -0.21 | 0.29 | -0.034 | 1 | -0.1 | 0.22 | 0.049 | 0.074 |
| purple | ME10 | 0.061 | -0.35 | -0.2 | -0.15 | 0.16 | -0.27 | 0.6 | 0.011 | -0.058 | -0.1 | 1 | 0.01 | -0.02 | -0.22 |
| greenyellow | ME11 | 0.14 | -0.46 | -0.23 | -0.25 | 0.32 | -0.0083 | -0.45 | 0.068 | -0.45 | 0.22 | 0.01 | 1 | -0.061 | 0.53 |
| tan | ME12 | 0.023 | -0.18 | 0.19 | 0.1 | 0.12 | -0.0087 | 0.11 | 0.039 | 0.31 | 0.049 | -0.02 | -0.061 | 1 | 0.018 |
| salmon | ME13 | -0.026 | -0.22 | -0.015 | 0.04 | 0.36 | 0.061 | -0.38 | -0.076 | -0.13 | 0.074 | -0.22 | 0.53 | 0.018 | 1 |

**Table S4**. Identification of the plant Transcription factors (TF), transcriptional regulators (TR) and Protein Kinase from the protein sequences of the co-expressed genes clustered in the modules of the Drought (**a**) and Water (**b**) networks. The number of elements is indicated in parentheses.

**(a)**

| | | | **Protein kinases** | | | | | | | | |
|---|---|---|---|---|---|---|---|---|---|---|---|
| **Modules** | **TF** | **TR** | **Group AGC** | **Group CAMK** | **Group CK1** | **Group CMGC** | **Group Others** | **Group Plant-specific** | **Group RLK-Pelle** | **Group STE** | **Group TKL** |
| D1 | Alfin-like (1); AP2/ERF-ERF (7); B3 (7); B3-ARF (4); bHLH (10); bZIP (6); C2C2-CO-like (4); C2C2-Dof (6); C2C2-LSD (1); C2C2-YABBY (2); C2C2-GATA (3); C2H2 (6); C3H (7); CPP (1); CSD (1); DDT (5); E2F-DP (2); FAR1 (5); GARP-G2-like (8); GARP-ARR-B (2); GeBP (4); GRAS (1); GRF (1); HB-PHD (1); HB-other (2); HB-HD-ZIP (3); HB-BELL (4); HSF (1); MADS-MIKC (1); MADS-M-type (2); MYB (3); | ARID (1); AUX/IAA (6); Coactivator p15 (2); IWS1 (1); Jumonji (2); MBF1 (1); SET (18); SNF2 (8); SWI/SNF-BAF60b (3) | AGC_NDR (nuclear Dbf2-related kinases) (1) | CAMK_CDPK (calcium-dependent protein kinases) (5); CAMK_CAMKL-CHK1 (CAMK-Like, Checkpoint Kinase 1) (4); CAMK_CAMKL-LKB (CAMK-Like, liver kinase B1) (1) | CK1_CK1-Pl (Cell Kinase 1, Plant-specific) (2); CK1_CK1 (Cell Kinase 1) (2) | CMGC_DYRK-YAK (Dual-specificity Y (tyrosine) Regulated Kinase, A DYRK subfamily found only in Dictystelium and fungi) (1); CMGC_DYRK-PRP4 (Dual-specificity Y (tyrosine) Regulated Kinase, precursor mRNA processing 4) (6); | SCY1_SCYL1 (SCY1_SCYL1) (1); IRE1 (Inositol REquiring (named after yeast homolog). Also known as ERN (ER-to-nucleus signaling)) (1) | Group-Pl-4 (Group Plant-specific 4) (1); Group-Pl-3 (Group Plant-specific 3) (1) | RLK-Pelle_DLSV (receptor-like kinase/Pelle, DUF26,SD-1, LRR-VIII and VWA, a moss-specific new RLK subfamily) (2); RLK-Pelle_RLCK-XII-1 (receptor-like kinase/Pelle, Receptor Like Cytoplasmic Kinase-XII-1) (1); RLK-Pelle_LRR-II (receptor-like kinase/Pelle, leucine-rich repeat-II) (4); RLK-Pelle_CR4L (receptor-like kinase/Pelle, CRINKLY4-like) (1); RLK-Pelle_LRR-V | STE_STE11 (MAP3K (MAP kinase kinase kinase) genes, homologous to yeast Ste 11) (2); STE_STE7 (MAP2K (MAP kinase kinase) genes, homologous to yeast Ste 7) (4) | TKL_CTR1-DRK-2 (CTR1-DRK-2) (2); TKL-Pl-4 (Plant-specific 4) (5); TKL-Pl-1 (Plant-specific 1) (1) |

MYB-related (11); NAC (8); NF-YB (3); NF-YC (1); NF-YA (1); RWP-RK (2); S1Fa-like (3); Tify (4); TUB (2); VOZ (1)

CMGC_CLK (CDC-Like Kinase, involved in splicing) (1); CMGC_CDK-Pl (Cyclin Dependent Kinase, Plant-specific) (2); CMGC_CDK-CDK8 (Cyclin Dependent Kinase, Cyclin Dependent Kinase subfamily 8) (1); CMGC_GSK (Glycogen synthase 3 kinase) (2); CMGC_CDK-CRK7-CDK9 (Cyclin Dependent Kinase, cdc2-related kinase 7 - Cyclin-

(receptor-like kinase/Pelle, leucine-rich repeat-V) (1); RLK-Pelle_RLCK-V (receptor-like kinase/Pelle, Receptor Like Cytoplasmic Kinase-V) (1); RLK-Pelle_LRK10L-2 (receptor-like kinase/Pelle, LRK10-like kinase type 2) (1); RLK-Pelle_Extensin (receptor-like kinase/Pelle, Extensin) (4); RLK-Pelle_LRR-VIII-1 (receptor-like kinase/Pelle, leucine-rich repeat-VIII-1) (5); RLK-Pelle_RLCK-VIIa-2 (receptor-like kinase/Pelle, Receptor Like Cytoplasmic Kinase-VIIa-2) (3); RLK-

| | | | | | | | | | |
|---|---|---|---|---|---|---|---|---|---|
| | | | | | | dependent kinase 9) (5) | | Pelle_RLCK-IV (receptor-like kinase/Pelle, Receptor Like Cytoplasmic Kinase-IV) (2); RLK-Pelle_SD-2b (receptor-like kinase/Pelle, S Domain 2b) (1) | |
| D2 | B3-ARF (2); BES1 (2); bZIP (7); C2C2-Dof (1); C2C2-LSD (5); C2H2 (1); C3H (6); DBB (2); DBP (1); GARP-G2-like (5); HB-KNOX (2); HSF (2); LIM (1); LOB (1); MYB (2); MYB-related (4); NAC (2); NF-YC (3); NF-YB (1); RWP-RK (2); SBP (1); Trihelix (5); WRKY (5); zf-HD (1) | ARID (2); AUX/IAA (1); GNAT (7); mTERF (3); Others (5); PHD (2) | AGC_RSK-2 (Ribosomal S6 Kinases 2) (1) | | CK1_CK1 (Cell Kinase 1) (2) | CMGC_RCK (CMGC, ros cross-hybridizing kinase) (1); CMGC_CDK-CRK7-CDK9 (Cyclin Dependent Kinase, cdc2-related kinase 7 - Cyclin-dependent kinase 9) (1); CMGC_CDK-CCRK (Cyclin Dependent Kinase, Cell Cycle | | RLK-Pelle_LRR-III (receptor-like kinase/Pelle, leucine-rich repeat-III) (3); RLK-Pelle_LRR-XI-1 (receptor-like kinase/Pelle, leucine-rich repeat-XI-1) (2); RLK-Pelle_RLCK-IXa (receptor-like kinase/Pelle, Receptor Like Cytoplasmic Kinase-IXa) (2); RLK-Pelle_LRR-I-1 (receptor-like kinase/Pelle, leucine-rich repeat-I-1) (1) | STE_STE20-Pl (MAP4K (MAP kinase kinase kinase kinase) genes, homologous to yeast Ste 20, Plant-specific) (4) |

| | | | | | | | | | | | |
|---|---|---|---|---|---|---|---|---|---|---|---|
| | | | | | | Regulated Kinase) (1) | | | | | |
| D3 | AP2/ERF-RAV (1); AP2/ERF-AP2 (2); bHLH (1); CAMTA (5); GARP-G2-like (1); GRAS (1); MYB (2); NAC (6); PLATZ (1); WRKY (5) | GNAT (1); HMG (2); IWS1 (3) | AGC_NDR (nuclear Dbf2-related kinases) (2) | CAMK_CDPK (calcium-dependent protein kinases) (5); CAMK_OST1L (open stomata-like kinase) (3) | CK1_CK1 (Cell Kinase 1) (1) | CMGC_MAPK (Mitogen Activated Protein Kinase) (2); CMGC_CDK-CRK7-CDK9 (Cyclin Dependent Kinase, cdc2-related kinase 7 - Cyclin-dependent kinase 9) (3) | NAK (Numb-Associated Kinase, named after Drosophila member) (1); SCY1_SCYL2 (SCY1_SCYL2) (1) | | RLK-Pelle_LRR-II (receptor-like kinase/Pelle, leucine-rich repeat-II) (6); RLK-Pelle_LRR-Xa (receptor-like kinase/Pelle, leucine-rich repeat-Xa) (3); RLK-Pelle_RLCK-IV (receptor-like kinase/Pelle, Receptor Like Cytoplasmic Kinase-IV) (4); RLK-Pelle_L-LEC (receptor-like kinase/Pelle, L-type lectin) (8); RLK-Pelle_WAK (receptor-like kinase/Pelle, Wall Associated Kinase) (4); RLK-Pelle_RLCK-Os (receptor-like kinase/Pelle, Receptor Like Cytoplasmic | STE_STE7 (MAP2K (MAP kinase kinase) genes, homologous to yeast Ste 7) (3) | TKL-Pl-5 (Plant-specific 5) (1); TKL-Pl-4 (Plant-specific 4) (1) |

Kinase-Os) (3);
RLK-Pelle_LRR-XII-1 (receptor-like kinase/Pelle, leucine-rich repeat-XII-1) (3);
RLK-Pelle_DLSV (receptor-like kinase/Pelle, DUF26,SD-1, LRR-VIII and VWA, a moss-specific new RLK subfamily) (10);
RLK-Pelle_LRR-Xb-1 (receptor-like kinase/Pelle, leucine-rich repeat-Xb-1) (1);
RLK-Pelle_SD-2b (receptor-like kinase/Pelle, S Domain 2b) (5);
RLK-Pelle_RLCK-VIIa-2 (receptor-like kinase/Pelle, Receptor Like Cytoplasmic Kinase-VIIa-2) (7); RLK-Pelle_LysM (receptor-like kinase/Pelle, LysM Domain-

containing Kinase) (3); RLK-Pelle_Extensin (receptor-like kinase/Pelle, Extensin) (1); RLK-Pelle_LRR-VIII-1 (receptor-like kinase/Pelle, leucine-rich repeat-VIII-1) (8); RLK-Pelle_LRK10L-2 (receptor-like kinase/Pelle, LRK10-like kinase type 2) (3); RLK-Pelle_LRR-VI-2 (receptor-like kinase/Pelle, leucine-rich repeat-VI-2) (5); RLK-Pelle_RLCK-VI (receptor-like kinase/Pelle, Receptor Like Cytoplasmic Kinase-VI) (1); RLK-Pelle_LRR-XIV (receptor-like kinase/Pelle, leucine-rich repeat-XIV) (2);

| | | | | | | | | |
|---|---|---|---|---|---|---|---|---|
| | | | | | | RLK-Pelle_LRR-XI-2 (receptor-like kinase/Pelle, leucine-rich repeat-XI-2) (1); RLK-Pelle_RLCK-IXb (receptor-like kinase/Pelle, Receptor Like Cytoplasmic Kinase-IXb) (2); RLK-Pelle_WAK_LRK10L-1 (receptor-like kinase/Pelle, Wall Associated Kinase, LRK10-like kinase type 1) (3); RLK-Pelle_LRR-XI-1 (receptor-like kinase/Pelle, leucine-rich repeat-XI-1) (2) | | |
| D4 | AP2/ERF-ERF (4); BES1 (1); bHLH (8); bZIP (1); C2C2-LSD (2); C2C2-GATA (1); C2H2 (6); CAMTA (1); GRAS (2); MYB (2); NAC (10); WRKY (9) | | CAMK_CDPK (calcium-dependent protein kinases) (3); CAMK_CAMKL-CHK1 (CAMK-Like, Checkpoint Kinase 1) (3) | CMGC_MAPK (Mitogen Activated Protein Kinase) (2) | | RLK-Pelle_LRR-III (receptor-like kinase/Pelle, leucine-rich repeat-III) (1); RLK-Pelle_DLSV (receptor-like kinase/Pelle, DUF26,SD-1, LRR-VIII and | STE_STE11 (MAP3K (MAP kinase kinase kinase) genes, homologous to yeast Ste 11) (2) | TKL-Pl-5 (Plant-specific 5) (3) |

VWA, a moss-specific new RLK subfamily) (5); RLK-Pelle_LysM (receptor-like kinase/Pelle, LysM Domain-containing Kinase) (1); RLK-Pelle_RLCK-VIIa-2 (receptor-like kinase/Pelle, Receptor Like Cytoplasmic Kinase-VIIa-2) (3); RLK-Pelle_RLCK-VIII (receptor-like kinase/Pelle, Receptor Like Cytoplasmic Kinase-VIII) (4); RLK-Pelle_LRR-IX (receptor-like kinase/Pelle, leucine-rich repeat-IX) (1); RLK-Pelle_RKF3 (receptor-like kinase/Pelle, Receptor-like kinase in flowers 3) (1); RLK-Pelle_PERK-1

| | | | | | | |
|---|---|---|---|---|---|---|
| | | | | | | (receptor-like kinase/Pelle, Plant External Response Like Kinase 1) (3) |
| D5 | | MBF1 (1); Others (1); PHD (1); TRAF (2) | | | | |
| D6 | bHLH (1); C2H2 (1); MYB (3); NAC (1) | IWS1 (1); mTERF (1) | | CMGC_MAPK (Mitogen Activated Protein Kinase) (5) | | |
| D7 | C2C2-CO-like (1); C2H2 (3); EIL (2) | AUX/IAA (1) | | | | |
| D8 | SNF2 (9) | | | | | |
| D9 | HB-KNOX (3) | | | | | |

**(b)**

| Modules | TF | TR | Protein kinases: Group AGC | Group CAMK | Group CK1 | Group CMGC | Group Others | Group Plant-specific | Group RLK-Pelle | Group STE | Group TKL |
|---|---|---|---|---|---|---|---|---|---|---|---|
| W1 | Alfin-like (3); AP2/ERF-ERF (1); B3-ARF (1); bHLH (4); bZIP (6); C2C2-GATA (5); C2C2-CO-like (2); C2C2-Dof (7); C2H2 (5); C3H (10); CAMTA (1); DDT (2); FAR1 (4); GARP-G2-like (1); GARP-ARR-B (4); GeBP (4); GRAS (3); GRF (1); HB-KNOX (3); HB-BELL (5); HSF (1); MADS-MIKC (1); MYB (3); MYB-related (17); NAC (2); NF-YA (4); NF-YB (1); NF-X1 (1); PLATZ (5); Trihelix (2); | ARID (4); AUX/IAA (1); Coactivator p15 (1); GNAT (10); IWS1 (3); Jumonji (3); LUG (1); MED6 (1); MED7 (3); mTERF (4); Others (4); PHD (3); RB (1); Rcd1-like (1); SET (3); SNF2 (6); SWI/SNF-BAF60b (3) | AGC_RSK-2 (Ribosomal S6 Kinases 2) (2); AGC_NDR (nuclear Dbf2-related kinases) (2) | CAMK_CAMKL-LKB (CAMK-Like, liver kinase B1) (1); CAMK_CDPK (calcium-dependent protein kinases) (8); CAMK_CAMKL-CHK1 (CAMK-Like, Checkpoint Kinase 1) (3); CAMK_OST1L (open stomata-like kinase) (2); CAMK_AMPK (AMP-activated protein kinase) (2) | CK1_CK1 (Cell Kinase 1) (1) | CMGC_DYRK-PRP4 (Dual-specificity Y (tyrosine) Regulated Kinase, precursor mRNA processing 4) (6); CMGC_RCK (CMGC, ros cross-hybridizing kinase) (1); CMGC_CDK-CRK7-CDK9 (Cyclin Dependent Kinase, cdc2-related kinase 7 - Cyclin-dependent kinase 9) (1); CMGC_GSK (Glycogen synthase 3 | NAK (Numb-Associated Kinase, named after Drosophila member) (1) | | RLK-Pelle_LRR-XI-1 (receptor-like kinase/Pelle, leucine-rich repeat-XI-1) (1); RLK-Pelle_LRR-VI-2 (receptor-like kinase/Pelle, leucine-rich repeat-VI-2) (1); RLK-Pelle_LRR-XII-1 (receptor-like kinase/Pelle, leucine-rich repeat-XII-1) (1); RLK-Pelle_RLCK-VIIa-2 (receptor-like kinase/Pelle, Receptor Like Cytoplasmic Kinase-VIIa-2) (3); RLK-Pelle_DLSV | STE_STE11 (MAP3K (MAP kinase kinase kinase) genes, homologous to yeast Ste 11) (1); STE_STE20-Fray (MAP4K (MAP kinase kinase kinase kinase) genes, homologous to yeast Ste 20, Fray (Named based on Drosophila family | TKL_CTR1-DRK-2 (CTR1-DRK-2) (1); TKL-Pl-1 (Plant-specific 1) (1); TKL_CTR1-DRK-1 (CTR1-DRK-1) (2); TKL-Pl-5 (Plant-specific 5) (4) |

| | | | | |
|---|---|---|---|---|
| TUB (2); WRKY (1); zf-HD (1) | | kinase) (2); CMGC_CLK (CDC-Like Kinase, involved in splicing) (1); CMGC_CDK-CDK7 (Cyclin Dependent Kinase, Cyclin Dependent Kinase subfamily 7) (1); CMGC_CK2 (Cell Kinase 2) (1); CMGC_CDK-CCRK (Cyclin Dependent Kinase, Cell Cycle Regulated Kinase) (1); CMGC_MAPK (Mitogen Activated Protein Kinase) (1) | (receptor-like kinase/Pelle, DUF26,SD-1, LRR-VIII and VWA, a moss-specific new RLK subfamily) (5); RLK-Pelle_LRR-V (receptor-like kinase/Pelle, leucine-rich repeat-V) (1); RLK-Pelle_LRR-XIIIa (receptor-like kinase/Pelle, leucine-rich repeat-XIIIa) (1); RLK-Pelle_CrRLK1L-1 (receptor-like kinase/Pelle, Catharanthus roseus RLK1-like) (2); RLK-Pelle_RLCK-IXa (receptor-like kinase/Pelle, Receptor Like Cytoplasmic Kinase-IXa) (2); RLK-Pelle_RLCK-VIIa-1 | member)) (1) |

| | | | | | | | | | |
|---|---|---|---|---|---|---|---|---|---|
| | | | | | | | (receptor-like kinase/Pelle, Receptor Like Cytoplasmic Kinase-VIIa-1) (2) | | |
| W2 | AP2/ERF-ERF (2); AP2/ERF-AP2 (2); bHLH (4); C2H2 (4); C3H (5); CSD (1); GeBP (2); MYB-related (1); NAC (1) | AUX/IAA (1); GNAT (1); HMG (1); Jumonji (2); mTERF (1); PHD (1); SET (2); SOH1 (1); SWI/SNF-SWI3 (1); SWI/SNF-BAF60b (1) | | | CMGC_CDK-CRK7-CDK9 (Cyclin Dependent Kinase, cdc2-related kinase 7 - Cyclin-dependent kinase 9) (1) | | RLK-Pelle_RLCK-V (receptor-like kinase/Pelle, Receptor Like Cytoplasmic Kinase-V) (1); RLK-Pelle_LRR-XIIIa (receptor-like kinase/Pelle, leucine-rich repeat-XIIIa) (3) | STE_STE20-Pl (MAP4K (MAP kinase kinase kinase kinase) genes, homologous to yeast Ste 20, Plant-specific) (4) | TKL-Pl-6 (Plant-specific 6) (7) |

| | | | | | | | | | | | |
|---|---|---|---|---|---|---|---|---|---|---|---|
| W3 | Alfin-like (1); AP2/ERF-ERF (1); B3-ARF (3); BES1 (1); bHLH (1); C2H2 (2); CAMTA (4); GRAS (1); S1Fa-like (3); Tify (1); WRKY (1) | GNAT (1); IWS1 (1); TRAF (3) | | CAMK_CAMKL-CHK1 (CAMK-Like, Checkpoint Kinase 1) (2); CAMK_CDPK (calcium-dependent protein kinases) (1) | | CMGC_MAPK (Mitogen Activated Protein Kinase) (7); CMGC_CDK-CRK7-CDK9 (Cyclin Dependent Kinase, cdc2-related kinase 7 - Cyclin-dependent kinase 9) (1) | NAK (Numb-Associated Kinase, named after Drosophila member) (1); SCY1_SCYL2 (SCY1_SCYL2) (1) | | RLK-Pelle_LRK10L-2 (receptor-like kinase/Pelle, LRK10-like kinase type 2) (3); RLK-Pelle_DLSV (receptor-like kinase/Pelle, DUF26,SD-1, LRR-VIII and VWA, a moss-specific new RLK subfamily) (4); RLK-Pelle_LRR-XI-1 (receptor-like kinase/Pelle, leucine-rich repeat-XI-1) (3); RLK-Pelle_LRR-VII-2 (receptor-like kinase/Pelle, leucine-rich repeat-VII-2) (1); RLK-Pelle_LRR-II (receptor-like kinase/Pelle, leucine-rich repeat-II) (1); RLK-Pelle_LysM | STE_STE7 (MAP2K (MAP kinase kinase) genes, homologous to yeast Ste 7) (3) | TKL-Pl-5 (Plant-specific 5) (1) |

(receptor-like
kinase/Pelle,
LysM Domain-
containing
Kinase) (1); RLK-
Pelle_LRR-III
(receptor-like
kinase/Pelle,
leucine-rich
repeat-III) (1);
RLK-Pelle_LRR-
VI-2 (receptor-
like
kinase/Pelle,
leucine-rich
repeat-VI-2) (5);
RLK-Pelle_SD-
2b (receptor-
like
kinase/Pelle, S
Domain 2b) (1);
RLK-Pelle_RLCK-
IXb (receptor-
like
kinase/Pelle,
Receptor Like
Cytoplasmic
Kinase-IXb) (2);
RLK-Pelle_LRR-
VIII-1 (receptor-
like
kinase/Pelle,
leucine-rich

| | | | | | | | | | | |
|---|---|---|---|---|---|---|---|---|---|---|
| | | | | | | | | repeat-VIII-1) (1) | | |
| W4 | B3-ARF (13); bHLH (1); bZIP (3); C2C2-GATA (1); C2C2-Dof (6); C2H2 (6); C3H (1); CPP (1); DDT (3); FAR1 (1); GeBP (1); GRAS (1); MADS-MIKC (1); MYB-related (3); NAC (4); NF-YC (1); RWP-RK (1); Tify (3); Trihelix (6); zf-HD (1) | AUX/IAA (1); GNAT (1); IWS1 (3); Jumonji (2); mTERF (1); Others (1); SET (17); SNF2 (9); SWI/SNF-BAF60b (2) | AGC_NDR (nuclear Dbf2-related kinases) (1) | CAMK_OST1L (open stomata-like kinase) (2); CAMK_CDPK (calcium-dependent protein kinases) (5); CAMK_AMPK (AMP-activated protein kinase) (1) | | CMGC_SRPK (SR Protein Kinase; phosphorylates SR splicing factors) (2); CMGC_CDK-Pl (Cyclin Dependent Kinase, Plant-specific) (2); CMGC_RCK (CMGC, ros cross-hybridizing kinase) (2); CMGC_MAPK (Mitogen Activated Protein Kinase) (1) | | RLK-Pelle_LRR-IX (receptor-like kinase/Pelle, leucine-rich repeat-IX) (1); RLK-Pelle_LysM (receptor-like kinase/Pelle, LysM Domain-containing Kinase) (1); RLK-Pelle_LRR-II (receptor-like kinase/Pelle, leucine-rich repeat-II) (3); RLK-Pelle_RLCK-VIII (receptor-like kinase/Pelle, Receptor Like Cytoplasmic Kinase-VIII) (5); RLK-Pelle_LRR-XI-1 (receptor- | STE_STE7 (MAP2K (MAP kinase kinase) genes, homologous to yeast Ste 7) (2) | TKL_CTR1-DRK-2 (CTR1-DRK-2) (1); TKL-Pl-4 (Plant-specific 4) (2) |

| | | | | | | | | | |
|---|---|---|---|---|---|---|---|---|---|
| | | | | | | | like kinase/Pelle, leucine-rich repeat-XI-1) (1); RLK-Pelle_RLCK-VIIa-2 (receptor-like kinase/Pelle, Receptor Like Cytoplasmic Kinase-VIIa-2) (1) | | |
| W5 | Alfin-like (1); AP2/ERF-RAV (1); B3-ARF (1); bHLH (1); bZIP (4); C2H2 (1); CAMTA (1); FAR1 (4); GRAS (2); HSF (1); LIM (1); MYB (1); NAC (4); NF-YC (1); WRKY (3) | GNAT (1); HMG (1); SWI/SNF-BAF60b (2); TRAF (1) | | CAMK_CDPK (calcium-dependent protein kinases) (7); CAMK_CAMKL-CHK1 (CAMK-Like, Checkpoint Kinase 1) (2) | CMGC_MAPK (Mitogen Activated Protein Kinase) (1); CMGC_CDK-CRK7-CDK9 (Cyclin Dependent Kinase, cdc2-related kinase 7 - Cyclin-dependent kinase 9) (1) | | RLK-Pelle_LRR-II (receptor-like kinase/Pelle, leucine-rich repeat-II) (6); RLK-Pelle_LRR-III (receptor-like kinase/Pelle, leucine-rich repeat-III) (1); RLK-Pelle_LRR-Xa (receptor-like kinase/Pelle, leucine-rich repeat-Xa) (3); RLK-Pelle_RLCK-IV (receptor-like kinase/Pelle, Receptor Like Cytoplasmic Kinase-IV) (4); | STE_STE11 (MAP3K (MAP kinase kinase kinase) genes, homologous to yeast Ste 11) (1); STE_STE7 (MAP2K (MAP kinase kinase) genes, homologous to yeast Ste 7) (3) | TKL-Pl-4 (Plant-specific 4) (2); TKL_CTR1-DRK-2 (CTR1-DRK-2) (2) |

RLK-Pelle_L-LEC (receptor-like kinase/Pelle, L-type lectin) (6); RLK-Pelle_WAK (receptor-like kinase/Pelle, Wall Associated Kinase) (3); RLK-Pelle_DLSV (receptor-like kinase/Pelle, DUF26,SD-1, LRR-VIII and VWA, a moss-specific new RLK subfamily) (13); RLK-Pelle_LRR-Xb-1 (receptor-like kinase/Pelle, leucine-rich repeat-Xb-1) (1); RLK-Pelle_LRR-VI-2 (receptor-like kinase/Pelle, leucine-rich repeat-VI-2) (2); RLK-Pelle_RLCK-VIIa-2 (receptor-like kinase/Pelle, Receptor Like

Cytoplasmic
Kinase-VIIa-2)
(6); RLK-
Pelle_SD-2b
(receptor-like
kinase/Pelle, S
Domain 2b) (3);
RLK-Pelle_LysM
(receptor-like
kinase/Pelle,
LysM Domain-
containing
Kinase) (3); RLK-
Pelle_LRK10L-2
(receptor-like
kinase/Pelle,
LRK10-like
kinase type 2)
(1); RLK-
Pelle_LRR-XIV
(receptor-like
kinase/Pelle,
leucine-rich
repeat-XIV) (1);
RLK-Pelle_LRR-
XI-2 (receptor-
like
kinase/Pelle,
leucine-rich
repeat-XI-2) (1);
RLK-
Pelle_WAK_LRK
10L-1 (receptor-
like

| | | | | |
|---|---|---|---|---|
| | | | | kinase/Pelle, Wall Associated Kinase, LRK10-like kinase type 1) (2); RLK-Pelle_LRR-I-1 (receptor-like kinase/Pelle, leucine-rich repeat-I-1) (1) |
| W6 | AP2/ERF-ERF (2); bHLH (5); C2H2 (2); HB-HD-ZIP (3); MYB (4); MYB-related (1); NAC (1); Tify (9); Whirly (1) | Others (4); TRAF (1) | CAMK_AMPK (AMP-activated protein kinase) (3); CAMK_CDPK (calcium-dependent protein kinases) (1) | RLK-Pelle_RLCK-V (receptor-like kinase/Pelle, Receptor Like Cytoplasmic Kinase-V) (3); RLK-Pelle_RLCK-VI (receptor-like kinase/Pelle, Receptor Like Cytoplasmic Kinase-VI) (3); RLK-Pelle_LRR-II (receptor-like kinase/Pelle, leucine-rich repeat-II) (4); RLK-Pelle_RLCK-VIIb (receptor-like |

kinase/Pelle,
Receptor Like
Cytoplasmic
Kinase-VIIb) (2);
RLK-Pelle_RLCK-
XIII (receptor-
like
kinase/Pelle,
Receptor Like
Cytoplasmic
Kinase-XIII) (1);
RLK-Pelle_RLCK-
VIII (receptor-
like
kinase/Pelle,
Receptor Like
Cytoplasmic
Kinase-VIII) (1);
RLK-Pelle_SD-
2b (receptor-
like
kinase/Pelle, S
Domain 2b) (3);
RLK-Pelle_RLCK-
VIIa-2
(receptor-like
kinase/Pelle,
Receptor Like
Cytoplasmic
Kinase-VIIa-2)
(4); RLK-
Pelle_LRR-III
(receptor-like
kinase/Pelle,

| | | | | | | |
|---|---|---|---|---|---|---|
| | | | | | | leucine-rich repeat-III) (1); RLK-Pelle_WAK_LRK10L-1 (receptor-like kinase/Pelle, Wall Associated Kinase, LRK10-like kinase type 1) (1) |
| W7 | AP2/ERF-ERF (5); DBP (1); GRAS (1); MYB-related (1); NAC (5); Trihelix (2) | Others (1) | CAMK_CDPK (calcium-dependent protein kinases) (3); CAMK_CAMKL-CHK1 (CAMK-Like, Checkpoint Kinase 1) (8); CAMK_OST1L (open stomata-like kinase) (1) | CMGC_CDK-PITSLRE (Cyclin Dependent Kinase, PITSLRE) (4) | | RLK-Pelle_RLCK-Os (receptor-like kinase/Pelle, Receptor Like Cytoplasmic Kinase-Os) (3); RLK-Pelle_RLCK-VIIa-2 (receptor-like kinase/Pelle, Receptor Like Cytoplasmic Kinase-VIIa-2) (2); RLK-Pelle_WAK (receptor-like kinase/Pelle, Wall Associated Kinase) (2); RLK-Pelle_LRR-VIII-1 (receptor-like kinase/Pelle, |

| | | | | | | | | |
|---|---|---|---|---|---|---|---|---|
| | | | | | | | leucine-rich repeat-VIII-1) (3); RLK-Pelle_RKF3 (receptor-like kinase/Pelle, Receptor-like kinase in flowers 3) (1); RLK-Pelle_RLCK-II (receptor-like kinase/Pelle, Receptor Like Cytoplasmic Kinase-II) (2); RLK-Pelle_DLSV (receptor-like kinase/Pelle, DUF26,SD-1, LRR-VIII and VWA, a moss-specific new RLK subfamily) (3) | |
| W8 | AP2/ERF-ERF (2); BES1 (1); bHLH (6); bZIP (1); C2C2-Dof (1); C2H2 (1); C3H (2); GRAS (3); HB-HD-ZIP (2); HSF (1); MYB (7); MYB-related | Others (4) | | CAMK_OST1L (open stomata-like kinase) (2); CAMK_CDPK (calcium-dependent protein kinases) (1) | CMGC_MAPK (Mitogen Activated Protein Kinase) (2) | | RLK-Pelle_PERK-2 (receptor-like kinase/Pelle, Plant External Response Like Kinase 2) (1); RLK-Pelle_RLCK-XII-1 (receptor-like | TKL-Pl-6 (Plant-specific 6) (3) |

| | | | | | | | | |
|---|---|---|---|---|---|---|---|---|
| | (2); NAC (3); WRKY (1) | | | | | | kinase/Pelle, Receptor Like Cytoplasmic Kinase-XII-1) (1); RLK-Pelle_RLCK-VIIa-2 (receptor-like kinase/Pelle, Receptor Like Cytoplasmic Kinase-VIIa-2) (1); RLK-Pelle_LRR-III (receptor-like kinase/Pelle, leucine-rich repeat-III) (1); RLK-Pelle_LRR-V (receptor-like kinase/Pelle, leucine-rich repeat-V) (3); RLK-Pelle_L-LEC (receptor-like kinase/Pelle, L-type lectin) (1) | |
| W9 | AP2/ERF-ERF (6); B3-ARF (3); BES1 (1); bHLH (6); bZIP (2); C2C2-GATA (1); C2C2-Dof (1); | AUX/IAA (1) | | CAMK_CDPK (calcium-dependent protein kinases) (6); CAMK_OST1L (open | CMGC_MAPK (Mitogen Activated Protein Kinase) (2) | | RLK-Pelle_DLSV (receptor-like kinase/Pelle, DUF26,SD-1, LRR-VIII and VWA, a moss-specific new | STE_STE7 (MAP2K (MAP kinase kinase) genes, homologo |

| | | | | | | |
|---|---|---|---|---|---|---|
| | C2C2-CO-like (1); C2H2 (4); DBP (2); GRAS (2); HB-HD-ZIP (2); MYB (3); NAC (11); Tify (2); WRKY (12) | | stomata-like kinase) (3); CAMK_CAMKL-CHK1 (CAMK-Like, Checkpoint Kinase 1) (2) | | RLK subfamily) (2); RLK-Pelle_LysM (receptor-like kinase/Pelle, LysM Domain-containing Kinase) (1); RLK-Pelle_RLCK-VIII (receptor-like kinase/Pelle, Receptor Like Cytoplasmic Kinase-VIII) (4); RLK-Pelle_RLCK-VIIa-1 (receptor-like kinase/Pelle, Receptor Like Cytoplasmic Kinase-VIIa-1) (1); RLK-Pelle_PERK-1 (receptor-like kinase/Pelle, Plant External Response Like Kinase 1) (3) | us to yeast Ste 7) (1); STE_STE11 (MAP3K (MAP kinase kinase kinase) genes, homologous to yeast Ste 11) (2) |
| W10 | C3H (4); MYB (1); MYB-related (1); NF-YB (1) | SET (1) | | CMGC_MAPK (Mitogen Activated Protein Kinase) (2) | | |

| | | | | | | | | |
|---|---|---|---|---|---|---|---|---|
| W11 | bZIP (1); MYB-related (8); OFP (1); RWP-RK (2) | | AGC_RSK-2 (Ribosomal S6 Kinases 2) (3); AGC_PDK1 (phosphoinositide-dependent protein kinase 1) (3) | | CMGC_GSK (Glycogen synthase 3 kinase) (1); CMGC_MAPK (Mitogen Activated Protein Kinase) (1) | | RLK-Pelle_SD-2b (receptor-like kinase/Pelle, S Domain 2b) (1); RLK-Pelle_DLSV (receptor-like kinase/Pelle, DUF26,SD-1, LRR-VIII and VWA, a moss-specific new RLK subfamily) (1) | |
| W12 | EIL (2); HB-HD-ZIP (1); OFP (1) | AUX/IAA (1); HMG (2); IWS1 (4); mTERF (1) | | | | | RLK-Pelle_DLSV (receptor-like kinase/Pelle, DUF26,SD-1, LRR-VIII and VWA, a moss-specific new RLK subfamily) (3); RLK-Pelle_RLCK-VI (receptor-like kinase/Pelle, Receptor Like Cytoplasmic Kinase-VI) (3) | TKL-Pl-4 (Plant-specific 4) (3) |
| W13 | | | | | | | | TKL-Pl-4 (Plant-specific 4) (1); TKL-Pl-6 |

(Plant-
specific
6) (7)

**Table S5.** Number and proportion of the DE isoforms according to the up- and downregulation in the drought condition and their assignation to the Drought (**a**) or Water (**b**) modules.

**(a)**

| Drought modules | DE isoforms per module | DE isoforms upregulated in drought condition per module | | DE isoforms downregulated in drought condition per module | | ratio (rounded) upregulated:downregulated in drought condition | |
|---|---|---|---|---|---|---|---|
| | | # | % | # | % | | |
| D0 | 2930 | 2369 | 80.9 | 561 | 19.1 | ↑ | 4:1 |
| D1 | 878 | 767 | 87.4 | 111 | 12.6 | ↑ | 7:1 |
| D2 | 622 | 139 | 22.3 | 483 | 77.7 | ↓ | 1:3.5 |
| D3 | 220 | 89 | 40.5 | 131 | 59.5 | ↓ | 1:1.5 |
| D4 | 84 | 54 | 64.3 | 30 | 35.7 | ↑ | 2:1 |
| D5 | 116 | 107 | 92.2 | 9 | 7.8 | ↑ | 12:1 |
| D6 | 35 | 34 | 97.1 | 1 | 2.9 | ↑ | 34:1 |
| D7 | 24 | 22 | 91.7 | 2 | 8.3 | ↑ | 11:1 |
| D8 | 4 | 2 | 50.0 | 2 | 50.0 | ↔ | 1:1 |
| D9 | 28 | 8 | 28.6 | 20 | 71.4 | ↓ | 1:2.5 |
| **Total Drougth network** | **4941** | **3591** | **72.7** | **1350** | **27.3** | **↑** | **2.5:1** |
| **Total proper Drought network (excluding zero module)** | **2011** | **1222** | **60.8** | **789** | **39.2** | **↑** | **1.5:1** |

**(b)**

| Water modules | DE isoforms per module | DE isoforms upregulated in water condition per module | | DE isoforms downregulated in water condition per module | | ratio (rounded) upregulated:downregulated in drought condition | |
|---|---|---|---|---|---|---|---|
| | | # | % | # | % | | |
| W0 | 2342 | 1618 | 69.1 | 724 | 30.9 | ↑ | 2:1 |
| W1 | 446 | 304 | 68.2 | 142 | 31.8 | ↑ | 2:1 |
| W2 | 600 | 579 | 96.5 | 21 | 3.5 | ↑ | 27.5:1 |
| W3 | 328 | 297 | 90.5 | 31 | 9.5 | ↑ | 9.5:1 |
| W4 | 138 | 54 | 39.1 | 84 | 60.9 | ↓ | 1:1.5 |
| W5 | 183 | 102 | 55.7 | 81 | 44.3 | ↑ | 1.5:1 |
| W6 | 283 | 264 | 93.3 | 19 | 6.7 | ↑ | 14:1 |
| W7 | 108 | 77 | 71.3 | 31 | 28.7 | ↑ | 2.5:1 |
| W8 | 212 | 211 | 99.5 | 1 | 0.5 | ↑ | 211:1 |
| W9 | 81 | 47 | 58.0 | 34 | 42.0 | ↑ | 1.5:1 |
| W10 | 154 | 17 | 11.0 | 137 | 89.0 | ↓ | 1:8 |
| W11 | 46 | 6 | 13.0 | 40 | 87.0 | ↓ | 1:6.5 |
| W12 | 19 | 15 | 78.9 | 4 | 21.1 | ↑ | 4:1 |
| W13 | 1 | 0 | 0.0 | 1 | 100.0 | ↓ | 0:1 |
| **Total Water network** | **4941** | **3591** | **72.7** | **1350** | **27.3** | **↑** | **2.5:1** |
| **Total proper Water network (excluding zero module)** | **2599** | **1973** | **75.91381301** | **626** | **24.086187** | **↑** | **3:1** |

**Table S6**. Number and proportion of the DE hub nodes (isoforms) per Drought (**a**) and Water (**b**) modules.

**(a)**

| | Hub nodes per Drought modules | | | | | | | | | |
|---|---|---|---|---|---|---|---|---|---|---|
| **ID modules** | **D1** | **D2** | **D3** | **D4** | **D5** | **D6** | **D7** | **D8** | **D9** | **TOTAL** |
| **Total hub nodes per Drought modules** | 72 | 100 | 114 | 58 | 34 | 24 | 11 | 16 | 11 | 440 |
| **DE hub nodes per Drought module** | 48 (66.7%) | 98 (98.0%) | 48 (42.1%) | 16 (27.6%) | 22 (64.7%) | 10 (41.7%) | 5 (45.5%) | 1 (6.25%) | 11 (100%) | 259 (58.9%) |

**(b)**

| | Hub nodes per Water modules | | | | | | | | | | | | | |
|---|---|---|---|---|---|---|---|---|---|---|---|---|---|---|
| **ID modules** | **W1** | **W2** | **W3** | **W4** | **W5** | **W6** | **W7** | **W8** | **W9** | **W10** | **W11** | **W12** | **W13** | **TOTAL** |
| **Total hub nodes per Water module** | 321 | 45 | 67 | 44 | 104 | 65 | 81 | 40 | 70 | 37 | 18 | 9 | 10 | 911 |
| **DE hub nodes per Water module** | 72 (22.4%) | 43 (95.6%) | 22 (32.8%) | 4 (9.1%) | 46 (44.2%) | 59 (90.8%) | 24 (29.6%) | 38 (95.0%) | 17 (24.3%) | 36 (97.3%) | 18 (100%) | 1 (11.1%) | 0 (0%) | 380 (41.7%) |

**Table S7.** Summary annotation of the 50 isoforms of the differentially expressed genes (DEG; 25 up- 25 down-regulated) under drought (D) and water (W) conditions. Isoform (*B. distachyon* 314 v3.1 reference transcriptome); regulation (regulation under drought condition compared to water condition); occupancy (core (33 accessions), soft-core (31-32 accessions) and shell (≤30 accessions)); matched Drought modules and matched Water modules (numeric identification of the module where the DE isoforms were assigned); DE hub node (DE gene detected such as hub gene in the Drought (D) or Water (D) modules); Pfam, KEGG/ec, GO annotations; arabi-defline and rice-defline (annotation in Arabidopsis and rice). The dashes indicate that no results were retrieved.

| DE isoforms | regulation | occupancy | matched Drought modules | matched Water modules | DE hub node (D module) | DE hub node (W module) | Pfam | KEGG/ec | GO | arabi-defline | rice-defline |
|---|---|---|---|---|---|---|---|---|---|---|---|
| Bradi4g25750.1 | up | soft-core (32) | D0 | W6 | | | PF00234 | - | - | lipid transfer protein 12 | LTPL11 - Protease inhibitor/seed storage/LTP family protein precursor, expressed |
| Bradi4g44410.1 | up | core (33) | D0 | W6 | | hub (W6) | PF00234 | - | - | lipid transfer protein 1 | LTPL10 - Protease inhibitor/seed storage/LTP family protein precursor, expressed |
| Bradi1g70390.1 | up | core (33) | D1 | W8 | | | PF05042 | 1.11.2.3 | - | Caleosin-related family protein | caleosin related protein, putative, expressed |
| Bradi4g24650.1 | up | core (33) | D2 | W6 | | | PF02496 | - | GO:0006950 | - | abscisic stress-ripening, putative, expressed |
| Bradi2g56260.1 | up | core (33) | D0 | W10 | | | PF00504 | - | - | Chlorophyll A-B binding family protein | chlorophyll A-B binding protein, |

| | | | | | | | | | | | |
|---|---|---|---|---|---|---|---|---|---|---|---|
| | | | | | | | | | | | putative, expressed |
| Bradi3g00910.1 | up | Shell (30) | D0 | W0 | | | PF08244,PF11837,PF00251 | 3.2.1.26, 2.4.1.99 | GO:0004575, GO:0004564 | Glycosyl hydrolases family 32 protein | glycosyl hydrolases, putative, expressed |
| Bradi2g03280.1 | up | core (33) | D0 | W0 | | | PF01439 | - | GO:0046872 | metallothionein 2A | metallothionein, putative, expressed |
| Bradi3g49260.1 | up | core (33) | D0 | W3 | | | PF00221 | 4.3.1.24 | - | PHE ammonia lyase 1 | phenylalanine ammonia-lyase, putative, expressed |
| Bradi2g06220.1 | up | Shell (25) | D0 | W0 | | | PF01439 | - | GO:0046872 | metallothionein 3 | metallothionein-like protein 3B, putative, expressed |
| Bradi1g75610.1 | up | soft-core (32) | D0 | W0 | | | PF01373 | 3.2.1.2 | GO:0016161, GO:0000272 | beta-amylase 1 | beta-amylase, putative, expressed |
| Bradi1g37410.1 | up | soft-core (32) | D0 | W0 | | | PF00257 | - | GO:0009415, GO:0006950 | dehydrin xero 1 | dehydrin, putative, expressed |
| Bradi1g20950.1 | up | core (33) | D6 | W3 | hub (D6) | hub (W3) | PF03018 | - | - | Disease resistance-responsive (dirigent-like protein) family protein | dirigent, putative, expressed |
| Bradi3g39800.1 | up | core (33) | D0 | W0 | | | PF03600 | - | GO:0055085, GO:0016021 | tonoplast dicarboxylate transporter | citrate transporter, putative, expressed |
| Bradi3g50220.1 | up | soft-core (32) | D0 | W0 | | | PF02183,PF00046 | - | GO:0043565, GO:0006355, GO:0003700, GO:0003677 | homeobox 7 | homeobox associated leucine zipper, putative, expressed |

| | | | | | | | | | | |
|---|---|---|---|---|---|---|---|---|---|---|
| Bradi5g17170.2 | up | soft-core (32) | D0 | W8 | | PF02183,PF00046 | - | GO:0043565, GO:0006355, GO:0003700, GO:0003677 | homeobox 7 | homeobox associated leucine zipper, putative, expressed |
| Bradi1g65780.1 | up | core (33) | D0 | W8 | | PF01679 | - | GO:0016021 | Low temperature and salt responsive protein family | OsRCI2-5 - Putative low temperature and salt responsive protein, expressed |
| Bradi3g51200.1 | up | core (33) | D0 | W6 | | PF00257 | - | GO:0009415, GO:0006950 | cold-regulated 47 | dehydrin, putative, expressed |
| Bradi3g14960.1 | up | core (33) | D0 | W3 | | PF00903 | 4.4.1.5 | - | glyoxalase I homolog | glyoxalase family protein, putative, expressed |
| Bradi5g10027.1 | up | core (33) | D0 | W0 | | PF02496 | - | GO:0006950 | - | abscisic stress-ripening, putative, expressed |
| Bradi1g55560.1 | up | Shell (29) | D0 | W3 | | PF00504 | - | - | Chlorophyll A-B binding family protein | early light-induced protein, chloroplast precursor, putative, expressed |
| Bradi2g54920.1 | up | core (33) | D0 | W8 | | PF00171,PF00696 | 1.2.1.41, 2.7.2.11 | GO:0055114, GO:0016491, GO:0008152 | delta 1-pyrroline-5-carboxylate synthase 2 | amino acid kinase, putative, expressed |
| Bradi3g05560.1 | up | core (33) | D0 | W0 | | PF04570 | - | - | Protein of unknown function (DUF581) | DUF581 domain containing protein, expressed |

| | | | | | | | | | | | |
|---|---|---|---|---|---|---|---|---|---|---|---|
| Bradi4g23800.2 | up | core (33) | D0 | W10 | | | PF04116 | 1.14.13.72 | GO:0055114, GO:0016491, GO:0006633, GO:0005506 | sterol C4-methyl oxidase 1-2 | fatty acid hydroxylase, putative, expressed |
| Bradi2g35620.1 | up | core (33) | D1 | W2 | hub (D1) | | PF00467,PF01479,PF08071,PF16121,PF00900 | - | GO:0003723, GO:0006412, GO:0005840, GO:0005622, GO:0003735 | Ribosomal protein S4 (RPS4A) family protein | 40S ribosomal protein S4, putative, expressed |
| Bradi2g05760.1 | up | core (33) | D0 | W0 | | | PF00249 | - | - | myb-like transcription factor family protein | Myb transcription factor, putative, expressed |
| Bradi2g11940.1 | down | core (33) | D9 | W0 | hub (D9) | | PF03242 | - | GO:0006950 | senescence-associated gene 21 | late embryogenesis abundant protein, putative, expressed |
| Bradi3g44820.2 | down | core (33) | D2 | W7 | | | PF00403 | - | GO:0046872, GO:0030001 | homolog of anti-oxidant 1 | heavy metal-associated domain containing protein, expressed |
| Bradi1g29000.1 | down | core (33) | D2 | W10 | hub (D2) | hub (W10) | PF02427 | - | GO:0015979, GO:0009538, GO:0009522 | photosystem I subunit E-2 | photosystem I reaction center subunit IV A, chloroplast precursor, putative, expressed |
| Bradi4g40740.1 | down | core (33) | D2 | W0 | | | - | 2.7.7.69 | GO:0080048 | galactose-1-phosphate guanylyltransferase (GDP)s;GDP-D- | VTC2, putative, expressed |

| | | | | | | | | | | | |
|---|---|---|---|---|---|---|---|---|---|---|---|
| | | | | | | | | | | glucose phosphorylases;quercetin 4\'-O-glucosyltransferases | |
| Bradi3g39750.2 | down | core (33) | D0 | W0 | | | PF00266 | 2.6.1.51, 2.6.1.44, 2.6.1.45 | - | alanine:glyoxylate aminotransferase | aminotransferase, putative, expressed |
| Bradi4g21170.1 | down | core (33) | D2 | W10 | | | PF00504 | - | GO:0016020, GO:0009765 | light harvesting complex of photosystem II 5 | chlorophyll A-B binding protein, putative, expressed |
| Bradi3g57990.1 | down | core (33) | D2 | W0 | | | PF03460,PF01077 | 1.7.7.1 | GO:0055114, GO:0016491, GO:0051536, GO:0020037 | nitrite reductase 1 | ferredoxin--nitrite reductase, putative, expressed |
| Bradi1g26530.1 | down | core (33) | D0 | W0 | | | PF00085,PF01507 | 1.8.4.10 | GO:0045454, GO:0008152, GO:0003824 | APS reductase 3 | OsAPRL1 adenosine 5\'-phosphosulfate reductase-like OsAPRL1, expressed |
| Bradi1g58160.1 | down | core (33) | D2 | W1 | hub (D2) | | PF01789 | - | GO:0019898, GO:0015979, GO:0009654, GO:0009523, GO:0005509 | photosystem II subunit P-1 | PsbP, putative, expressed |
| Bradi1g18050.1 | down | Shell (30) | D2 | W10 | hub (D2) | hub (W10) | PF06596 | - | GO:0016020, GO:0015979, GO:0009523 | photosystem II subunit X | ultraviolet-B-repressible protein, putative, expressed |
| Bradi1g76470.1 | down | core (33) | D2 | W0 | hub (D2) | | PF02672,PF0004 | 1.2.1.12, 1.2.1.13 | GO:0055114, GO:0016620 | glyceraldehyde-3-phosphate | glyceraldehyde-3-phosphate dehydrogenase, |

| | | | | | | | | | | | |
|---|---|---|---|---|---|---|---|---|---|---|---|
| | | | | | | | 4,PF02800 | | | dehydrogenase B subunit | putative, expressed |
| Bradi5g09530.1 | down | core (33) | D2 | W10 | | | - | 1.97.1.12 | - | photosystem I subunit O | membrane protein, putative, expressed |
| Bradi2g44856.2 | down | core (33) | D2 | W10 | hub (D2) | hub (W10) | PF00484 | 4.2.1.1 | GO:0008270, GO:0004089 | carbonic anhydrase 2 | carbonic anhydrase, chloroplast precursor, putative, expressed |
| Bradi1g24980.1 | down | core (33) | D2 | W0 | | | PF00355,PF08802 | 1.10.9.1 | GO:0055114, GO:0051537, GO:0016491, GO:0042651, GO:0009496, GO:0016679 | photosynthetic electron transfer C | cytochrome b6-f complex iron-sulfur subunit, chloroplast precursor, putative, expressed |
| Bradi1g59220.1 | down | core (33) | D1 | W1 | | | PF00155 | 2.3.2.2,2.6.1.4,2.6.1.2,2.6.1.44 | GO:0030170, GO:0009058 | alanine-2-oxoglutarate aminotransferase 2 | aminotransferase, classes I and II, domain containing protein, expressed |
| Bradi1g09300.3 | down | core (33) | D2 | W1 | | | PF00464 | 2.1.2.1 | GO:0016740 | serine transhydroxymethyltransferase 1 | serine hydroxymethyltransferase, mitochondrial precursor, putative, expressed |
| Bradi1g24870.1 | down | soft-core (32) | D2 | W10 | hub (D2) | hub (W10) | PF00504 | - | GO:0016020, GO:0009765 | light harvesting complex photosystem II | chlorophyll A-B binding protein, putative, expressed |
| Bradi3g13170.1 | down | soft-core (32) | D2 | W10 | hub (D2) | hub (W10) | PF00111 | - | GO:0051536, GO:0009055 | 2Fe-2S ferredoxin-like | 2Fe-2S iron-sulfur cluster |

| | | | | | | | | | | | |
|---|---|---|---|---|---|---|---|---|---|---|---|
| | | | | | | | | | | superfamily protein | binding domain containing protein, expressed |
| Bradi2g51480.1 | down | soft-core (31) | D2 | W10 | hub (D2) | | PF07123 | - | GO:0015979, GO:0009523, GO:0009507 | photosystem II reaction center W | photosystem II reaction center W protein, chloroplast precursor, putative, expressed |
| Bradi1g78086.2 | down | Shell (20) | D1 | W4 | | | PF01737 | - | GO:0042549, GO:0015979, GO:0009539, GO:0009523 | YCF9 | photosystem II reaction center protein Z, putative, expressed |
| Bradi1g05801.1 | down | - | D0 | W0 | | | PF00177 | - | GO:0006412 | ribosomal protein S7 | chloroplast 30S ribosomal protein S7, putative, expressed |
| Bradi4g08500.2 | down | core (33) | D2 | W10 | hub (D2) | hub (W10) | PF12338 ,PF0010 1 | 4.1.1.39 | - | ribulose bisphosphate carboxylase small chain 1A | ribulose bisphosphate carboxylase small chain, chloroplast precursor, putative, expressed |
| Bradi1g56580.1 | down | core (33) | D2 | W10 | hub (D2) | hub (W10) | PF01716 | - | GO:0042549, GO:0019898, GO:0015979, GO:0009654, GO:0009523, GO:0005509 | photosystem II subunit O-2 | oxygen-evolving enhancer protein 1, chloroplast precursor, putative, expressed |

| | | | | | | | | | | | |
|---|---|---|---|---|---|---|---|---|---|---|---|
| Bradi4g08800.1 | down | soft-core (32) | D2 | W10 | hub (D2) | hub (W10) | PF12338,PF00101 | 4.1.1.39 | - | ribulose bisphosphate carboxylase small chain 1A | ribulose bisphosphate carboxylase small chain, chloroplast precursor, putative, expressed |
| Bradi4g09125.1 | down | core (33) | D2 | W10 | hub (D2) | | PF00004 | - | GO:0005524 | rubisco activase | AAA-type ATPase family protein, putative, expressed |

**Table S8.** Summary annotation of the differentially expressed (DE) isoforms of the Drought module 5 (D5). Isoform (*B. distachyon* 314 v3.1 reference transcriptome); regulation (regulation under drought condition comparing to water condition); occupancy (core (33 accessions), soft-core (31-32 accessions) and shell (≤30 accessions)); DE hub node (DE gene detected such as hub gene in the module); Pfam, KEGG/ec, GO annotations; arabi-defline and rice-defline (annotation in Arabidopsis and rice). The dashes indicate that no results were retrieved.

| Isoform | regulation | occupancy | DE hub node | Pfam | KEGG/ec | GO | arabi-defline | rice-defline |
|---|---|---|---|---|---|---|---|---|
| Bradi1g01507.1 | | core (33) | | PF00684, PF00226, PF01556 | - | GO:0051082, GO:0031072 | gametophytic factor 2 | chaperone protein dnaJ, putative, expressed |
| Bradi1g03590.1 | | core (33) | | PF08609 | - | - | Fes1A | armadillo/beta-catenin-like repeat containing protein, expressed |
| Bradi1g03590.2 | | core (33) | | PF08609 | - | - | Fes1A | armadillo/beta-catenin-like repeat containing protein, expressed |
| Bradi1g03610.1 | up | core (33) | | PF13460 | - | - | NAD(P)-binding Rossmann-fold superfamily protein | expressed protein |
| Bradi1g06340.1 | | core (33) | | PF00684, PF00226, PF01556 | - | GO:0051082, GO:0031072 | DNAJ homologue 2 | chaperone protein dnaJ, putative, expressed |
| Bradi1g06980.4 | | shell (27) | | - | - | - | Chaperone DnaJ-domain superfamily protein | heat shock protein DnaJ, putative, expressed |
| Bradi1g06980.6 | | shell (27) | | - | - | - | Chaperone DnaJ-domain superfamily protein | heat shock protein DnaJ, putative, expressed |
| Bradi1g06980.7 | | shell (27) | | - | - | - | Chaperone DnaJ-domain superfamily protein | heat shock protein DnaJ, putative, expressed |
| Bradi1g08410.4 | up | soft-core (31) | | - | - | - | Heavy metal transport/detoxification superfamily protein | anther-specific proline-rich protein APG precursor, putative, expressed |
| Bradi1g08870.1 | up | core (33) | x | PF01027 | - | - | Bax inhibitor-1 family protein | transmembrane BAX inhibitor motif-containing protein, putative, expressed |

| | | | | | | | | |
|---|---|---|---|---|---|---|---|---|
| Bradi1g09950.2 | up | core (33) | x | - | - | - | ethylene-dependent gravitropism-deficient and yellow-green-like 3 | peptidase, M50 family, putative, expressed |
| Bradi1g11630.1 | | core (33) | | PF08799, PF02840 | - | GO:0008380, GO:0005681 | splicing factor Prp18 family protein | pre-mRNA-splicing factor, putative, expressed |
| Bradi1g16190.1 | up | core (33) | x | PF07728, PF10431, PF00004, PF02861 | - | GO:0016887, GO:0005524, GO:0019538 | casein lytic proteinase B3 | chaperone protein clpB 1, putative, expressed |
| Bradi1g19860.1 | up | core (33) | | PF00166 | - | GO:0006457, GO:0005737 | chaperonin 10 | chaperonin, putative, expressed |
| Bradi1g20360.1 | | core (33) | | PF00076 | - | GO:0003676 | SC35-like splicing factor 33 | RNA recognition motif containing protein, putative, expressed |
| Bradi1g20620.1 | up | core (33) | | PF01926, PF02824, PF16897 | - | GO:0005525 | GTP-binding protein-related | GTP-binding protein, putative, expressed |
| Bradi1g23653.1 | | shell (25) | | - | - | - | - | - |
| Bradi1g29040.1 | up | core (33) | | PF13347 | - | - | Major facilitator superfamily protein | integral membrane transporter family protein, putative, expressed |
| Bradi1g30247.1 | | core (33) | | PF00240 | - | GO:0005515 | Ubiquitin-like superfamily protein | ubiquitin family protein, putative, expressed |
| Bradi1g30247.2 | | core (33) | | PF00240 | - | GO:0005515 | Ubiquitin-like superfamily protein | ubiquitin family protein, putative, expressed |
| Bradi1g30247.3 | | core (33) | | PF00240 | - | GO:0005515 | Ubiquitin-like superfamily protein | ubiquitin family protein, putative, expressed |
| Bradi1g32970.1 | up | core (33) | | PF04818, PF01805 | - | GO:0006396, GO:0003723 | SWAP (Suppressor-of-White-APricot)/surp RNA-binding domain-containing protein | surp module family protein, putative, expressed |
| Bradi1g32990.1 | up | core (33) | x | PF02679 | 4.4.1.19 | GO:0019295 | Aldolase-type TIM barrel family protein | phosphosulfolactate synthase-related protein, putative, expressed |
| Bradi1g35736.1 | | soft-core (32) | | PF01370 | - | GO:0050662, GO:0003824 | NAD(P)-binding Rossmann-fold superfamily protein | reductase, putative, expressed |

| | | | | | | | | |
|---|---|---|---|---|---|---|---|---|
| Bradi1g36290.1 | up | core (33) | | PF03660 | - | - | PHF5-like protein | PHF5-like protein domain containing protein, expressed |
| Bradi1g37080.1 | | soft-core (32) | | PF01381, PF08523 | - | GO:0043565 | multiprotein bridging factor 1C | endothelial differentiation-related factor 1, putative, expressed |
| Bradi1g37175.1 | | core (33) | | PF00106 | 1.1.1.300 | GO:0016491, GO:0008152 | NAD(P)-binding Rossmann-fold superfamily protein | dehydrogenase/reductase SDR family member 12, putative, expressed |
| Bradi1g37175.2 | | core (33) | | PF00106 | 1.1.1.300 | GO:0016491, GO:0008152 | NAD(P)-binding Rossmann-fold superfamily protein | dehydrogenase/reductase SDR family member 12, putative, expressed |
| Bradi1g37175.3 | | core (33) | | PF00106 | 1.1.1.300 | GO:0016491, GO:0008152 | NAD(P)-binding Rossmann-fold superfamily protein | dehydrogenase/reductase SDR family member 12, putative, expressed |
| Bradi1g37750.7 | up | core (33) | | PF03062 | 2.3.1.20 | GO:0008374 | membrane bound O-acyl transferase (MBOAT) family protein | O-acyltransferase, putative, expressed |
| Bradi1g38760.1 | up | core (33) | | PF01965 | - | - | Class I glutamine amidotransferase-like superfamily protein | DJ-1 family protein, putative, expressed |
| Bradi1g42730.1 | up | core (33) | | PF13857, PF00651 | - | GO:0005515 | ankyrin repeat family protein | ABB1 - Ankyrin repeat region with 2 Bric-a-Brac, Tramtrack, Broad Complex BTB domains, expressed |
| Bradi1g44230.1 | | core (33) | x | PF00011 | - | - | HSP20-like chaperones superfamily protein | hsp20/alpha crystallin family protein, putative, expressed |
| Bradi1g45080.1 | up | core (33) | | PF04969 | - | - | HSP20-like chaperones superfamily protein | CS domain containing protein, putative, expressed |
| Bradi1g45160.1 | up | core (33) | x | PF07728, PF01434 | - | GO:0016887, GO:0005524, GO:0006508, GO:0004222 | FTSH protease 6 | OsFtsH6 FtsH protease, homologue of AtFtsH6, expressed |
| Bradi1g46530.1 | | core (33) | | PF02441 | 4.1.1.36 | GO:0003824 | HAL3-like protein A | phosphopantothenoylcysteine decarboxylase, putative, expressed |

| | | | | | | | | |
|---|---|---|---|---|---|---|---|---|
| Bradi1g50370.1 | up | soft-core (32) | | - | - | - | F-box family protein with a domain of unknown function (DUF295) | OsFBX182 - F-box domain containing protein, expressed |
| Bradi1g51610.1 | up | core (33) | x | PF10248 | - | - | glycine-rich protein | expressed protein |
| Bradi1g51610.2 | up | core (33) | x | PF10248 | - | - | glycine-rich protein | expressed protein |
| Bradi1g53295.1 | up | shell (20) | | - | - | - | - | - |
| Bradi1g53850.1 | up | soft-core (31) | | PF00011 | - | - | HSP20-like chaperones superfamily protein | hsp20/alpha crystallin family protein, putative, expressed |
| Bradi1g55090.1 | | core (33) | | PF00628, PF01426 | - | GO:0005515, GO:0003682 | PHD finger family protein / bromo-adjacent homology (BAH) domain-containing protein | ES43 protein, putative, expressed |
| Bradi1g55620.1 | up | shell (26) | | PF00504 | - | - | Chlorophyll A-B binding family protein | early light-induced protein, chloroplast precursor, putative, expressed |
| Bradi1g61040.1 | up | core (33) | x | PF04969 | - | - | HSP20-like chaperones superfamily protein | CS domain containing protein, putative, expressed |
| Bradi1g61170.1 | | core (33) | | PF06423 | - | GO:0016746, GO:0016021, GO:0006506, GO:0005789 | transferases, transferring acyl groups | GPI-anchored wall transfer protein 1, putative, expressed |
| Bradi1g61320.2 | up | core (33) | | PF07983, PF00332 | 3.2.1.39 | GO:0005975, GO:0004553 | O-Glycosyl hydrolases family 17 protein | glycosyl hydrolases family 17 protein, expressed |
| Bradi1g62420.1 | | core (33) | | PF06203 | - | GO:0005515 | B-box type zinc finger protein with CCT domain | CCT/B-box zinc finger protein, putative, expressed |
| Bradi1g63816.1 | up | core (33) | | PF02987 | - | - | late embryogenesis abundant protein, putative / LEA protein, putative | late embryogenesis abundant protein 1, putative, expressed |
| Bradi1g64850.1 | | core (33) | | PF00226 | - | - | Chaperone DnaJ-domain superfamily protein | heat shock protein DnaJ, putative, expressed |
| Bradi1g65480.1 | up | core (33) | | PF00226 | - | - | DNAJ heat shock N-terminal domain-containing protein | heat shock protein DnaJ, putative, expressed |
| Bradi1g66800.1 | up | core (33) | x | - | - | - | Fes1A | expressed protein |
| Bradi1g66800.2 | up | core (33) | x | - | - | - | Fes1A | expressed protein |

| | | | | | | | | |
|---|---|---|---|---|---|---|---|---|
| Bradi1g66800.3 | up | core (33) | x | - | - | - | Fes1A | expressed protein |
| Bradi1g67040.1 | up | core (33) | x | PF00011 | - | - | HSP20-like chaperones superfamily protein | hsp20/alpha crystallin family protein, putative, expressed |
| Bradi1g68720.1 | | soft-core (32) | | PF05603 | - | - | - | expressed protein |
| Bradi1g72086.1 | up | core (33) | | PF00400 | - | GO:0005515 | transducin family protein / WD-40 repeat family protein | WD domain, G-beta repeat domain containing protein, expressed |
| Bradi1g72950.1 | | core (33) | | PF07685 | 4.3.3.6 | GO:0009236, GO:0003824 | pyridoxine biosynthesis 2 | pyridoxal biosynthesis protein PDX2, putative, expressed |
| Bradi1g73890.1 | up | shell (29) | | PF04788 | - | - | Protein of unknown function (DUF620) | expressed protein |
| Bradi1g74300.1 | up | core (33) | | PF01680 | 4.3.3.6 | GO:0042823, GO:0042819 | Aldolase-type TIM barrel family protein | SOR/SNZ family protein, putative, expressed |
| Bradi1g77637.1 | up | core (33) | x | PF00012 | - | - | mitochondrial HSO70 2 | DnaK family protein, putative, expressed |
| Bradi2g00831.1 | up | shell (26) | | PF03893, PF01764 | 3.1.1.3 | GO:0016042, GO:0006629 | alpha/beta-Hydrolases superfamily protein | calmodulin-binding heat-shock protein, putative, expressed |
| Bradi2g02400.1 | up | shell (26) | | PF00011 | - | - | HSP20-like chaperones superfamily protein | hsp20/alpha crystallin family protein, putative, expressed |
| Bradi2g03840.1 | up | shell (23) | | PF02775, PF00205, PF02776 | 4.1.1.1 | GO:0030976, GO:0003824, GO:0000287 | Thiamine pyrophosphate dependent pyruvate decarboxylase family protein | thiamine pyrophosphate enzyme, C-terminal TPP binding domain containing protein, expressed |
| Bradi2g07040.1 | | soft-core (32) | | PF01965 | - | - | Class I glutamine amidotransferase-like superfamily protein | DJ-1 family protein, putative, expressed |
| Bradi2g07040.2 | | soft-core (32) | | PF01965 | - | - | Class I glutamine amidotransferase-like superfamily protein | DJ-1 family protein, putative, expressed |
| Bradi2g08330.1 | up | core (33) | | PF00226, PF01556 | - | - | DNAJ heat shock family protein | dnaJ domain containing protein, expressed |
| Bradi2g10650.1 | | core (33) | | PF12710, PF16209, PF16212 | 3.6.3.1 | GO:0016021, GO:0015914, GO:0005524, | aminophospholipid ATPase 1 | phospholipid-transporting ATPase, putative, expressed |

| | | | | | | | | |
|---|---|---|---|---|---|---|---|---|
| | | | | | | GO:0004012, GO:0000287 | | |
| Bradi2g12830.1 | up | core (33) | | PF04535 | - | - | Uncharacterised protein family (UPF0497) | membrane associated DUF588 domain containing protein, putative, expressed |
| Bradi2g13060.1 | | core (33) | | PF00043, PF02798 | 2.5.1.18 | GO:0005515 | Glutathione S-transferase family protein | glutathione S-transferase, putative, expressed |
| Bradi2g13110.1 | | core (33) | | PF02798, PF00043 | 2.5.1.18 | GO:0005515 | Glutathione S-transferase family protein | glutathione S-transferase, putative, expressed |
| Bradi2g13870.1 | | core (33) | x | - | - | - | ARM repeat superfamily protein | armadillo/beta-catenin-like repeat family protein, expressed |
| Bradi2g13870.10 | | core (33) | x | - | - | - | ARM repeat superfamily protein | armadillo/beta-catenin-like repeat family protein, expressed |
| Bradi2g13870.13 | | core (33) | x | - | - | - | ARM repeat superfamily protein | armadillo/beta-catenin-like repeat family protein, expressed |
| Bradi2g13870.5 | | core (33) | x | - | - | - | ARM repeat superfamily protein | armadillo/beta-catenin-like repeat family protein, expressed |
| Bradi2g13870.6 | | core (33) | x | - | - | - | ARM repeat superfamily protein | armadillo/beta-catenin-like repeat family protein, expressed |
| Bradi2g13870.8 | | core (33) | x | - | - | - | ARM repeat superfamily protein | armadillo/beta-catenin-like repeat family protein, expressed |
| Bradi2g15950.1 | | shell (30) | | PF11341 | - | - | - | expressed protein |
| Bradi2g16660.1 | | core (33) | | PF00226, PF01556 | - | - | DNAJ heat shock family protein | dnaJ domain containing protein, expressed |
| Bradi2g17200.3 | down | core (33) | | PF00111 | - | GO:0051536, GO:0009055 | 2Fe-2S ferredoxin-like superfamily protein | 2Fe-2S iron-sulfur cluster binding domain containing protein, expressed |
| Bradi2g17620.1 | up | soft-core (31) | | PF00291 | 4.2.3.1 | - | Pyridoxal-5\'-phosphate-dependent enzyme family protein | threonine synthase, chloroplast precursor, putative, expressed |
| Bradi2g19540.1 | up | core (33) | | PF07728, PF02861, PF10431, PF00004 | - | GO:0016887, GO:0005524, GO:0019538 | heat shock protein 101 | heat shock protein 101, putative, expressed |

| | | | | | | | | |
|---|---|---|---|---|---|---|---|---|
| Bradi2g22460.1 | up | core (33) | | PF00248 | 1.1.1.21 | - | NAD(P)-linked oxidoreductase superfamily protein | oxidoreductase, aldo/keto reductase family protein, putative, expressed |
| Bradi2g23040.2 | | core (33) | | PF16876, PF04571, PF08235 | 3.1.3.4 | - | Lipin family protein | lipin, N-terminal conserved region family protein, expressed |
| Bradi2g23040.3 | | core (33) | | PF16876, PF04571, PF08235 | 3.1.3.4 | - | Lipin family protein | lipin, N-terminal conserved region family protein, expressed |
| Bradi2g23040.4 | | core (33) | | PF16876, PF04571, PF08235 | 3.1.3.4 | - | Lipin family protein | lipin, N-terminal conserved region family protein, expressed |
| Bradi2g23040.5 | | core (33) | | PF16876, PF04571, PF08235 | 3.1.3.4 | - | Lipin family protein | lipin, N-terminal conserved region family protein, expressed |
| Bradi2g23250.1 | up | soft-core (31) | x | PF00012 | - | - | heat shock protein 70 | DnaK family protein, putative, expressed |
| Bradi2g24230.1 | up | core (33) | | PF17032 | - | - | - | expressed protein |
| Bradi2g25030.1 | | soft-core (32) | | PF00034 | - | GO:0020037, GO:0009055 | cytochrome c-2 | cytochrome c, putative, expressed |
| Bradi2g25613.1 | | core (33) | | PF04419 | - | - | Uncharacterised protein family SERF | 4F5 protein family protein, expressed |
| Bradi2g27900.1 | up | core (33) | | PF04398 | - | - | Protein of unknown function, DUF538 | expressed protein |
| Bradi2g33360.3 | | core (33) | | PF04511 | - | - | DERLIN-1 | Der1-like family domain containing protein, expressed |
| Bradi2g33682.2 | | core (33) | | PF00012 | - | - | heat shock protein 91 | DnaK family protein, putative, expressed |
| Bradi2g37480.1 | up | core (33) | | PF13410, PF13417 | 1.8.5.1 | GO:0005515 | dehydroascorbate reductase 2 | glutathione S-transferase, N-terminal domain containing protein, expressed |
| Bradi2g41230.3 | | soft-core (32) | | PF06201 | - | - | Protein of unknown function (DUF1000) | thioredoxin, putative, expressed |

| | | | | | | | | |
|---|---|---|---|---|---|---|---|---|
| Bradi2g45030.1 | up | core (33) | | - | - | - | - | tat pathway signal sequence family protein, expressed |
| Bradi2g45840.1 | up | core (33) | | PF01544 | - | GO:0055085, GO:0046873, GO:0030001, GO:0016020 | Magnesium transporter CorA-like family protein | expressed protein |
| Bradi2g47530.3 | down | core (33) | | - | - | - | - | expressed protein |
| Bradi2g47530.4 | down | core (33) | | - | - | - | - | expressed protein |
| Bradi2g47530.5 | down | core (33) | | - | - | - | - | expressed protein |
| Bradi2g48230.1 | | core (33) | | - | 1.10.99.3 | - | violaxanthin de-epoxidase-related | violaxanthin de-epoxidase, putative, expressed |
| Bradi2g49280.1 | | core (33) | | PF00365 | 2.7.1.11 | GO:0006096, GO:0003872 | phosphofructokinase 3 | 6-phosphofructokinase, putative, expressed |
| Bradi2g49660.1 | | core (33) | x | PF07728, PF02861, PF10431, PF00004 | - | GO:0016887, GO:0005524, GO:0019538 | heat shock protein 101 | heat shock protein 101, putative, expressed |
| Bradi2g50300.1 | up | core (33) | x | PF09032, PF04969 | - | - | SGS domain-containing protein | SGS domain containing protein, expressed |
| Bradi2g51240.1 | up | core (33) | x | - | - | - | - | mal d 1-associated protein, putative, expressed |
| Bradi2g51980.1 | down | core (33) | | PF03358 | 1.6.5.2 | GO:0016491 | flavodoxin-like quinone reductase 1 | NADPH-dependent FMN reductase domain containing protein, expressed |
| Bradi2g54570.1 | | core (33) | x | PF00012 | - | - | heat shock protein 70 | DnaK family protein, putative, expressed |
| Bradi2g56190.1 | up | core (33) | | PF04842 | - | - | Plant protein of unknown function (DUF639) | expressed protein |
| Bradi2g56190.2 | up | core (33) | | PF04842 | - | - | Plant protein of unknown function (DUF639) | expressed protein |
| Bradi2g56490.1 | up | core (33) | x | - | - | - | - | expressed protein |
| Bradi2g58835.1 | up | soft-core (32) | | - | - | - | - | expressed protein |

| | | | | | | | | |
|---|---|---|---|---|---|---|---|---|
| Bradi2g61767.1 | | core (33) | | - | - | - | Nuclear transport factor 2 (NTF2) family protein | expressed protein |
| Bradi3g02200.1 | | soft-core (31) | | PF13301 | - | - | - | expressed protein |
| Bradi3g02520.2 | | core (33) | | PF01027 | - | - | BAX inhibitor 1 | transmembrane BAX inhibitor motif-containing protein, putative, expressed |
| Bradi3g03532.1 | | soft-core (32) | | PF09229 | - | GO:0051087, GO:0001671 | chaperone binding;ATPase activators | activator of 90 kDa heat shock protein ATPase homolog, putative, expressed |
| Bradi3g05530.1 | up | soft-core (32) | x | PF13837 | - | - | sequence-specific DNA binding transcription factors | transcription factor, putative, expressed |
| Bradi3g06107.2 | up | core (33) | x | PF07728, PF10431, PF00004, PF02861 | - | GO:0016887, GO:0005524, GO:0019538 | casein lytic proteinase B4 | chaperone protein clpB 1, putative, expressed |
| Bradi3g08730.1 | up | shell (27) | | PF05553 | - | - | - | expressed protein |
| Bradi3g08770.1 | | core (33) | | PF12799 | - | - | Leucine-rich repeat (LRR) family protein | protein phosphatase 1 regulatory subunit SDS22, putative, expressed |
| Bradi3g11130.1 | up | core (33) | | PF01168, PF00278 | 4.1.1.20 | GO:0003824 | Pyridoxal-dependent decarboxylase family protein | pyridoxal-dependent decarboxylase protein, putative, expressed |
| Bradi3g19893.1 | up | shell (29) | | - | - | - | - | - |
| Bradi3g22980.2 | | core (33) | | PF00107, PF08240 | 1.1.1.183 | GO:0055114, GO:0016491, GO:0008270 | cinnamyl-alcohol dehydrogenase | dehydrogenase, putative, expressed |
| Bradi3g22980.3 | | core (33) | | PF00107, PF08240 | 1.1.1.183 | GO:0055114, GO:0016491, GO:0008270 | cinnamyl-alcohol dehydrogenase | dehydrogenase, putative, expressed |
| Bradi3g22980.5 | | core (33) | | PF00107, PF08240 | 1.1.1.183 | GO:0055114, GO:0016491, GO:0008270 | cinnamyl-alcohol dehydrogenase | dehydrogenase, putative, expressed |

| | | | | | | | |
|---|---|---|---|---|---|---|---|
| Bradi3g22980.6 | | core (33) | PF00107, PF08240 | 1.1.1.183 | GO:0055114, GO:0016491, GO:0008270 | cinnamyl-alcohol dehydrogenase | dehydrogenase, putative, expressed |
| Bradi3g22980.7 | | core (33) | PF00107, PF08240 | 1.1.1.183 | GO:0055114, GO:0016491, GO:0008270 | cinnamyl-alcohol dehydrogenase | dehydrogenase, putative, expressed |
| Bradi3g22980.8 | | core (33) | PF00107, PF08240 | 1.1.1.183 | GO:0055114, GO:0016491, GO:0008270 | cinnamyl-alcohol dehydrogenase | dehydrogenase, putative, expressed |
| Bradi3g22980.9 | | core (33) | PF00107, PF08240 | 1.1.1.183 | GO:0055114, GO:0016491, GO:0008270 | cinnamyl-alcohol dehydrogenase | dehydrogenase, putative, expressed |
| Bradi3g26967.1 | | soft-core (32) | PF12695 | 3.1.1.14 | - | chlorophyllase 2 | chlorophyllase-2, chloroplast precursor, putative, expressed |
| Bradi3g28070.1 | | core (33) | PF00118 | - | GO:0005524 | heat shock protein 60 | T-complex protein, putative, expressed |
| Bradi3g29797.1 | up | soft-core (32) | PF16499 | 3.2.1.22 | - | alpha-galactosidase 1 | alpha-galactosidase precursor, putative, expressed |
| Bradi3g30880.1 | up | core (33) | PF00566 | - | - | Ypt/Rab-GAP domain of gyp1p superfamily protein | TBC domain containing protein, expressed |
| Bradi3g33300.1 | | core (33) | PF08458, PF05703 | - | - | Plant protein of unknown function (DUF828) with plant pleckstrin homology-like region | expressed protein |
| Bradi3g33544.1 | down | core (33) | PF01625 | 1.8.4.11 | GO:0055114, GO:0008113, GO:0030091, GO:0016671, GO:0006979 | peptide met sulfoxide reductase 4 | peptide methionine sulfoxide reductase, putative, expressed |
| Bradi3g34980.1 | | shell (25) | PF08240, PF13602 | 1.3.1.74, 1.3.1.105 | GO:0055114, GO:0016491, GO:0008270 | Oxidoreductase, zinc-binding dehydrogenase family protein | dehydrogenase, putative, expressed |
| Bradi3g37260.1 | up | core (33) | PF07093 | - | - | - | SGT1 protein, putative, expressed |

| | | | | | | | | |
|---|---|---|---|---|---|---|---|---|
| Bradi3g37790.1 | | core (33) | | PF08327, PF09229 | - | GO:0006950, GO:0051087, GO:0001671 | Aha1 domain-containing protein | activator of 90 kDa heat shock protein ATPase homolog, putative, expressed |
| Bradi3g37820.3 | up | core (33) | | PF00067 | 1.14.13.192 | GO:0055114, GO:0020037, GO:0016705, GO:0005506 | cytochrome P450, family 76, subfamily C, polypeptide 4 | cytochrome P450, putative, expressed |
| Bradi3g38770.1 | down | soft-core (31) | | PF03931, PF01466 | - | GO:0006511 | SKP1-like 4 | Skp1 family, dimerisation domain containing protein, expressed |
| Bradi3g44947.1 | | shell (1) | | - | - | - | - | - |
| Bradi3g44950.1 | | core (33) | | - | - | - | F1F0-ATPase inhibitor protein, putative | expressed protein |
| Bradi3g45030.1 | up | core (33) | | PF04968 | - | - | protein binding;zinc ion binding | rar1, putative, expressed |
| Bradi3g45030.2 | up | core (33) | | PF04968 | - | - | protein binding;zinc ion binding | rar1, putative, expressed |
| Bradi3g45030.3 | up | core (33) | | PF04968 | - | - | protein binding;zinc ion binding | rar1, putative, expressed |
| Bradi3g45030.4 | up | core (33) | | PF04968 | - | - | protein binding;zinc ion binding | rar1, putative, expressed |
| Bradi3g48240.1 | | core (33) | | PF03194 | - | GO:0006376, GO:0005685, GO:0003729 | LUC7 related protein | RNA-binding protein Luc7-like, putative, expressed |
| Bradi3g48590.1 | | core (33) | | PF00061 | 1.14.13.90 | - | temperature-induced lipocalin | OsTIL-1 Temperature-induced lipocalin-1, expressed |
| Bradi3g50110.1 | | core (33) | | PF13181, PF00515, PF13414 | - | GO:0005515 | stress-inducible protein, putative | heat shock protein STI, putative, expressed |
| Bradi3g52367.2 | | core (33) | | PF00004, PF12037 | - | GO:0005524 | AAA-type ATPase family protein | AAA-type ATPase family protein, putative, expressed |
| Bradi3g54267.1 | up | soft-core (31) | x | PF05536 | - | - | neurochondrin family protein | neurochondrin family protein, putative, expressed |
| Bradi3g54330.1 | up | soft-core (31) | | - | - | - | serine-rich protein-related | expressed protein |

| | | | | | | | |
|---|---|---|---|---|---|---|---|
| Bradi3g54550.1 | up | core (33) | PF00684, PF01556, PF00226 | - | GO:0051082, GO:0031072 | Molecular chaperone Hsp40/DnaJ family protein | chaperone protein dnaJ, putative, expressed |
| Bradi3g57013.1 | | shell (1) | - | - | - | - | - |
| Bradi3g57525.1 | up | shell (22) | PF00582 | - | GO:0006950 | Adenine nucleotide alpha hydrolases-like superfamily protein | universal stress protein domain containing protein, putative, expressed |
| Bradi3g58590.1 | | soft-core (32) | PF00011 | - | - | HSP20-like chaperones superfamily protein | heat shock 22 kDa protein, mitochondrial precursor, putative, expressed |
| Bradi3g58590.2 | | soft-core (32) | PF00011 | - | - | HSP20-like chaperones superfamily protein | heat shock 22 kDa protein, mitochondrial precursor, putative, expressed |
| Bradi3g59726.1 | up | core (33) | PF07728 | 3.6.1.3 | GO:0016887, GO:0005524 | P-loop containing nucleoside triphosphate hydrolases superfamily protein | AAA-type ATPase family protein, putative, expressed |
| Bradi3g59726.2 | up | core (33) | PF00004 | 3.6.1.3 | GO:0005524 | P-loop containing nucleoside triphosphate hydrolases superfamily protein | AAA-type ATPase family protein, putative, expressed |
| Bradi3g59726.3 | up | core (33) | PF07728 | 3.6.1.3 | GO:0016887, GO:0005524 | P-loop containing nucleoside triphosphate hydrolases superfamily protein | AAA-type ATPase family protein, putative, expressed |
| Bradi3g60090.1 | | core (33) | PF00226 | - | - | Chaperone DnaJ-domain superfamily protein | heat shock protein DnaJ, putative, expressed |
| Bradi3g60100.1 | up | soft-core (31) | PF00011 | - | - | HSP20-like chaperones superfamily protein | hsp20/alpha crystallin family protein, putative, expressed |
| Bradi4g05210.1 | up | core (33) | PF00582 | - | GO:0006950 | Adenine nucleotide alpha hydrolases-like superfamily protein | universal stress protein domain containing protein, putative, expressed |
| Bradi4g06030.1 | up | core (33) | PF07883 | - | - | endoplasmic reticulum auxin binding protein 1 | auxin-binding protein 4 precursor, putative, expressed |
| Bradi4g06370.1 | up | core (33) | PF02518, PF00183 | - | GO:0051082, GO:0006950, | HEAT SHOCK PROTEIN 89.1 | heat shock protein, putative, expressed |

| | | | | | | | | |
|---|---|---|---|---|---|---|---|---|
| Bradi4g06370.2 | up | core (33) | | PF02518, PF00183 | - | GO:0006457, GO:0005524 GO:0051082, GO:0006950, GO:0006457, GO:0005524 | HEAT SHOCK PROTEIN 89.1 | heat shock protein, putative, expressed |
| Bradi4g07520.1 | up | soft-core (31) | | PF02893 | - | - | GRAM domain family protein | GRAM domain containing protein, expressed |
| Bradi4g08550.2 | up | core (33) | | PF01370 | - | GO:0050662, GO:0003824 | NAD(P)-binding Rossmann-fold superfamily protein | NAD dependent epimerase/dehydratase family protein, putative, expressed |
| Bradi4g08790.1 | | core (33) | | PF13646 | - | - | protein phosphatase 2A subunit A2 | HEAT repeat family protein, putative, expressed |
| Bradi4g09120.1 | | core (33) | | PF00004 | - | GO:0005524 | rubisco activase | AAA-type ATPase family protein, putative, expressed |
| Bradi4g09120.2 | | core (33) | x | PF00004 | - | GO:0005524 | rubisco activase | AAA-type ATPase family protein, putative, expressed |
| Bradi4g17200.1 | up | core (33) | | - | - | - | - | expressed protein |
| Bradi4g17336.1 | up | shell (5) | | - | - | - | - | - |
| Bradi4g20537.1 | up | core (33) | | PF01256 | 4.2.1.93 | GO:0052855 | pfkB-like carbohydrate kinase family protein | expressed protein |
| Bradi4g20537.2 | up | core (33) | | PF01256 | 4.2.1.93 | GO:0052855 | pfkB-like carbohydrate kinase family protein | expressed protein |
| Bradi4g20537.3 | up | core (33) | | PF01256 | 4.2.1.93 | GO:0052855 | pfkB-like carbohydrate kinase family protein | expressed protein |
| Bradi4g20537.4 | up | core (33) | | PF01256 | 4.2.1.93 | GO:0052855 | pfkB-like carbohydrate kinase family protein | expressed protein |
| Bradi4g25430.4 | up | core (33) | | PF06058 | - | - | decapping 1 | mRNA-decapping enzyme, putative, expressed |
| Bradi4g29506.2 | | shell (23) | | - | - | - | - | - |
| Bradi4g31060.2 | | soft-core (32) | | PF02517 | 2.3.1.84 | GO:0016020 | alpha/beta-Hydrolases superfamily protein | CAAX amino terminal protease family protein, putative, expressed |

| | | | | | | | | |
|---|---|---|---|---|---|---|---|---|
| Bradi4g31070.1 | | core (33) | | PF07082 | - | - | Protein of unknown function (DUF1350) | expressed protein |
| Bradi4g31070.2 | | core (33) | | PF07082 | - | - | Protein of unknown function (DUF1350) | expressed protein |
| Bradi4g32941.1 | up | core (33) | x | PF02518, PF00183 | - | GO:0051082, GO:0006950, GO:0006457, GO:0005524 | Chaperone protein htpG family protein | heat shock protein, putative, expressed |
| Bradi4g32941.2 | up | core (33) | x | PF02518, PF00183 | - | GO:0051082, GO:0006950, GO:0006457, GO:0005524 | Chaperone protein htpG family protein | heat shock protein, putative, expressed |
| Bradi4g34056.1 | up | shell (29) | | - | - | - | - | expressed protein |
| Bradi4g34980.1 | | soft-core (32) | | PF10349 | - | - | - | arabinogalactan protein, putative, expressed |
| Bradi4g35170.2 | | core (33) | | - | - | - | - | expressed protein |
| Bradi4g35790.1 | up | core (33) | | PF01073 | 5.1.3.2 | GO:0055114, GO:0016616, GO:0006694, GO:0003854 | UDP-D-glucose/UDP-D-galactose 4-epimerase 1 | UDP-glucose 4-epimerase, putative, expressed |
| Bradi4g36690.1 | | core (33) | x | - | - | - | - | expressed protein |
| Bradi4g37770.1 | | core (33) | | PF00175, PF00258, PF00667 | 1.6.2.4 | GO:0055114, GO:0016491, GO:0010181 | P450 reductase 2 | NADPH reductase, putative, expressed |
| Bradi4g40197.1 | up | core (33) | | PF13414 | - | - | Tetratricopeptide repeat (TPR)-like superfamily protein | tetratricopeptide repeat protein 1, putative, expressed |
| Bradi4g42710.1 | up | core (33) | | - | - | - | - | expressed protein |
| Bradi4g43760.1 | up | core (33) | | PF04117 | - | GO:0016021 | Peroxisomal membrane 22 kDa (Mpv17/PMP22) family protein | Mpv17 / PMP22 family domain containing protein, expressed |
| Bradi5g02037.1 | | core (33) | x | PF02518, PF00183 | - | GO:0051082, GO:0006950, GO:0006457, GO:0005524 | heat shock protein 90.1 | heat shock protein, putative, expressed |

| | | | | | | | |
|---|---|---|---|---|---|---|---|
| Bradi5g06390.1 | up | core (33) | PF13414, PF00254 | 5.2.1.8 | GO:0006457 | FKBP-type peptidyl-prolyl cis-trans isomerase family protein | peptidyl-prolyl isomerase, putative, expressed |
| Bradi5g10250.1 | | core (33) | PF01025 | - | GO:0051087, GO:0042803, GO:0006457, GO:0000774 | Co-chaperone GrpE family protein | co-chaperone GrpE protein, putative, expressed |
| Bradi5g10450.1 | up | core (33) | PF06884 | - | - | Protein of unknown function (DUF1264) | DUF1264 domain containing protein, putative, expressed |
| Bradi5g10460.1 | up | soft-core (31) | PF13664 | - | - | late embryogenesis abundant domain-containing protein / LEA domain-containing protein | expressed protein |
| Bradi5g11630.1 | | soft-core (32) | PF00248 | - | - | NAD(P)-linked oxidoreductase superfamily protein | oxidoreductase, aldo/keto reductase family protein, putative, expressed |
| Bradi5g11980.2 | down | core (33) | PF00481 | 3.1.3.16 | GO:0003824, GO:0006470, GO:0004722 | Protein phosphatase 2C family protein | protein phosphatase 2C, putative, expressed |
| Bradi5g13930.1 | up | core (33) | PF02681 | - | - | Acid phosphatase/vanadium-dependent haloperoxidase-related protein | Divergent PAP2 family domain containing protein, expressed |
| Bradi5g13930.3 | up | core (33) | PF02681 | - | - | Acid phosphatase/vanadium-dependent haloperoxidase-related protein | Divergent PAP2 family domain containing protein, expressed |
| Bradi5g13930.4 | up | core (33) | PF02681 | - | - | Acid phosphatase/vanadium-dependent haloperoxidase-related protein | Divergent PAP2 family domain containing protein, expressed |
| Bradi5g14550.1 | up | core (33) | PF13893, PF14237, PF00076 | - | GO:0003676 | RNA binding (RRM/RBD/RNP motifs) family protein | RNA recognition motif containing protein, putative, expressed |
| Bradi5g15300.2 | | core (33) | PF00168, PF01764 | 3.1.1.3 | GO:0005515, GO:0006629 | triglyceride lipases;triglyceride lipases | lipase class 3 family protein, putative, expressed |

| | | | | | | | |
|---|---|---|---|---|---|---|---|
| Bradi5g15300.3 | | core (33) | PF00168, PF01764 | 3.1.1.3 | GO:0005515, GO:0006629 | triglyceride lipases;triglyceride lipases | lipase class 3 family protein, putative, expressed |
| Bradi5g15835.1 | up | core (33) | - | - | - | - | expressed protein |
| Bradi5g15970.1 | | core (33) | PF00230 | - | GO:0016020, GO:0006810, GO:0005215 | plasma membrane intrinsic protein 2 | aquaporin protein, putative, expressed |
| Bradi5g18290.1 | | core (33) | PF06244 | - | - | - | coiled-coil domain-containing protein 124, putative, expressed |
| Bradi5g20410.1 | | core (33) | PF03803 | - | - | scramblase-related | scramblase, putative, expressed |
| Bradi5g22240.1 | down | core (33) | PF13460 | - | - | NAD(P)-binding Rossmann-fold superfamily protein | NAD dependent epimerase/dehydratase family protein, putative, expressed |
| Bradi5g23544.1 | up | shell (27) | - | - | - | - | - |
| Bradi5g23910.1 | up | core (33) | PF00892 | - | GO:0016021, GO:0016020 | EamA-like transporter family | thiamine-repressible mitochondrial transport protein THI74, putative, expressed |
| Bradi5g25450.1 | up | core (33) | PF12937, PF00022 | - | GO:0005515 | actin-related protein 8 | actin-6, putative, expressed |
| Bradi5g25450.2 | up | core (33) | PF12937, PF00022 | - | GO:0005515 | actin-related protein 8 | actin-6, putative, expressed |

### Supplementary Figures

**Figure S1.** Clustering dissimilarity dendrograms based on the topological overlap, together with assigned module colors from weighted gene co-expression networks (WGCN) of the Drought **(a)** and Water **(b)** networks, and hub Drought **(c)** and Water **(d)** networks (excluding gray or "zero" module) represented for the hub nodes and their connections using the R package wgcna2igraph with the parameters kME.threshold = 0.8, adjacency.threshold = 0.1, adj.power = 6. Modules are color-coded and their corresponding numerical codes are shown in Table 1.

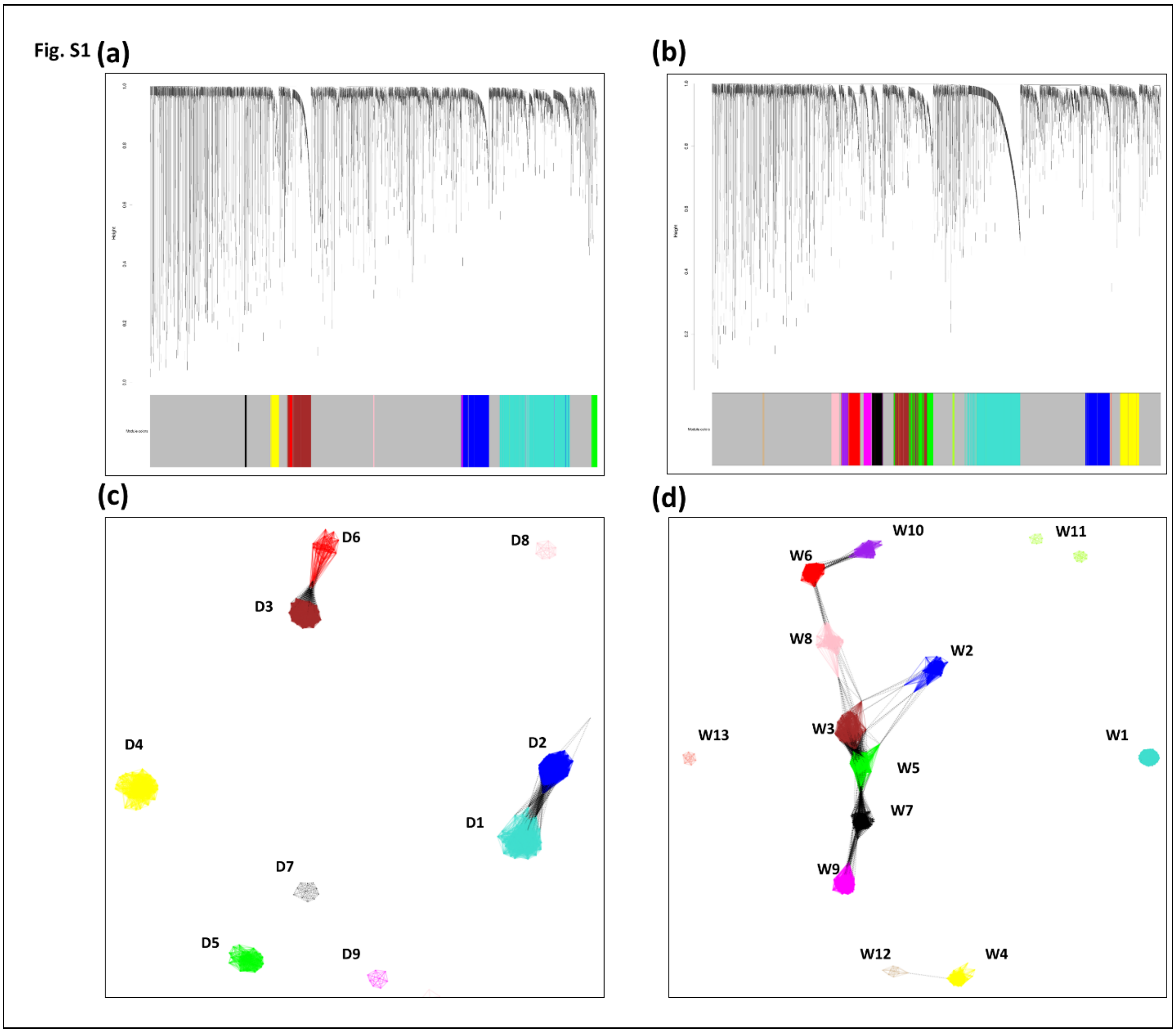

**Figure S2.** Dendrograms of the module eigengenes (ME) for each Drought (D) (**a**) and Water (W) (**b**) networks.

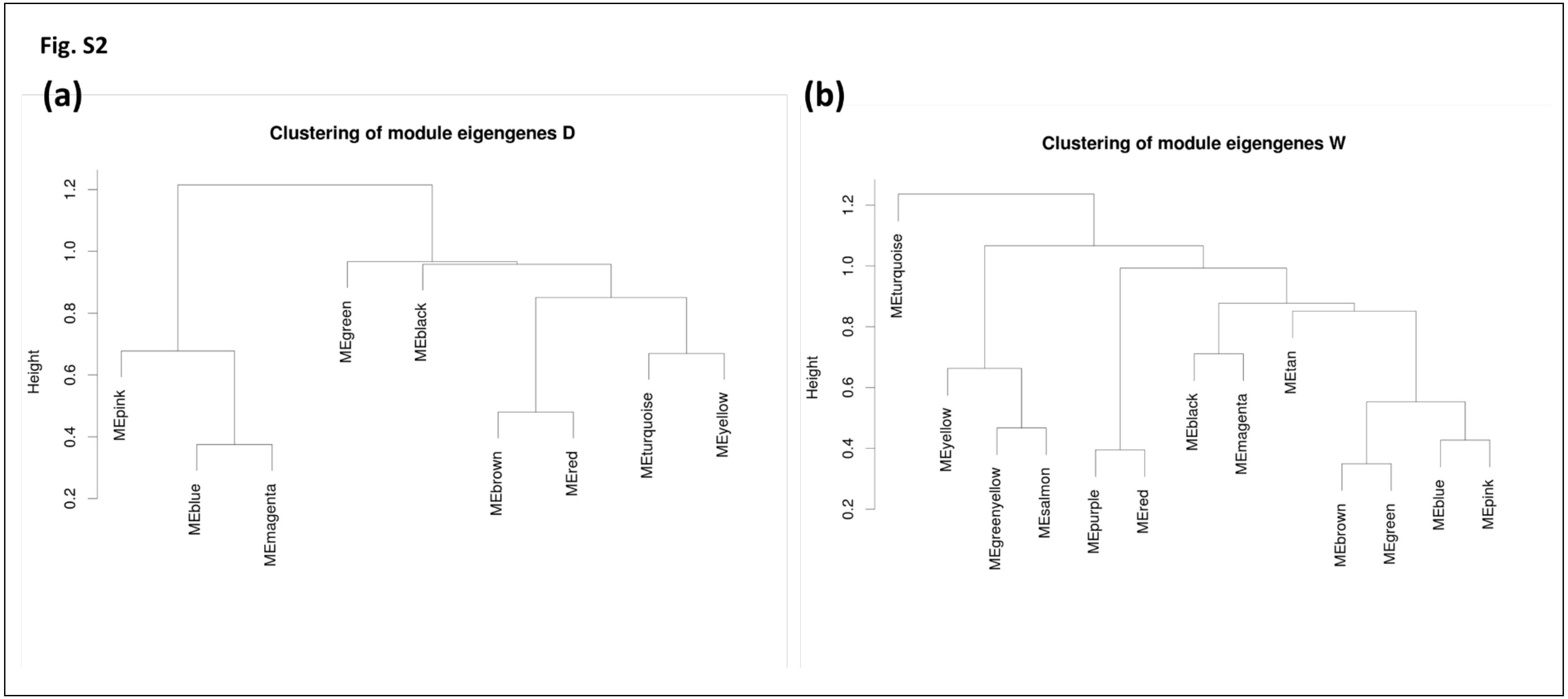

**Figure S3.** Correlations between module membership (MM) and intra-modular connectivity ($k_{IM}$) in Drought (**a**) and Water (**b**) networks.

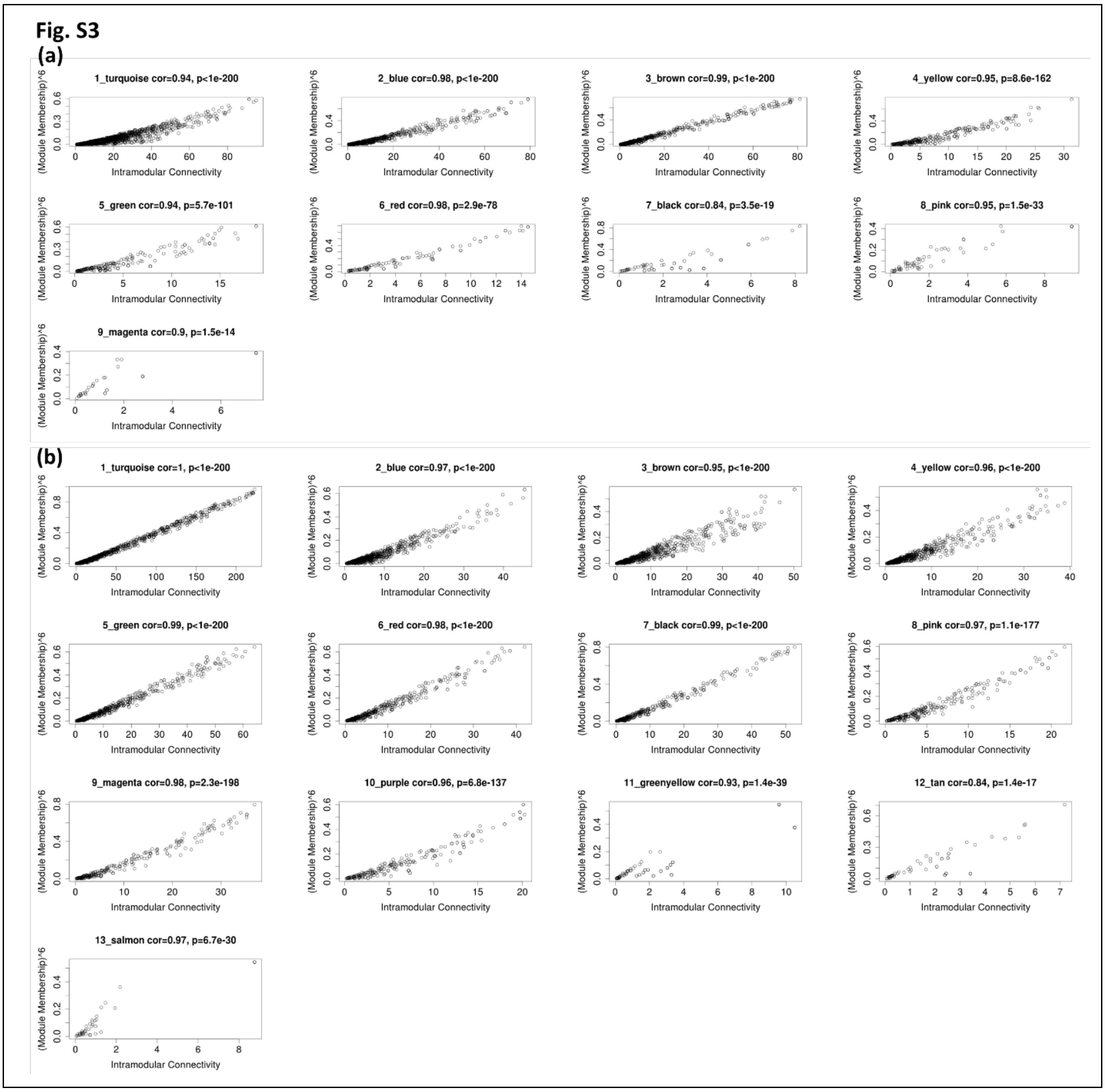

**Figure S4.** Boxplots of topological features (Connectivity, Clustering-Coefficient, MAR (Maximum Adjacency Ratio) and Scaled-Connectivity) of Drought (red) and Water (blue) networks (excluding gray or zero module). Significance is noted by Wilcoxon test (N.S. not-significant; ***P ≤ 0.001).

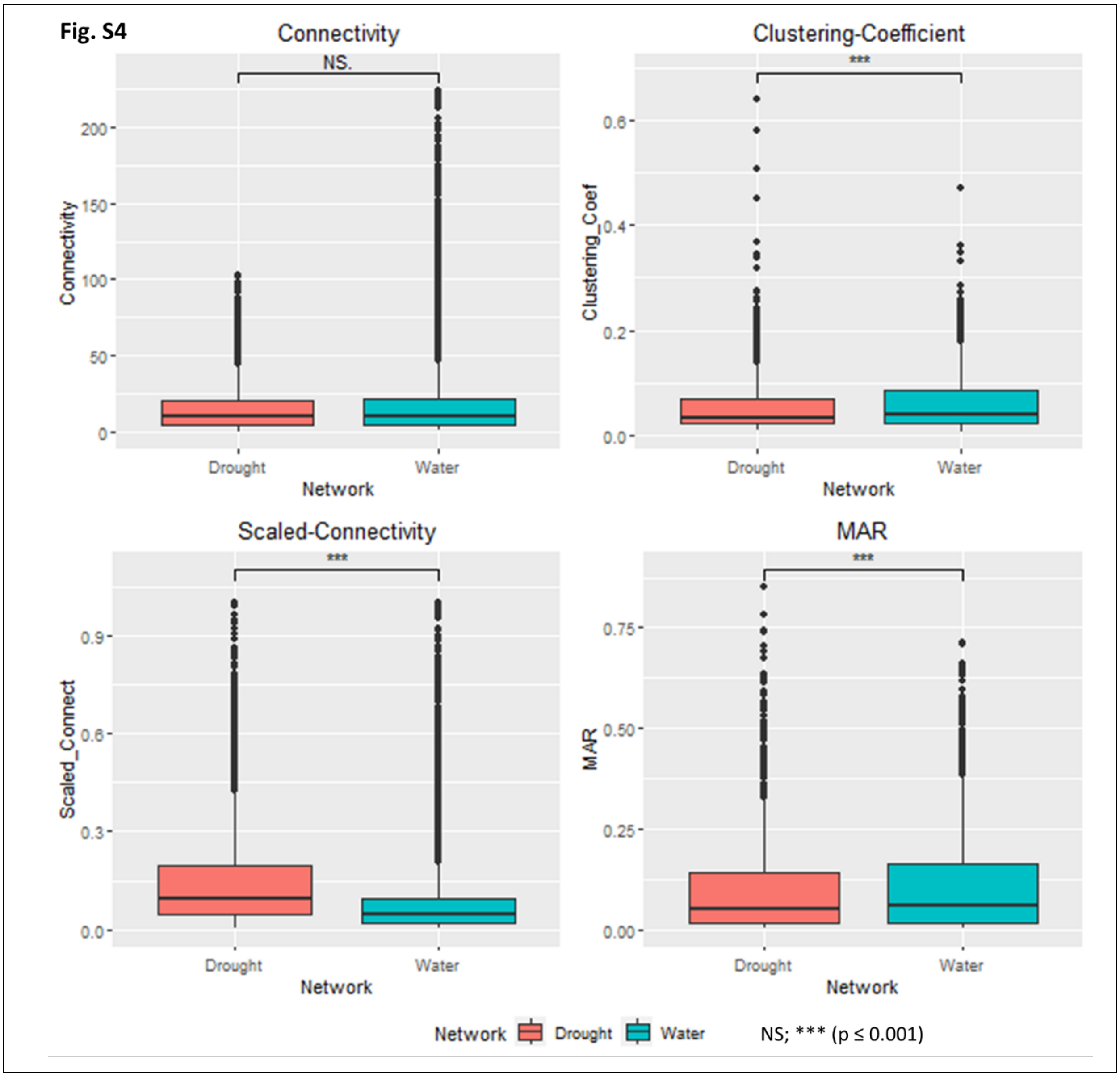

**Figure S5**. Discovery DNA motifs/Cis-regulatory elements analysis of co-expressed genes of the Drought (**a**) and Water (**b**) modules, adding 50 negative controls of equal size showing the significance of target module compared to random modules. GO enrichment, peaks-oligo, peaks-dyad and footprintDB analyses were conducted. The reports can be browsed at http://rsat.eead.csic.es/plants/data/modulesD and http://rsat.eead.csic.es/plants/data/modulesW/, where black bars correspond to co-expressed regulons and grey bars to negative controls.

### Fig. S5

### (a)

| name | GO enrichment | peaks-oligo | peaks-dyad | footprintDB |
|---|---|---|---|---|
| cluster D1 sequences: 1976 | GO:0006807 nitrogen compound metabolic process | cagccca\|tgggctg | ccan{1}ccc\|gggn{1}tgg | |
| cluster D2 sequences: 750 | GO:0044699 single-organism process | taaaa\|tttta | cacn{2}ccc\|gggn{2}gtg | |
| cluster D3 sequences: 627 | GO:0044699 single-organism process | gtcaa\|ttgac | agtcaa\|ttgact | |
| cluster D4 sequences: 259 | GO:0044238 primary metabolic process | accgcgt\|acgcggt | acgcgc\|gcgcgt | |
| cluster D5 sequences: 169 | GO:0006457 protein folding | tccaga\|tctgga | gaan{7}gaa\|ttcn{7}ttc | |
| cluster D6 sequences: 95 | GO:0044710 single-organism metabolic process | cacac\|gtgtg | acan{3}cac\|gtgn{3}tgt | |
| cluster D7 sequences: 48 | GO:0050789 regulation of biological process | ataaga\|tcttat | attn{6}taa\|ttan{6}aat | |
| cluster D8 sequences: 53 | GO:0005840 ribosome | atagaa\|ttctat | atagaa\|ttctat | |
| cluster D9 sequences: 27 | GO:1901566 organonitrogen compound biosynthetic process | acgtg\|cacgt | atan{11}tga\|tcan{11}tat | |

### (b)

| name | GO enrichment | peaks-oligo | peaks-dyad | footprintDB |
|---|---|---|---|---|
| cluster W1 sequences: 1437 | GO:0044763 single-organism cellular process | aaaatt\|aatttt | aaan{17}aaa\|tttn{17}ttt | |
| cluster W2 sequences: 728 | GO:0006412 translation | ccagccc\|gggctgg | cccn{20}ccc\|gggn{20}ggg | |
| cluster W3 sequences: 697 | GO:0044699 single-organism process | taaaa\|tttta | aan{1}tta\|taan{1}ttt | |
| cluster W4 sequences: 582 | GO:0044237 cellular metabolic process | aaaaa\|ttttt | taan{5}aaa\|tttn{5}tta | |
| cluster W5 sequences: 509 | GO:0044699 single-organism process | agtcaa\|ttgact | agtcaa\|ttgact | |
| cluster W6 sequences: 358 | GO:0044699 single-organism process | accaacc\|ggttggt | accaac\|gttggt | |
| cluster W7 sequences: 314 | GO:1901564 organonitrogen compound metabolic process | gtcaa\|ttgac | gtcaaa\|tttgac | |
| cluster W8 sequences: 245 | GO:0044699 single-organism process | cgcac\|gtgcg | cacacg\|cgtgtg | |
| cluster W9 sequences: 239 | GO:0050794 regulation of cellular process | accgcgt\|acgcggt | acacgc\|gcgtgt | |
| cluster W10 sequences: 185 | GO:0015979 photosynthesis | ccaca\|tgtgg | cagcca\|tggctg | |
| cluster W11 sequences: 55 | GO:0065007 biological regulation | aaagg\|ccttt | aaan{9}aac\|gttn{9}ttt | |
| cluster W12 sequences: 51 | GO:0050794 regulation of cellular process | aaata\|tattt | atan{7}ata\|tatn{7}tat | |
| cluster W13 sequences: 36 | GO:0034645 cellular macromolecule biosynthetic process | agaat\|attct | aatn{5}ttc\|gaan{5}att | |

**Supplementary Files:**

**File S1.** Detailed statistics of the enrichment analysis according to the statistically significant GO biological process detected for the genes (all, core, soft-core and shell genes) clustered in the Drought (D) and Water (W) modules applying the statistical overrepresentation test of Panther (http://pantherdb.org/) tool and sorted by the lowest FDR. Every tab corresponds to the all, core, soft-core or shell genes results for each module (e.g. D_1: enrichment results for all genes of the Drought module 1; D_1_core: enrichment results for the core genes of the Drought module 1; …). There are not tabs to the "No statistically significant results". The headers of the table correspond to Ref-list (number of genes of the reference *Brachypodium distachyon* list (34,230 genes) from Ensembl source that map to this particular annotation data category); Client Text Box Input (number of genes in the query uploaded list that map to this particular annotation data category); expected (number of genes you would expect in your list for this category, based on the reference list); +/- (A plus/minus sign indicates over/under-representation of this category in your experiment (you observed more/fewer genes than expected based on the reference list for this category)); fold Enrichment (genes observed in the uploaded list with respect to the expected genes (number of genes in the query list divided by the expected number of genes). If it is greater than 1, it indicates that the category is overrepresented in your experiment. Conversely, the category is underrepresented if it is less than 1); raw p-value (determined by Fisher's exact test. This is the probability that the number of genes you observed in this category occurred by chance (randomly), as determined by your reference list); False Discovery Rate (FDR) (by default a critical value of 0.05 is used to filter results, therefore all results shown are valid for an overall FDR<0.05 even if the FDR for an individual comparison is greater than that value).

**File S2.** Expression of filtered and normalized (by Sleuth package) TPM (transcripts per million) of the 4,941 differentially expressed (DE) isoforms (3,489 DE genes) under the water (W) and drought (D) conditions. The averages TPM per treatment for each isoform and their differences between Drought and Water treatments were computed.

**File S3.** Summary annotation of the 4,941 differentially expressed (DE) isoforms (3,489 DE genes) under drought (D) and water (W) conditions. Isoform (*B. distachyon*_314_v3.1 (Bd21 accession) reference transcriptome); regulation under drought condition compared to water condition; occupancy (core (33 accessions), soft-core (31-32 accessions) and shell (≤30 accessions)); matched Drought modules and matched Water modules (numeric identification of the module where the DE isoforms were assigned); DE hub node (DE gene detected such as hub gene in the Drought (D) or Water (D) modules); Pfam, KEGG/ec, GO annotations; arabi-defline and rice-defline (annotation in Arabidopsis and rice).
